# Host pathways associated with human bacterial infections extend to commensal *Wolbachia-Drosophila* endosymbiosis

**DOI:** 10.1101/2022.07.02.498523

**Authors:** Zinat Sharmin, Hani Samarah, Rafael Aldaya Bourricaudy, Laura Renee Serbus

## Abstract

*Wolbachia* bacteria are among the most successful endosymbionts in nature, carried by half of all insect species. Unlike human bacterial pathogens that kill host cells and tissues, *Wolbachia* endosymbionts are generally carried by insects with little adverse effect. The striking difference in outcome raises a basic question of what aspects of infection mechanisms are conserved across systems. In this study, 37 small molecule inhibitors were used to test whether 14 candidate host processes that affect the abundance of other intracellular bacteria also affect *Wolbachia*. Compounds that significantly affected the absolute abundance of the *Wolbachia surface protein (wsp)* gene in *D. melanogaster* were retested in *D. simulans* flies. 5 compounds that consistently increased *wsp* abundance in both systems were associated with the Imd pathway, Calcium signaling, Ras/mTOR signaling, and the Wnt pathway. By contrast, the only compound to suppress *wsp* abundance was a Ubiquitin-proteasome pathway inhibitor. The implicated host processes were retested for impact on *Wolbachia* using constitutive and inducible RNAi expression systems in *D. melanogaster*. These tests corroborated a function for the host *target of rapamycin* (*tor*) and *armadillo* (*arm*) genes in affecting bodywide *wsp* abundance. Prior studies have reported that Ras/mTOR and Wnt pathways interact with ATG6 (Beclin-1), representing a possible convergence point for signaling impacts on *Wolbachia*. ATG6 disruption tests, driven by inducible RNAi expression, also elevated *wsp* abundance. This work suggests that combined effects of the Wnt pathway, Ras/mTOR signaling, and autophagy normally support *Wolbachia* containment, moderating the *Wolbachia-*host endosymbiosis.

**IMPORTANCE:** Disease-related microbes have been intensively studied as a model for infection. An intrinsic complication of such studies is bacterial induction of cell stress and cell death. To expand our understanding of cellular infection mechanisms, we studied a bacterial endosymbiont of insects, called *Wolbachia*, that does not kill the cells it infects. We asked whether cellular processes involved in pathogen infection are also associated with *Wolbachia* infections. Chemical and genetic tests were used to investigate cellular effects on *Wolbachia* abundance within fruit flies. We identified a subset of cellular processes with robust, repeatable effects on *Wolbachia* infection: the Wnt pathway and the Ras/mTor pathway. The results also suggest that multiple cellular pathways act together, which collectively suppresses *Wolbachia* abundance in vivo. Active host containment may explain in part why *Wolbachia* is mostly regarded as a neutral endosymbiont, and not as a pathogen.

## INTRODUCTION

Resident intracellular microbes, also referred to as “endosymbionts,” are widespread in nature. Endosymbiotic microbes are commonly thought of as “mutualists,” in which the interaction between host and microbe benefits both partners of the symbiosis. However, endosymbionts can also exhibit relatively inert (commensal) or detrimental (parasitic) interactions with a host organism. Evidence suggests that some commensal/mutualistic microbes are descendants of formerly parasitic ancestors (Sachs et al., 2014). Other endosymbionts have been found to exhibit context-dependent plasticity in their symbiotic interactions, such as for *Salmonella*, which is carried innocuously by chickens, but causes severe infection in humans. To account for this diversity, intracellular bacteria are now described in terms of a symbiotic spectrum, ranging from mutualism to parasitism (Lewis, 1985).

Efforts focused on biomedical forms of parasitism, including human bacterial disease, have been a major focus of endosymbiosis research to date. Cellular studies of infection face the generalized challenge that cell stress and lethality unavoidably coincide with experimental readouts, complicating data interpretation. Far less is known about the molecular mechanisms of infection used by commensal and mutualistic bacteria. Due to the extent of shared attributes across human and insect systems (Greenspan & Kreitman, 2008; Pandey & Nichols, 2011), bacterial endosymbionts can be reasonably expected to encounter similar host cellular resources as well as constraints upon infection. Fundamental questions remain regarding what host cellular processes have consensus involvement in endosymbiotic infection mechanisms. It also remains unclear what functional attributes distinguish commensal and beneficial infections from virulent scenarios.

For any endosymbiont, high prevalence in nature is the mark of success. *Wolbachia* bacteria are naturally widespread bacterial endosymbionts, carried by lineages of mites, crustaceans, and nematodes as well as 50% of all insect species. *Wolbachia* are often described as reproductive parasites because the infection affects host reproduction in certain species (Werren et al., 2008). In some instances, *Wolbachia* serve as mutualists, by sustaining host viability and reproduction (Landmann et al., 2011; Pannebakker et al., 2007; Taylor et al., 2005), as well as by repelling viral infections harmful to the host (Cogni et al., 2021; Hedges et al., 2008; Teixeira et al., 2008). For this to be possible, *Wolbachia* bacteria must be able to colonize the host without inducing generalized host cell lethality. However, most *Wolbachia* infections may be best described as commensal, in the absence of clear benefits or negative effects on the host. The *Wolbachia-*host symbiosis thus provides a new and valuable perspective for investigating, defining and understanding the cellular basis of infection.

To date, studies of *Wolbachia* infection have shared a goal of understanding how *Wolbachia* amount (titer) is specified within tissue culture cells, endogenous host cells and tissues, and in whole organisms overall. *Wolbachia* titer has been assessed as a function of various host/strain combinations as well as in response to host age, crowding, temperature, diet, genetic background, microbiota and chemical exposure (Hoffmann et al., 1998; López-Madrigal & Duarte, 2019; Unckless et al., 2009; Veneti et al., 2004; Voronin et al., 2012; Wiwatanaratanabutr & Kittayapong, 2009)

A patchwork of cytological and qPCR-based methods have been used across assessments thus far. A unifying premise in many of such studies is that communication between the host cell and *Wolbachia* is an important contributor to *Wolbachia* titer outcomes. The field is now in a strong position to investigate how *Wolbachia-*host interactions can broadly inform mechanisms of infection.

The cellular and molecular basis of bacterial titer control has been well-studied over decades for disease related microbes. For example, plant cells use the ubiquitin-proteasome system as part of an antibacterial resistance mechanism against *Pseudomonas* (Üstün et al., 2016). By contrast, host calcium and Wnt signaling reportedly facilitate growth of *Coxiella, Legionella, Brucella, Rickettsia, Ehrlichia, Chlamydia*, and *Ehrlichia* (Czyż et al., 2014; Rikihisa et al., 1995; Kessler et al., 2012; Luo et al., 2016). Epidermal Growth Factor Receptor (EGFR) signaling has been shown to promote host cell invasion by pathogens like *Salmonella* and *Neisseria* (Galán et al., 1992; Slanina et al., 2014). Other processes, such as the mTOR/autophagy pathway, exert differential density effects depending on the bacterial strain. For example, mTOR signaling disruptions reduce intracellular loads for *Ehrlichia* (Luo et al., 2017), *Chlamydia, Listeria* (Derré et al., 2007), and *Salmonella* (Birmingham et al., 2006), but increase titers for *Anaplasma* and *Rickettsia* (Bechelli et al., 2018; Niu et al., 2008). Recurrent titer effects for host cellular processes on unrelated bacterial taxa invoke the possibility of generalized infection roles for host cellular pathways.

In this study, we asked how *Wolbachia* titer responds to host cell processes that are otherwise associated with human bacterial infections. Functional tests were performed on 14 candidate host pathways and processes, using complementary chemical and genetic tools. Whole-body *Wolbachia* abundance was assessed in all cases by real-time qPCR, to determine absolute counts of the *Wolbachia surface protein* (*wsp*) gene. This work yielded a subset of host functions for further pursuit, with implications for the basis of commensalism as described below.

## MATERIALS AND METHODS

### *Drosophila* stocks and maintenance

Two fly strains were used in this study. Preliminary screening was performed using *Drosophila melanogaster* of the genotype *w*; *Sp/Cyo*; *Sb/TM6B* carrying the endogenous *w*Mel *Wolbachia* strain (Christensen et al., 2016; Serbus & Sullivan, 2007). *Drosophila simulans* infected with the endogenous *Wolbachia riverside* (*w*Ri) strain was used for further analyses (Hoffmann et al., 1990; Serbus & Sullivan, 2007). GAL4 lines sourced from the Bloomington Drosophila Stock Center (BDSC) were also used to drive RNAi expression. Constitutive body-wide expression was driven by the Actin5C-GAL4 driver *w; P{Act5C-GAL4-w}E1/Cyo* (BDSC# 25374) and the daughterless-GAL4 driver *w; P{w+, GMR12B08-GAL4}attP2* (BDSC# 48489). Drug-inducible gene expression was driven by the GeneSwitch-GAL4 driver *yw {hs-FLP}; {w+, UAS-GFP}; {w+, Act5C-GS-GAL4}/TM6B, Tb* (BDSC #9431).

Flies are usually maintained in plastic bottles/vials containing standard fly food media. The recipe is derived from Bloomington stock center as described previously (Christensen et al., 2016). The flies were raised in an Invictus *Drosophila* incubator at 25°C, under standard 12/12h light-dark cycle. For the experiments, “0-day old” flies were collected and kept on standard fly food medium for 2-days. 2-days old flies were then used for drug treatments in vials or within a plate assay format as described previously (Christensen et al., 2019). Only female flies were used for the plate-based screening experiments, to reduce possible variation in population behavior per well.

### Chemical food preparation

Two or more chemicals were used to alter the functionality of each of the candidate host processes, comprising a total of 37 chemicals. The chemicals were purchased from different vendors (Table S1). All the drugs were dissolved in DMSO. Most stock solutions, including Rifampicin, were prepared in advance as 10 mM solutions, aliquoted and stored at -20°C. Rapamycin was ordered as a 5mM solution in DMSO, also stored at -20°C (Table S1). Light-sensitive drugs were stored in the dark.

Immediately before use, chemical stocks were thawed and diluted 100X into fly food that had been re-melted, then cooled. As a carrier control, equivalent amounts of DMSO were added to re-melted and cooled fly food as well, prepared in parallel with the chemical treatment vials. In all cases, the final concentration of DMSO in food was capped at 1%. For the chemical screen, a minimum of 10mL drug food was prepared per condition, to be further dispensed in approximately 1mL amounts per treatment well. For *GS-GAL4* induction that was carried out in vials, food containing control DMSO and DMSO-solubilized mifepristone was prepared in 200mL volumes, to be dispensed into vials as 5mL amounts. After pouring, plates and vials were cooled and solidified in the fume hood, with foil wrappings used to protect light-sensitive compounds. Treatment vials were stored in Ziplock bags at 4 degrees C as needed.

Chemical treatments were administered to 2-day old flies. For the chemical screen,10 female flies were transferred to each treatment well. After 3-days of feeding, pools of 5 female flies were removed from each well and processed as a group for *wsp* quantification using real-time qPCR. For the vial-based experiments using DMSO and mifepristone, flies were incubated on treatment food as groups of 15 females and 5 males. Flies were transferred to new treatment vials every 3 days, using DMSO and mifepristone vials from the 4C fridge that had been re-warmed. After 14 days of feeding were completed, pools of 5 female flies were removed from each vial and processed as a group for *wsp* quantification using real-time qPCR.

### Genetic manipulations

To put *Wolbachia* into all driver lines, GAL4 driver males were crossed to virgin females of the genotype *w*; *Sp/Cyo*; *Sb/TM6B*, carrying the *w*Mel Wolbachia strain (Christensen et al., 2016), hereafter referred to as DB wMel. This process eventually established *Wolbachia-*infected *Act5C-GAL4* and *da-GAL4* driver stocks, with the same genotypes as the originally uninfected lines. The final genotype for the *Wolbachia-*infected *Act5C-GS-GAL4* line was *yw {hs-FLP, w+}; {UAS-GFP, w+}/Cyo; {Act-GS-GAL4, w+}/TM6B*, missing the tubby marker on TM6B.

To generate RNAi expressing flies, *w*Mel-infected virgin females were selected from freshly emerging bottles of each driver stock. These females were crossed to males that carried a responder UAS (upstream activating sequence) adjacent to a ds-RNA coding sequence. The parent flies were removed from the vials after 3-4 days of mating. Emerging F1 flies were collected in daily cohorts and aged for 5 days. The F1s that carried both the GAL4 driver and the UAS responder were visually identified and separated within each cohort. Because this subset of flies is capable of driving expression of snap-back RNA, complementary to target genes listed in Table S2, they are hereafter referred to as “expressing” genotypes. Control siblings that contained either the GAL4 or UAS responder, but not both, are collectively referred to as “non-expressing” flies. A separate control set was also prepared in parallel with *Wolbachia-*infected virgin females from the driver stocks outcrossed to Oregon R (OreR) males. The RNAi-expressing group, the non-expressing controls, and OreR controls were processed in parallel for *wsp* quantification.

### DNA extraction and qPCR for whole body *Wolbachia* quantification

To quantify whole body *wsp* abundance, DNA was extracted from the female flies as previously described (Christensen et al., 2019). Absolute measurements of the *Wolbachia surface protein* gene from the extracted DNA samples were obtained using reference plasmid standards, specifically a PGEM-T vector carrying a 160 bp PCR-amplified fragment of the *(wsp)* gene (Christensen et al., 2016). Real-time PCR was carried out on a Bio-Rad CFX96 Connect Optics Module Real-Time System. Absolute copy numbers were obtained by comparing cycle threshold (Ct) values to standard curve generated from the plasmid standard. The primers used to target *wsp* gene were: Fwd 5’ CATTGGTGTTGGTGTTGGTG 3’ and Reverse 5’ ACCGAAATAACGAGCTCCAG 3’ primers, used at 5 μM (Christensen et al., 2016).

### Data display and analyses

Graphical displays showing “normalized” *wsp* counts as a scatter plot were created for display purposes only. To generate such graphs, median *wsp* counts for the DMSO controls per replicate were identified, then compared to the median *wsp* count of the entire dataset. A scaling factor was then identified and applied to each replicate, to normalize the median *wsp* value for the DMSO control and all associated experimental data. All other scatter plot graphs, as well as all supplemental tables, directly show the raw absolute count data. Statistical analyses were conducted on raw data within each experimental replicate for all experiments. Statistics appropriate to data normality and homogeneity were identified and applied as previously (Christensen et al., 2019). Power analysis was performed with alpha set at 0.05 using a MATLAB-based data sub-sampling program, designed by Dr. Philip K. Stoddard. This program has the advantage that analyses can be customized to the statistical test appropriate to each dataset (Christensen et al., 2019).

## RESULTS

### *Wolbachia surface protein (wsp)* responds to chemical disruption of the proteasome

To identify candidate host pathways and processes that affect whole-body *wsp* levels, a chemical feeding approach was used. This provides straightforward control over treatment conditions as well as the advantage of portability into genetically intractable model systems. We previously showed that Rifampicin significantly reduces *Wolbachia* titer within a 3-day assay time frame, but have not previously tested this on other non-antibiotic compounds. To validate parameters for chemical screening, test runs were first performed that target the ubiquitin-proteasome system (UPS). This pathway was selected due to the prominence of UPS impact on *Wolbachia* in tissue culture and germline cells (Grobler et al., 2018; White et al., 2017a).

To test the UPS for possible effects on whole-body *Wolbachia* titer, flies were exposed to the proteasome inhibitor bortezomib (Adams et al., 1999; Richardson et al., 2003). Experiments were carried out in a 24-well plate format as previously for consistency across technical replicates (Momtaz et al 2020). After 3 days exposure to 100 µM bortezomib, as used in previous fly and cell culture screens (Kim et al., 2008; Markstein et al., 2014; Serbus et al., 2012), *wsp* absolute counts were measured using the real-time qPCR (n= 6 wells per condition, 3 technical replicates each). The bortezomib-treated flies exhibited 49–52% of the median *wsp* abundance detected in DMSO control flies (*p* <0.001) (n= 18; 6 wells with 3 technical replicates each) (Fig. 1A, Table S3, Table S4). Power analysis further indicated that sampling was sufficient to detect a significant difference between DMSO and bortezomib conditions (β < 0.002 at n ≥ 4; total n = 18) (Fig. 1B).

**Figure 1.**
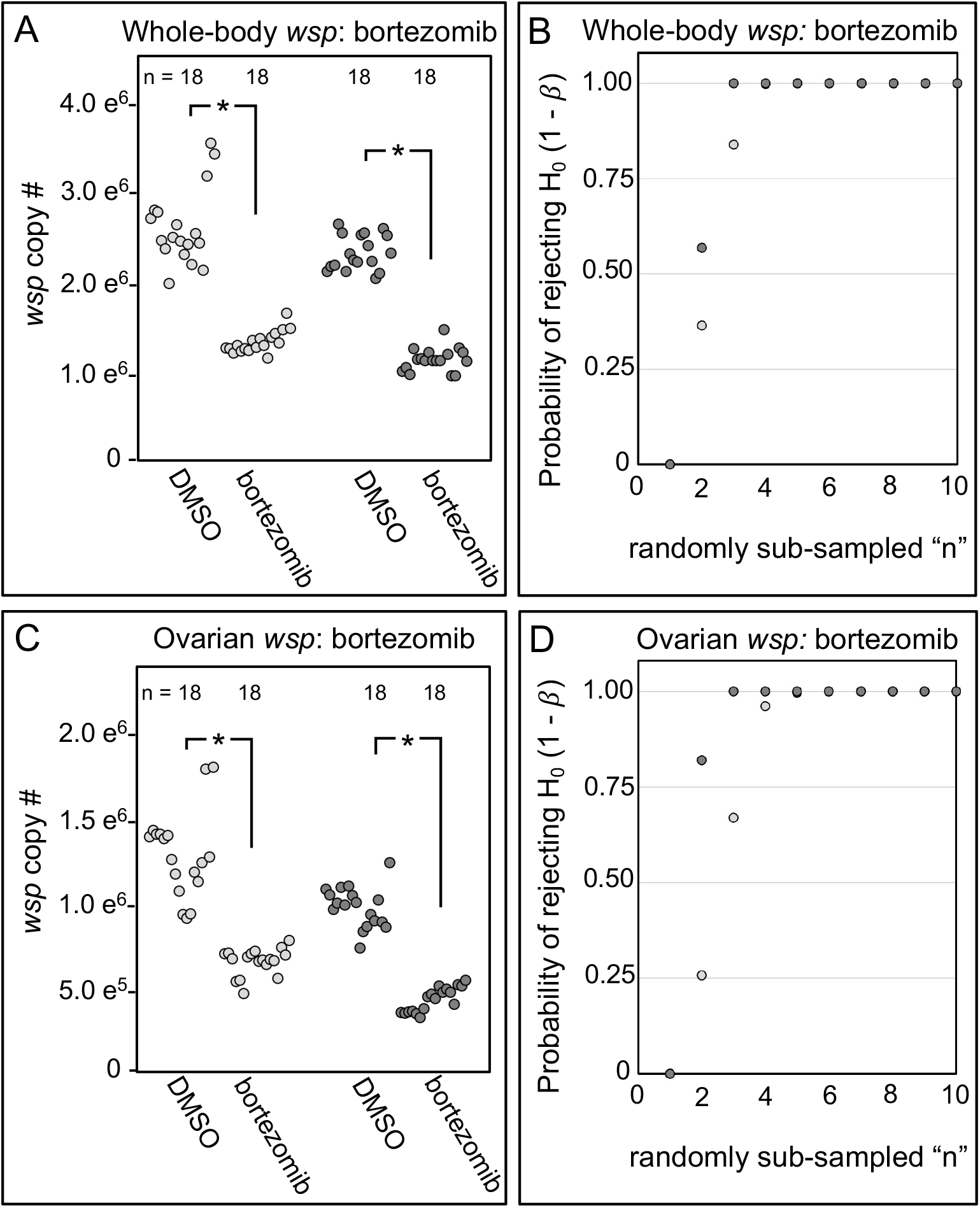
*wsp* abundance in whole body as indicated by real-time qPCR. Data show DMSO controls versus 100 μM bortezomib in DMSO. Panels show data from 2 independent biological replicates. A) Body-wide *wsp* abundance measurements. B) Power analysis, testing the likelihood of significance as a function of sample size in Panel A. C) Ovarian *wsp* abundance measurements. D) Power analysis, testing the likelihood of significance in Panel C. Significance was set at * p < 0.05.

To confirm penetrance of chemical treatment into the fly body, *wsp* levels were also tested in ovarian tissues of bortezomib-treated flies, prepared in parallel with the above. The ovaries of bortezomib-treated flies carried 44–53% of the median *wsp* abundance detected in DMSO control flies (*p* <0.001) (n= 18; 6 wells with 3 technical replicates each) (Fig. 3A, Table S5, Table S6). Power analysis also supports the significance of this result (β < 0.003 at n ≤ 5; total n = 18) (Fig. 1D). These data indicate that bortezomib has access to the body cavity of the insect, and ultimately reduces body-wide absolute abundance of *wsp*, consistent previously reported UPS effects on *Wolbachia* titer. As such, adult drug feedings can provide insight into host pathway effects on *Wolbachia* in general. These results also provide guidelines in reasonable sampling (now set at n = 6) for chemical tests of other candidate *Wolbachia*-interacting host pathways.

### Literature search for candidate host processes that broadly affect bacterial infection

Host effects on the density of intracellular bacteria have been studied for a number of bacterial pathogens. It is unclear whether host factors that influence pathogen density also affect colonization of eukaryotic cells by non-pathogenic bacteria. To identify potentially conserved host effects on intracellular bacteria titer, we searched the literature for host factors that affect intracellular abundance of commonly studied microbes. After investigating 52 species from 17 genera, we found 26 bacterial species for which involvement of host genes/pathways on density regulation has been discussed (Table S7). Many host processes have thus far been associated with a single bacterial class, which could reflect specificity of host involvement, or alternatively, heterogeneity in existing host-pathogen literature. Follow-up literature searches highlighted 14 host mechanisms that affect the intracellular abundance of multiple bacterial classes, prioritized for pursuit in the *Wolbachia-Drosophila* endosymbiosis model (Table S8) (Additional File S1).

To test candidate host processes for effects on whole-body *Wolbachia* abundance, candidate compounds were selected that are already known to target each process. 2 or more compounds were identified for testing each of the 14 classes of host targets (Table 1) (Table S1). Where possible, compounds with opposite effects on the process of interest were included, such as for the microtubule-depolymerizing drug, colchicine, and the microtubule-stabilizing drug, Taxol; and also for the phospholipase C (PLC) inhibitor, U73122, and the PLC activator, 3-m3mfbs. This culminated in selection of 37 total candidate compounds to test as a pilot chemical screen (Table S1).

**Table 1.**
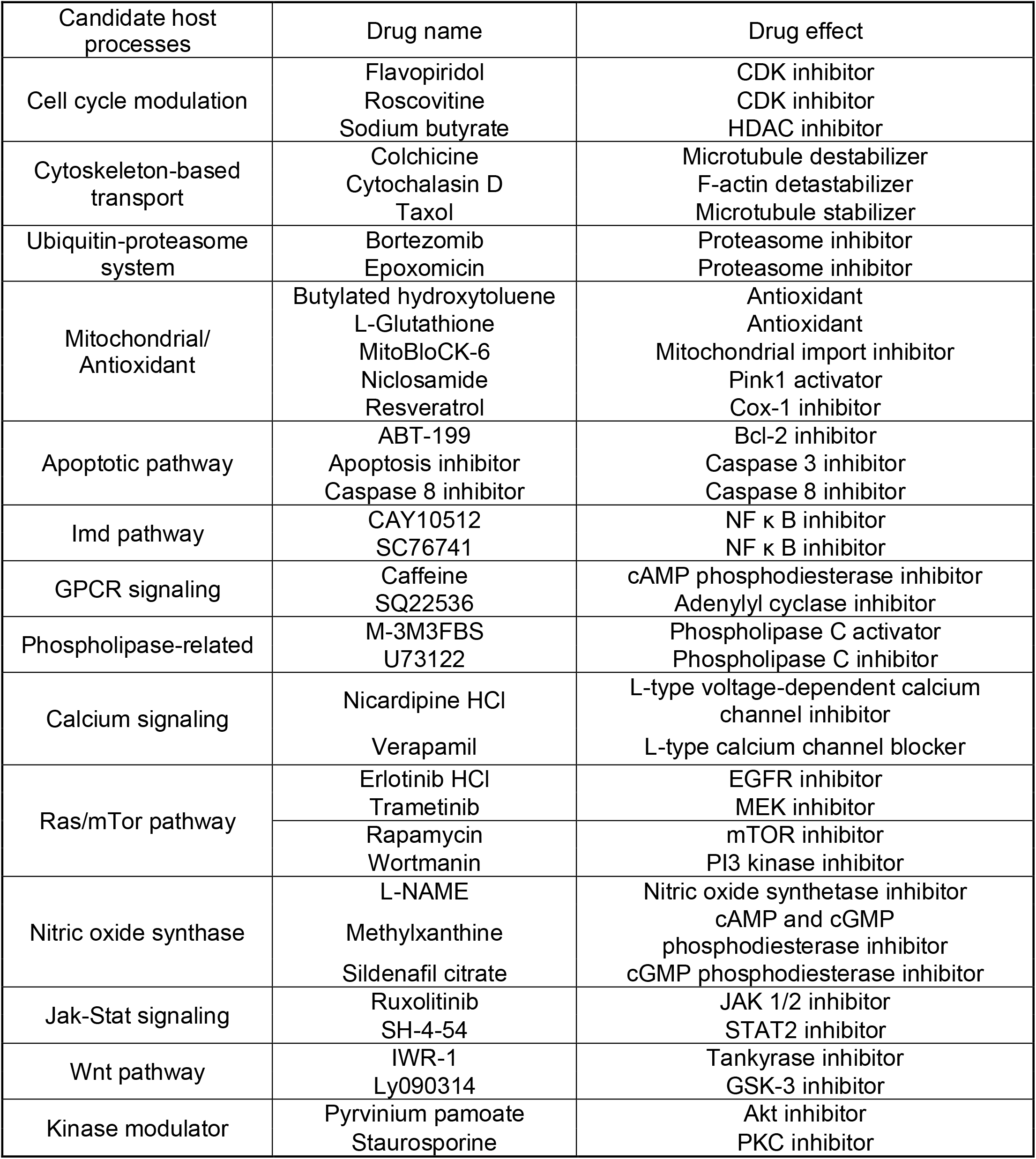
Screening targets and chemical tools

| Candidate host processes | Drug name | Drug effect |
| --- | --- | --- |
| Cell cycle modulation | Flavopiridol<br>Roscovitine<br>Sodium butyrate | CDK inhibitor<br>CDK inhibitor<br>HDAC inhibitor |
| Cytoskeleton-based transport | Colchicine<br>Cytochalasin D<br>Taxol | Microtubule destabilizer<br>F-actin detastabilizer<br>Microtubule stabilizer |
| Ubiquitin-proteasome system | Bortezomib<br>Epoxomicin | Proteasome inhibitor<br>Proteasome inhibitor |
| Mitochondrial/<br>Antioxidant | Butylated hydroxytoluene<br>L-Glutathione<br>MitoBloCK-6<br>Niclosamide<br>Resveratrol | Antioxidant<br>Antioxidant<br>Mitochondrial import inhibitor<br>Pink1 activator<br>Cox-1 inhibitor |
| Apoptotic pathway | ABT-199<br>Apoptosis inhibitor<br>Caspase 8 inhibitor | Bcl-2 inhibitor<br>Caspase 3 inhibitor<br>Caspase 8 inhibitor |
| Imd pathway | CAY10512<br>SC76741 | NF $\kappa$ B inhibitor<br>NF $\kappa$ B inhibitor |
| GPCR signaling | Caffeine<br>SQ22536 | cAMP phosphodiesterase inhibitor<br>Adenylyl cyclase inhibitor |
| Phospholipase-related | M-3M3FBS<br>U73122 | Phospholipase C activator<br>Phospholipase C inhibitor |
| Calcium signaling | Nicardipine HCl<br>Verapamil | L-type voltage-dependent calcium channel inhibitor<br>L-type calcium channel blocker |
| Ras/mTor pathway | Erlotinib HCl<br>Trametinib | EGFR inhibitor<br>MEK inhibitor |
|  | Rapamycin<br>Wortmanin | mTOR inhibitor<br>PI3 kinase inhibitor |
| Nitric oxide synthase | L-NAME<br>Methylxanthine<br>Sildenafil citrate | Nitric oxide synthetase inhibitor<br>cAMP and cGMP phosphodiesterase inhibitor<br>cGMP phosphodiesterase inhibitor |
| Jak-Stat signaling | Ruxolitinib<br>SH-4-54 | JAK 1/2 inhibitor<br>STAT2 inhibitor |
| Wnt pathway | IWR-1<br>Ly090314 | Tankyrase inhibitor<br>GSK-3 inhibitor |
| Kinase modulator | Pyrvinium pamoate<br>Staurosporine | Akt inhibitor<br>PKC inhibitor |

### Host-targeting small molecules alter *wsp* abundance in adult *Drosophila* hosts

The screening approach utilized absolute quantification of *wsp* by real-time PCR to assess the impact of the candidate drugs on body-wide *Wolbachia* titer (Christensen et al., 2019). *Drosophila melanogaster* flies, carrying the endogenous *w*Mel *Wolbachia* strain, were used for preliminary screening. Because this strain is double-balanced, it is referred to as DB wMel from this point forward. Female DB wMel flies were exposed to food supplemented with DMSO-solubilized drugs or control food exposing equivalent amounts of DMSO. A DMSO-solubilized rifampicin control was also run on every qPCR plate to confirm ongoing susceptibility of *Wolbachia* titer to compound treatments. For the chemical screen, and all other experiments performed in this study, *wsp* abundance was measured in pools of 5-day old flies by real-time PCR, using the absolute count method.

In the initial screen, treatments were tested for impact on whole-body *wsp* levels across 2 independent plate replicates. Treatments that significantly changed *wsp* abundance in both plates were identified as preliminary hits. Of 37 chemicals tested, the primary screen identified 16 compounds that significantly affected *wsp* counts. The 16 preliminary hit compounds were re-tested for reproducibility in a third plate replicate. 11 compounds were reconfirmed as hits, implicating a total of 9 host processes (Fig 2A) (Table 2) (Table S9-S12). Besides the rifampicin control, the proteasome inhibitor, bortezomib, was the only compound to significantly reduce whole-body *wsp* counts, ranging from 71% to as low as 48% of the DMSO control (p <0.001, n=6 amplifications per plate replicate). All other “hit” drugs elicited an increase in body-wide *wsp* abundance, with median values ranging 6%-57% above the control (p <0.001-0.036, n=6 per plate replicate) (Fig 2A) (Table 2) (Table S11-12). These results open the possibility of a generalized suppression effect, in which standard functions of multiple host pathways normally reduce whole-body *Wolbachia* loads.

**Figure 2.**
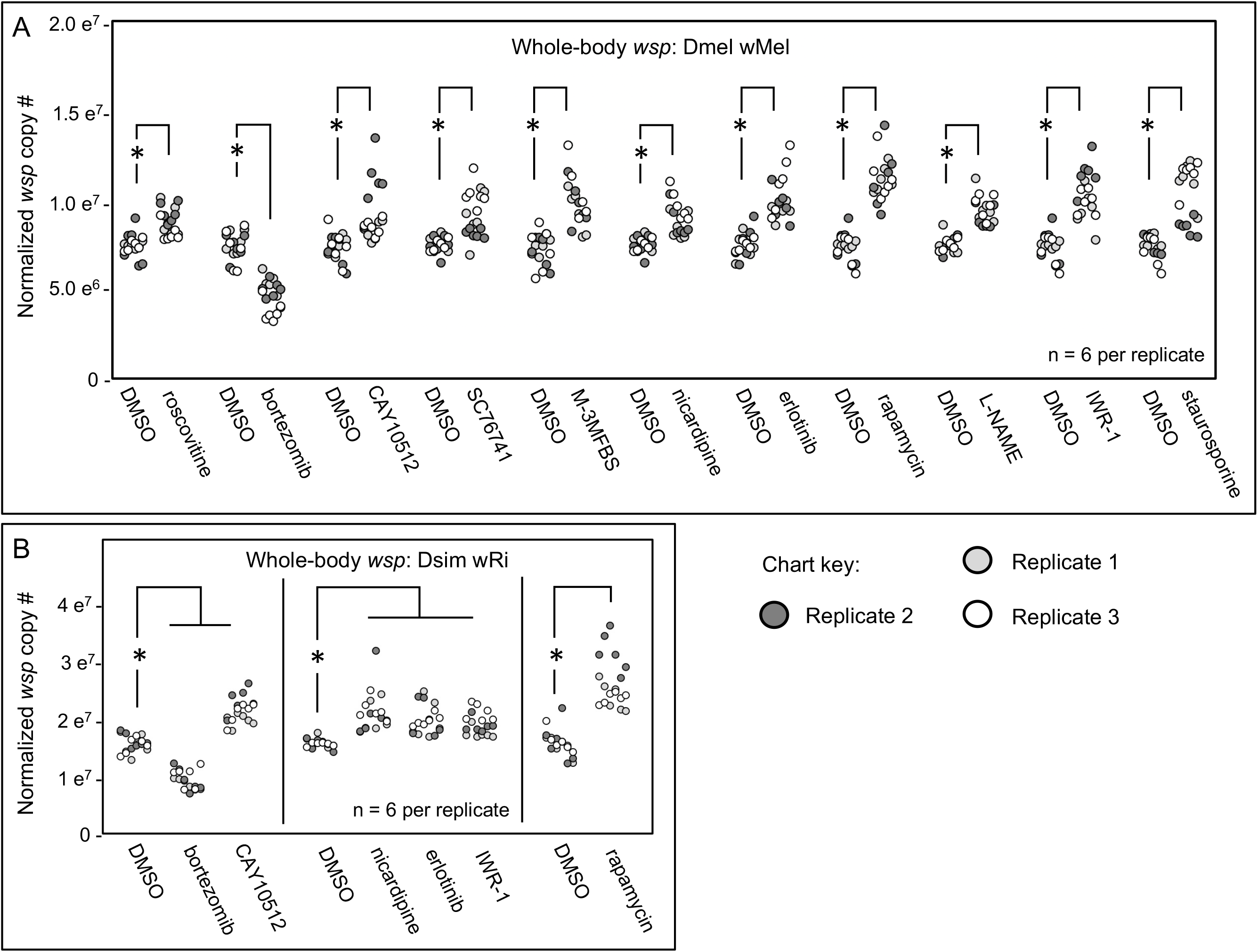
Whole body *wsp* abundance in response to chemical treatments. Display shows DMSO controls normalized across replicates, and the corresponding drug treatment data scaled accordingly. A) Chemical treatment effects on body-wide *wsp* abundance in Dmel wMel. B) Chemical treatment effects on body-wide *wsp* abundance in Dsim wRi. Significance was set at * p < 0.05, and is displayed only for conditions where all replicates met this standard.

**Table 2.** Chemical screen outcomes: comparing Dmel wMel hits to Dsim wRi results.

| Host cellular process | Drug name | p-value range for Dmel wMel | p-value range for Dsim wRi | Hit in both systems? |
| --- | --- | --- | --- | --- |
| Cell cycle modulation | Roscovitine | 0.003 - 0.009 | 0.006 - 0.283 | no |
| Ubiquitin-proteasome system | Bortezomib | <0.001 (all) | <0.001 (all) | yes |
| Imd pathway | CAY10512 | 0.001 - 0.036 | <0.001 - 0.001 | yes |
|  | SC76741 | <0.001 - 0.022 | 0.171 - 0.936 | no |
| Phospholipase-related | M-3M3FBS | 0.002 - 0.004 | 0.009 - 0.840 | no |
| Calcium signaling | Nicardipine HCl | <0.001 - 0.016 | <0.001 - 0.006 | yes |
| Ras/mTor pathway | Erlotinib HCl | <0.001 - 0.002 | <0.001 - 0.024 | yes |
|  | Rapamycin | <0.001 - 0.001 | <0.001 (all) | yes |
| Nitric oxide synthase | L-NAME | <0.001 - 0.002 | 0.001 - 0.056 | no |
| Wnt pathway | IWR-1 | <0.001 - 0.001 | <0.001 - 0.041 | yes |
| Kinase modulator | Staurosporine | 0.002 - 0.006 | 0.003 - 0.061 | no |

If host mechanisms exert conserved effects on the intracellular abundance of bacteria, we would expect *Wolbachia* titer regulation mechanisms to be shared across *Wolbachia*-host combinations as well. To investigate a role for candidate processes across systems, we retested all the 11 hits from DB wMel against the *D. simulans* (Dsim) model, which naturally carries the *w*Ri *Wolbachia* strain. The Dsim wRi re-screen yielded 6 treatments that significantly affected body-wide *wsp* counts across 3 plate replicates (Fig 2B) (Table 2) (Table S13-S14). In all cases, the observed effects were consistent with that seen previously for DB wMel. Bortezomib depleted median body-wide *wsp* counts to the range of 56%-69% of the DMSO control (p < 0.001, n=6 amplifications per plate replicate). By contrast, cay10512, nicardipine HCl, erlotinib HCl, rapamycin and IWR-1 treatments increased whole-body *wsp* abundance to 15%-52% above the DMSO control (p-value range: < 0.001-0.041, n=6 per plate replicate) (Fig 2B) (Table 2) (Table S13-S14). These findings implicate the host Ubiquitin-proteasome system, Imd signaling, Calcium signaling, Ras/mTOR signaling, and Wnt signaling as exerting generalized influence on *Wolbachia* titer across host-strain combinations.

### Constitutive genetic disruptions confirm some host pathway effects on *wsp* level

To examine the effect of host cellular processes on whole-body *Wolbachia* abundance, genetic manipulation experiments were performed, focusing on the 5 pathways that were dually implicated by chemical screening of DB wMel and Dsim wRi. These directed genetic experiments were performed using the GAL4::UAS expression system. GAL4 is a yeast-derived transcription factor, which drives expression of target genes by binding to the upstream activation sequence (UAS) adjacent to the target gene (Brand & Perrimon, 1993; Fischer et al., 1988). Thousands of GAL4 driver lines and UAS responder lines are publicly available, including UAS lines used for RNAi disruptions. GAL4-driven expression of each snap-back RNA construct creates a ds-RNA molecule that drives RNAi-based suppression of the corresponding gene product (Perkins et al., 2015; Perrimon et al., 2010). In this study, each host pathway was tested by two different UAS-dsRNA responder lines (Table S15). These responder lines were selected to emphasize re-testing the major protein target associated with each “hit” compound identified in the chemical screen (Table 2), while also addressing major targets associated with the same pathways (Table S15).

To carry out the genetic tests of host pathway effects on *Wolbachia*, the *w*Mel strain was crossed into well-established GAL4 driver lines that drive constitutive whole-body expression, including the reputedly “strong” Actin5C-GAL4 driver (*Act-5C*), and the “milder” daughterless-GAL4 driver (*da-GAL4*) (Table S16). Few to no F1 progeny were recovered that carried *Act5C-GAL4* as well as *UAS-dsRNA* chromosomes, indicating lethality for such genetic combinations. However, crossing the *UAS-dsRNA* lines to *da-GAL4* (Serbus et al., 2015) yielded ample F1 flies for analysis that are capable of RNAi expression. Control flies were collected in parallel, including non-expressing F1 siblings (when available), as well as F1 progeny generated by outcrossing the *da-GAL4* driver to OreR males.

Using this strategy, constitutive RNAi expression tools confirmed effects of some host pathways, but not others, on whole-body *wsp* abundance (Table S17, Table S18). Inconsistent *wsp* abundance was detected in association with *da-GAL4::UAS-dsRNA* progeny carrying disruptions in the Ubiquitin-proteasome system genes *Ubc6*, which encodes an E2 subunit, and *Prosalpha6*, which encodes a 20S proteasomal subunit. Inconsistent *wsp* levels were also associated with disruptions to the Imd pathway, specifically knockdown of NF-kappa-B/*Rel*, which encodes a culminating element of the Imd cascade, and *Tak1*, involved in Rel activation. No changes in *wsp* abundance were detected in response to disruptions of Calcium signaling by knockdown of *Ca1alphaD* and *Cac*, which both encode proteins with homology to voltage-gated, L-type calcium channels (Table S17, Table S18). Thus, the genetic data did not corroborate an effect for these host pathways on whole-body *Wolbachia* abundance.

The genetic data did show a significant *wsp* response to constitutive RNAi disruption of the host Wnt pathway. The *shaggy (sgg)* gene, encodes a GSK-3 protein that normally stabilizes β-catenin (Wu & Pan, 2010), was initially targeted by dsRNA, yielding inconsistent *wsp* abundance measurements (Table S17, Table S18). However, dsRNA disruption of *armadillo (arm)*, the fly homologue of β-catenin, increased median whole-body *wsp* counts to 9-15% above the OreR-outcrossed control (p-value range: <0.001-0.034, n = 6 per plate replicate) (Fig. 3) (Table S17) (Table S18).

**Figure 3.**
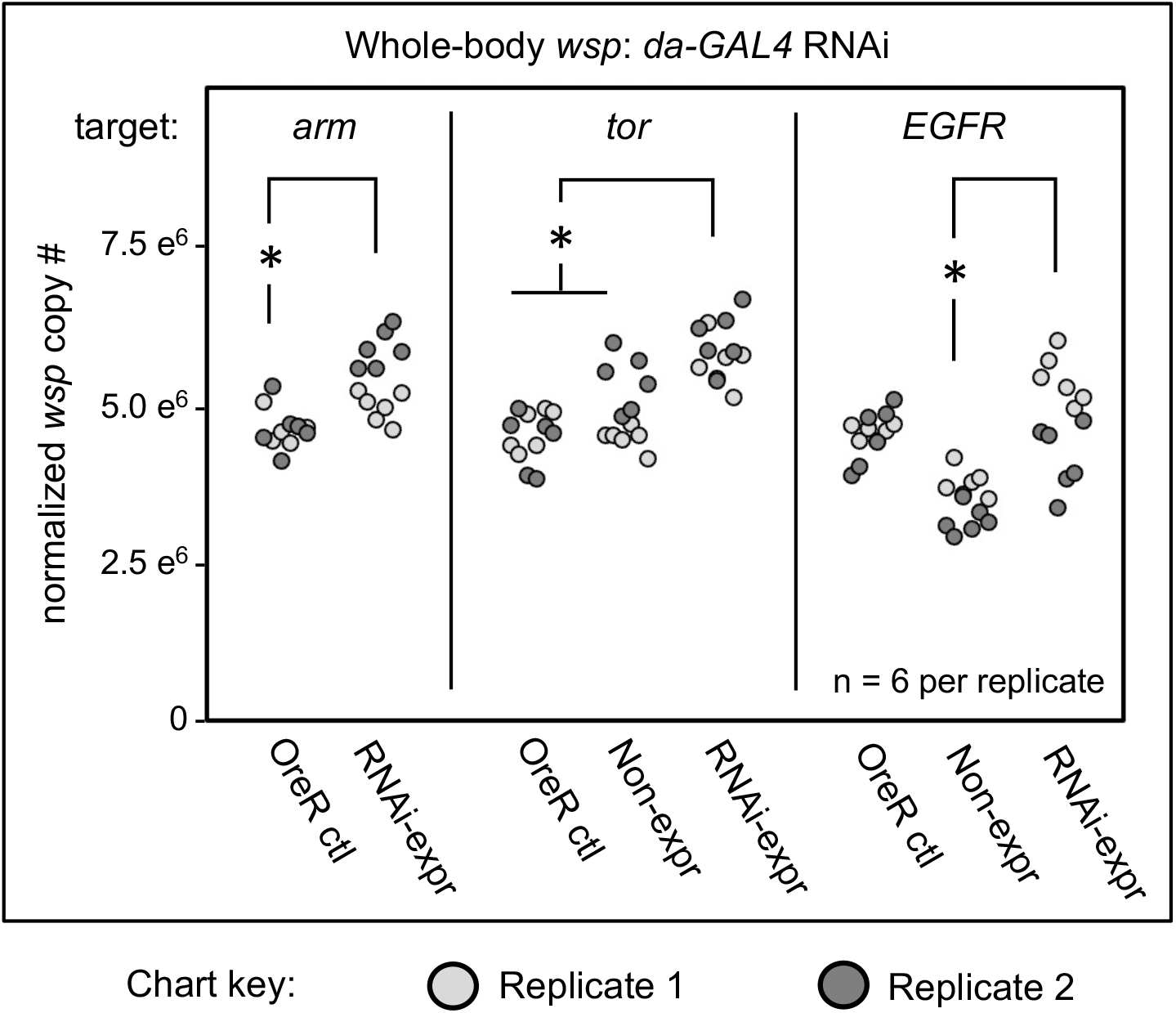
Whole body *wsp* abundance in control vs. *da-GAL4:UAS-RNAi* knockdown flies. Oregon-R outcross controls are included throughout, and non-expressing sibling controls are shown where available. Display shows OreR controls normalized across replicates, and the corresponding conditions scaled accordingly. Panel shows data from 2 independent biological replicates. RNAi disruption conditions shown from left to right: *armadillo, tor*, and *EGFR*. Significance was set at * p < 0.05, and is displayed only for conditions where all replicates met this standard.

A significant *wsp* response was also detected in response to constitutive RNAi disruption of the Ras/mTor pathway. *Target of rapamycin (Tor)* disruption flies exhibited higher whole-body *wsp* abundance, at 23-31% above sibling controls as well as OreR-outcrossed controls (p<0.001, n = 6 per plate replicate) (Fig. 3) (Table S17) (Table S18). Ras/mTOR signaling was also retested by knockdown of the *Epidermal growth factor receptor (EGFR)*, a major regulator of Ras activity. Comparing the EGFR condition to OreR-outcrossed controls yielded inconsistent *wsp* abundance measurements. However, in comparison to sibling controls, EGFR disruption yielded a 35-44% increase in median *wsp* counts (p-value range: < 0.001-0.002, n = 6 per plate replicate) (Fig. 3) (Table S17) (Table S18). This suggests that detection of *wsp* responses is at least somewhat context-dependent.

To confirm an effect of Wnt and Ras/mTor pathways on *Wolbachia*, the strongest *da-GAL4::UAS-dsRNA* outcomes were retested. *arm* RNAi elicited a 16-50% increase in median *wsp* abundance over the OreR-outcrossed control (p < 0.001, n = 18) (Figure 4A) (Table S19) (Table 20). Power analysis indicated the *arm* RNAi outcome to be robust (β < 0.003 at n ≥ 12; total n = 18) (Fig. 4B). *Tor* RNAi triggered a 38-39% increase in *wsp* abundance as compared to OreR-outcrossed controls (p < 0.001, n = 18) (Figure 4C) (Table S19) (Table 20). These results were also well-supported by power analysis (β < 0.002 at n ≥ 4; total n = 18) (Fig. 4D). Taken together, these data suggest that RNAi disruption of Wnt and Ras/mTOR signaling, even when driven by the mild *da-GAL4* driver, significantly affects *Wolbachia* load carried by whole insects.

**Figure 4.**
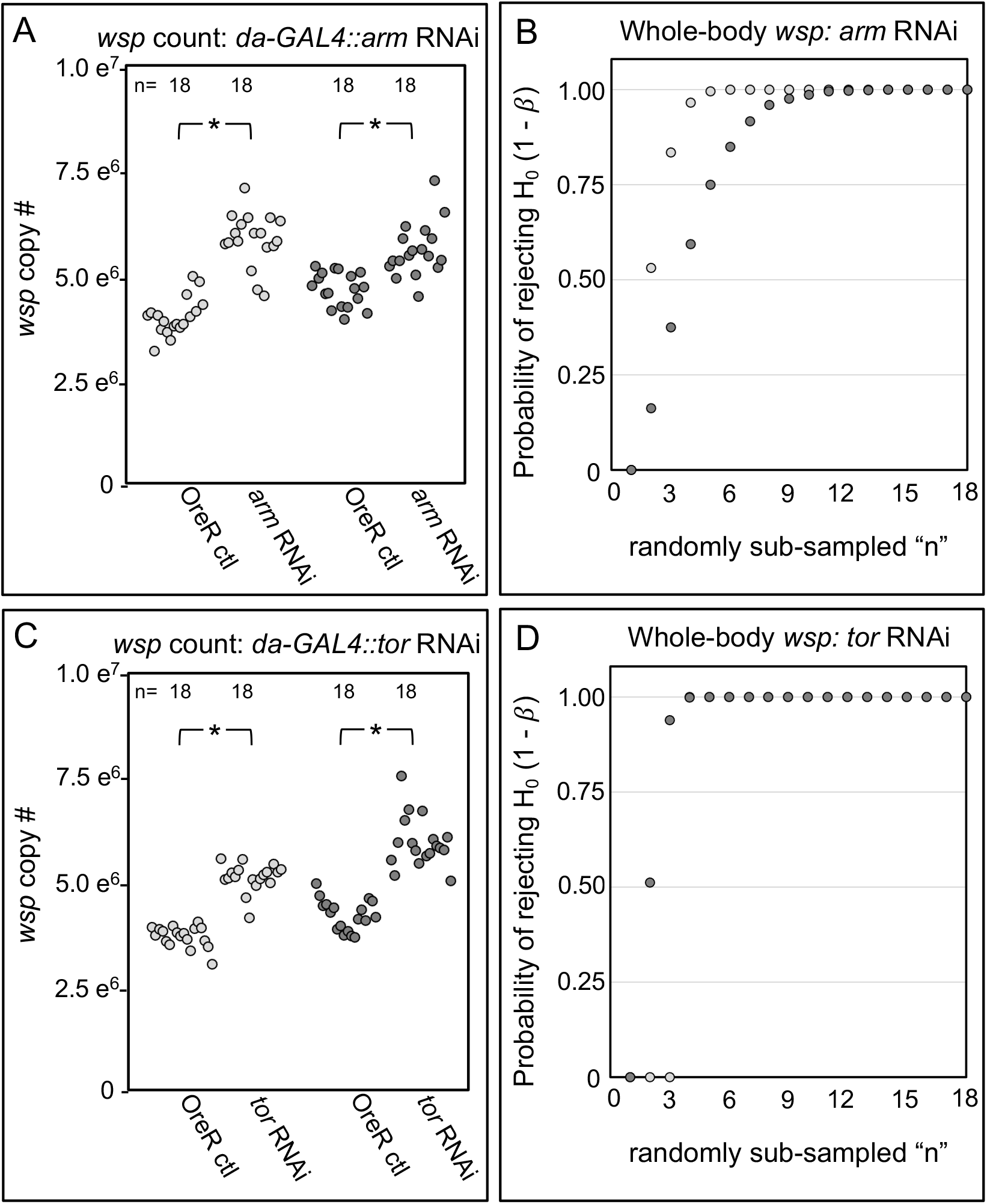
Whole body *wsp* abundance in control vs. *da-GAL4::UAS-RNAi* knockdown flies. Panels show data from 2 independent biological replicates. A) Body-wide *wsp* abundance in *arm-*RNAi conditions. B) Power analysis, testing the likelihood of significance as a function of sample size in Panel A. C) Body-wide *wsp* abundance in *tor-*RNAi conditions. D) Power analysis, testing the likelihood of significance in Panel C. Significance was set at * p < 0.05.

### Confirmation of host effects on *wsp* by inducting genetic disruptions in adult hosts

An intrinsic aspect of many genetic disruption tools, including classical mutants and RNAi induction experiments, is that they can generate cumulative disruption effects. This distinguishes the *da-GAL4::dsRNA* experiments from adult-only chemical disruptions performed in this study. To test for an adult-specific role for host pathways in regulating whole-body *wsp* abundance, “Gene-switch” GAL4 driver flies were used. The Gene-switch version of GAL4 carries an inhibitory domain that blocks GAL4 function, until a de-repressor compound, Mifepristone, is added (Roman et al., 2001). Publicly available stocks were used to create *Wolbachia-*infected *GS-GAL4::UAS-dsRNA* flies, then *wsp* abundance was measured after 2 weeks of Mifepristone exposure. Initial trials on DB wMel flies yielded some extent of background effects from Mifepristone use, with median *wsp* counts increasing by 13-15% (p-value range: < 0.001-0.011, n = 18). Power analysis indicated Mifepristone effects on *wsp* to be mild (β > 0.05 at n = 18) (Supplementary Figure S1) (Table S21) (Table S22).

To test an adult function for the Wnt pathway, flies that are genetically capable of mifepristone-induced *arm* dsRNA expression were generated, referred to as “RNAi+”, as compared to non-expressing “RNA-” siblings. No significant difference in *wsp* level was observed between control RNAi– siblings nor DMSO-fed *arm* RNAi+ flies. By contrast, mifepristone-fed RNA+ flies showed a significant decrease in *wsp* abundance, ranging from 45-71% of the *wsp* count exhibited by all other control conditions (p-value range: < 0.001-0.033, n = 9), suggesting that *arm* disruption reduces *Wolbachia* titer (Figure 5A) (Table S23) (Table S24). Adult-specific *arm* effects are consistent with ongoing *Wolbachia* sensitivity to Wnt signaling. The data also demonstrate that Mifepristone-fed conditions do not necessarily increase *wsp* counts by default.

**Figure 5.**
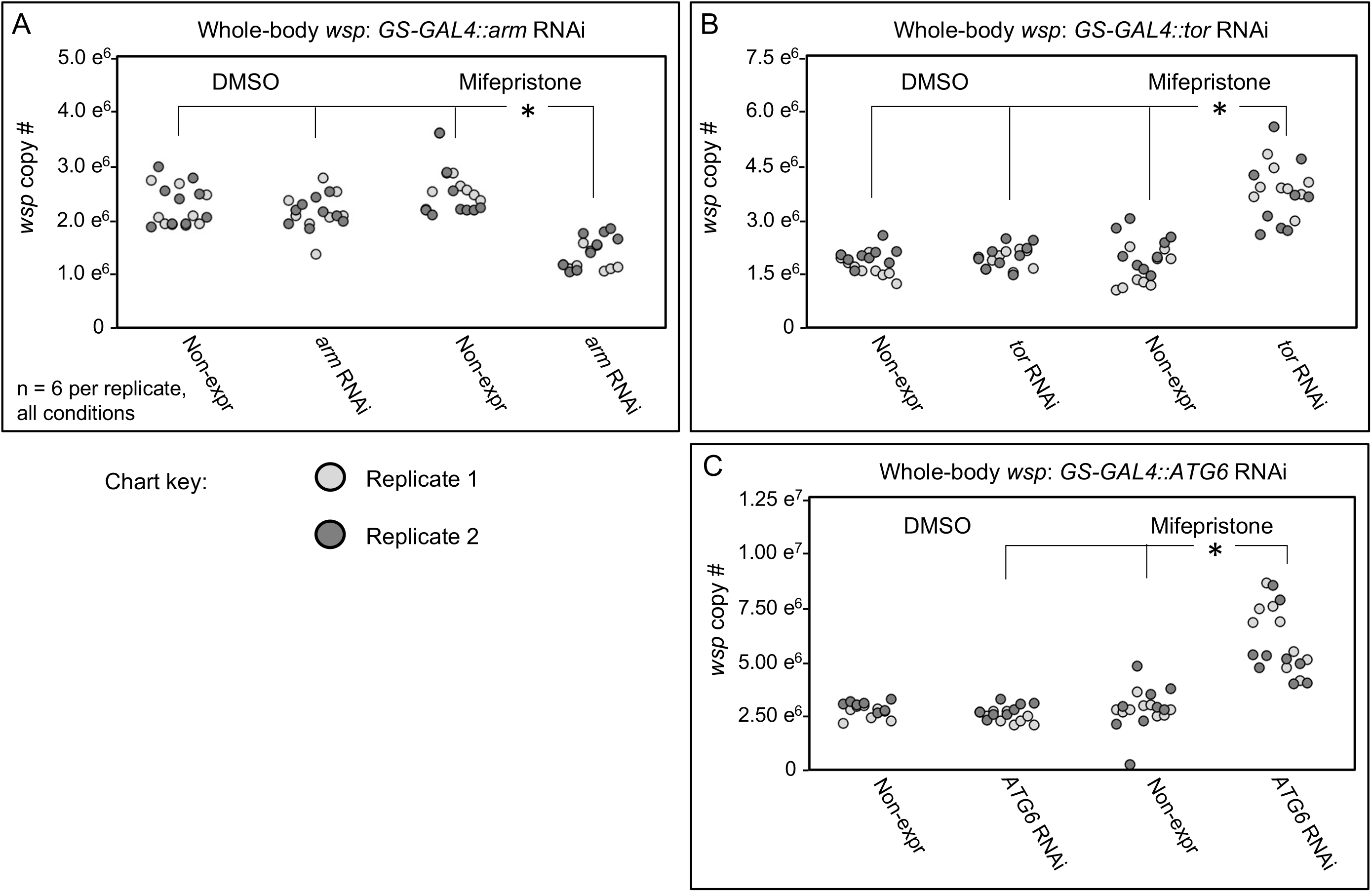
Whole body *wsp* abundance in control vs. *GSGAL4::UAS-RNAi* knockdown flies. Flies capable of dsRNA expression were compared against nonexpressing siblings, in the presence of DMSO or Mifepristone dissolved into DMSO. Genetic disruptions tested: A) *arm* RNAi, B) *tor* RNAi, C) *ATG6* RNAi. Data out of range for panel C: 3 outliers for the Non-expressing DMSO condition at 2.01×10^7^, 2.54×10^7^, 2.064×10^7^, and 1 outlier for the ATG6 RNAi DMSO condition at 2.28×10^7^. Significance was set at * p < 0.05, and is displayed only for conditions where all replicates met this standard.

Adult-induced genetic tests of the Ras/mTOR pathway continued to focus on disruption of *tor* gene expression. As for the *arm* disruption tests above, no significant differences were observed in *wsp* levels when control RNAi– siblings were compared to each other and to DMSO-fed *tor* RNAi+ flies. However, flies expected to carry a functional *tor* disruption, specifically the mifepristone-fed *tor* RNAi+ condition, carried 81%-184% more *wsp* than all other conditions (p-value range: < 0.001-0.031, n = 9) (Figure 5B) (Table S23) (Table S24). This is consistent with all other evidence showing Ras/mTor disruption elevates whole-body *wsp*.

### Test of autophagy as a convergence point for regulating *Wolbachia* abundance

One way to reconcile the effects for Wnt and Tor effects on *wsp* abundance is to consider the possibility that both act on a consensus *Wolbachia* regulator. The literature indicates that Wnt can activate autophagy in some instances, but usually suppresses it (Pérez-Plasencia et al., 2020), perhaps in part by down-regulating expression of Beclin-1, also known as ATG6 (Tao et al., 2017), mTORC1 is known to inhibit ATG6, and thus autophagy as well, by suppressing the protein ULK1 that activates ATG6 (Hill et al., 2019). These findings position ATG6 as a possible convergence point for Wnt- and Tor-based suppression of *Wolbachia* abundance. If so, one prediction would be that ATG6 disruption should reduce *wsp* abundance in vivo (Figure 6A).

**Figure 6.**
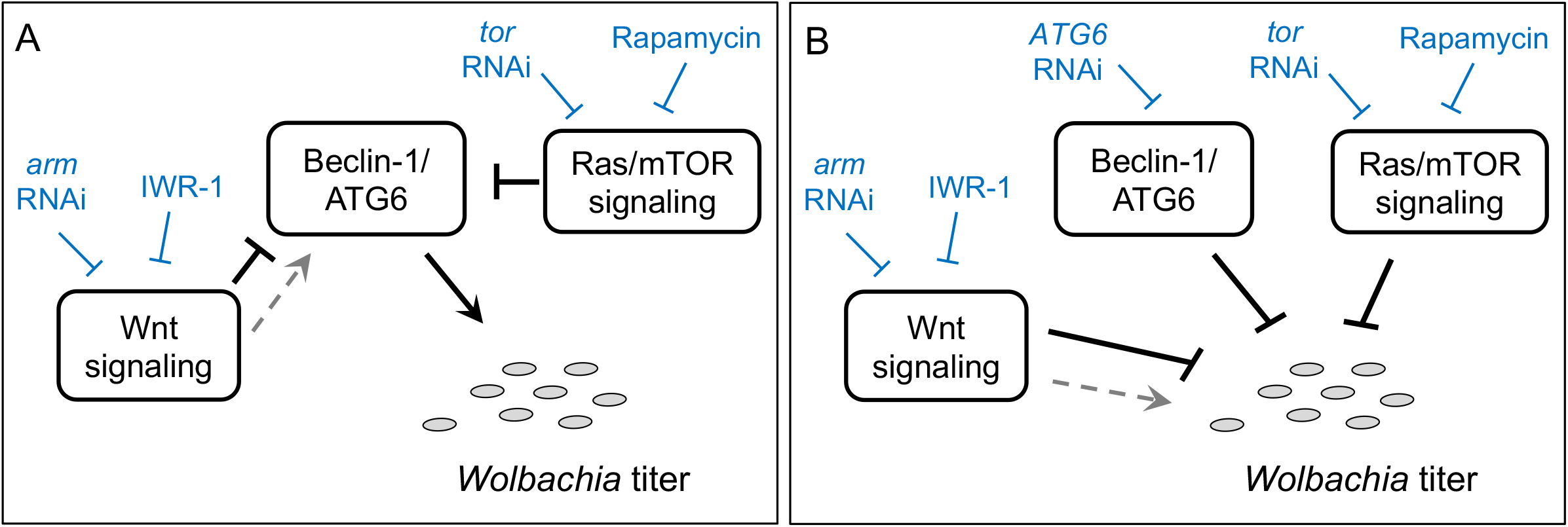
Models of host effects on whole-body *Wolbachia* titer. A) Prediction for possible Beclin-1/ATG6 effects on *Wolbachia* titer in vivo. B.) Results summary, presented as a model with consideration of Beclin-1/ATG6 experimental data thus far. Blue: chemical and RNAi disruption results. Boxes: host functions. Grey ovals: *Wolbachia* bacteria.

To test the effect of *ATG6* on whole body *Wolbachia* abundance, the *GS-GAL4::UAS-dsRNA* system was used. The condition expected to disrupt *ATG6* function, mifepristone-fed *ATG6* RNAi+, did not differ significantly from the DMSO-fed *ATG6* RNAi-condition, due to statistical outliers in the DMSO control (p = 0.326, n = 9) (Figure 5C) (Table S23) (Table S24). However, the mifepristone-fed *ATG6* RNAi+ condition did exhibit 67-173% higher *wsp* levels than all other conditions (p-value range: < 0.001-0.045, n = 9) (Figure 5C) (Table S23) (Table S24). These data imply that ATG6 normally suppresses whole body *wsp* abundance, consistent with studies that implicate autophagy as a suppressor of *Wolbachia* titer (Figure 6B) (Strunov et al., 2022; Voronin et al., 2012).

## DISCUSSION

In this study, the relationships between *Wolbachia* and the *Drosophila* host were explored by investigating the effect of candidate host processes on *Wolbachia* titer. We performed a pilot drug screen coupled with absolute quantification of the *wsp* gene from whole insect samples, using real-time qPCR. Tissue culture cells have been performed in the past as a comprehensive approach to begin addressing *Wolbachia-*host interactions (Grobler et al., 2018; White et al., 2017a). The organismal feeding approach provides the advantages of a whole-body exposure, an in-vivo response, measurements free from possible cell invasion artifacts (White, Pietri, et al., 2017) and ease of retesting in new, non-model systems (Momtaz et al., 2020). There are limits to the resolution of an organismal feed, as a whole-body readout does not inform the similarity of *wsp* response across tissues within the organism. Absolute counts by real-time PCR remain critical for the success of such efforts, to avoid artifacts attributable to variable host ploidy, which may not always be forseeable across tissues, systems, organismal age, and nutritional conditions (Christensen et al., 2019; Ren et al., 2020).

The pilot chemical screen in this study targeted candidate cellular functions that are associated with human pathogenic infections, to test their involvement in commensal *Wolbachia* infection. The screen yielded 11 compounds that consistently altered body-wide *wsp* in *w*Mel-infected *D. melanogaster*, 6 of which repeated in *w*Ri-infected *D. simulans*. Based upon the literature, these “hit” compounds reflect 5 consensus biological processes, namely the Ubiquitin-proteasome pathway, the Imd pathway, Calcium signaling, the Ras/mTOR pathway, and the Wnt pathway (Table 1). As for any drug-based screen, negative results could be due to a basic incompatibility of the experimental system with the drug, rather than the lack of target involvement in the process of interest. Heterogeneous drug availability across functional classes is also an intrinsic attribute of this and all chemical screening analyses. Given the number and diversity of host treatments that affected *wsp* in *D. melanogaster* but not *D. simulans*, it is likely that many system-specific host regulators of endosymbiont abundance still await discovery.

A surprising outcome in this study is the similarity of *Wolbachia* responses to host-side chemical treatments. The proteasome inhibitor, Bortezomib, reduced whole body *wsp* levels, consistent *Wolbachia* titer reductions due to nutritional depletion and/or failure to eliminate *Wolbachia-*suppressing agents (White et al., 2017a). By contrast, all other drug treatments that significantly affected *wsp* yielded an apparent titer increase, with a subset of effects repeating across Dmel and Dsim experimental systems. *Wolbachia* over proliferation was previously reported in response to ribosome disruption (Grobler et al., 2018). Perhaps *Wolbachia* abundance is collectively suppressed by a suite of host cellular processes. In such case, disruption of the relevant host cell functions allows a favorable shift in *Wolbachia* life cycle dynamics. It makes sense to expect commensalism as an artifact of endosymbiont genome reduction, with virulence factors eliminated over time, along with most other genes (Latorre & Manzano-Marín, 2017; Sachs et al., 2014). However, *Wolbachia* suppression by host cellular mechanisms demonstrates that commensalism is not necessarily free of conflict (Keeling & McCutcheon, 2017). Rather, the data suggest that commensalism is achieved in part through ongoing, active containment of the endosymbiont by its host.

A distinctive aspect of this study is use of RNAi testing to confirm the basis for chemical disruption effects. Several host processes implicated in *Wolbachia* titer control by chemical screening were not phenocopied by RNAi. Such results are unfortunately inconclusive, as internal redundancies may render certain knockouts ineffective, and developmental tolerance limits may preclude analysis of the strongest knockdown effects. Within that context, this study consistently identified *wsp* abundance as sensitive to host Wnt and Ras/mTor signaling (Figure 6). *wnt*-related disruptions elevated *wsp* levels when chemical and constitutive genetic disruption tools were used, but decreased *wsp* when the inducible Gene-Switch RNAi tools were used. *tor*-related disruptions consistently increased whole-body *wsp* abundance, regardless of the disruption method used (Figure 6). Perhaps the lengthy duration of the Gene-Switch experiments triggers activation of compensatory functions in some pathways, but not others. Due to this, the precise implication of Beclin-1/ATG6 disruption effects on *Wolbachia* remains open at this time (Figure 6B). An additional consideration is the finding that autophagy feedback mechanisms can de-activate *wnt* signaling (Petherick et al., 2013). The extent of connectivity versus independence for Wnt, ATG6 and Ras/mTor effects on whole-body *Wolbachia* abundance thus awaits further study (Figure 6). Autophagy involvement is supported to some extent by this work as well as host-pathogen literature (Table S24), highlighting autophagy as a consensus aspect of diverse infection mechanisms.

## DATA AVAILABILITY

All datasets generated by this study are included in the accompanying Supplementary Tables.

## AUTHOR CONTRIBUTIONS

ZS, HS, RAB and LS designed the experiments.

ZS, HS, RAB and LS supervised the experiments.

ZS, HS and RAB conducted the experiments.

ZS, HS, RAB and LS analyzed and discussed the data.

ZS and LS wrote the manuscript.

ZS, HS, RAB and LS reviewed the manuscript.

## CONFLICT OF INTEREST

This research was conducted in the absence of any commercial or financial relationships that could be construed as a potential conflict of interest.

## ACKNOWLEDGEMENTS

Funding for this project came from FIU Academic Affairs and the NSF Division of Integrated Organismal Systems (#1656811).

We sincerely thank Steen Christensen, Moises Camacho, Anthony Bellantuono, AJM Zehadee Momtaz, Erasmo Perera and the FIU Department of Biological Sciences for helpful discussions and logistical support. We also thank the FIU College of Arts and Sciences and the University Graduate School for supporting our students.

## Notes

### Competing Interest Statement

The authors have declared no competing interest.

