## Supplemental files for "Host pathways associated with human bacterial infections extend to commensal *Wolbachia-Drosophila* endosymbiosis"

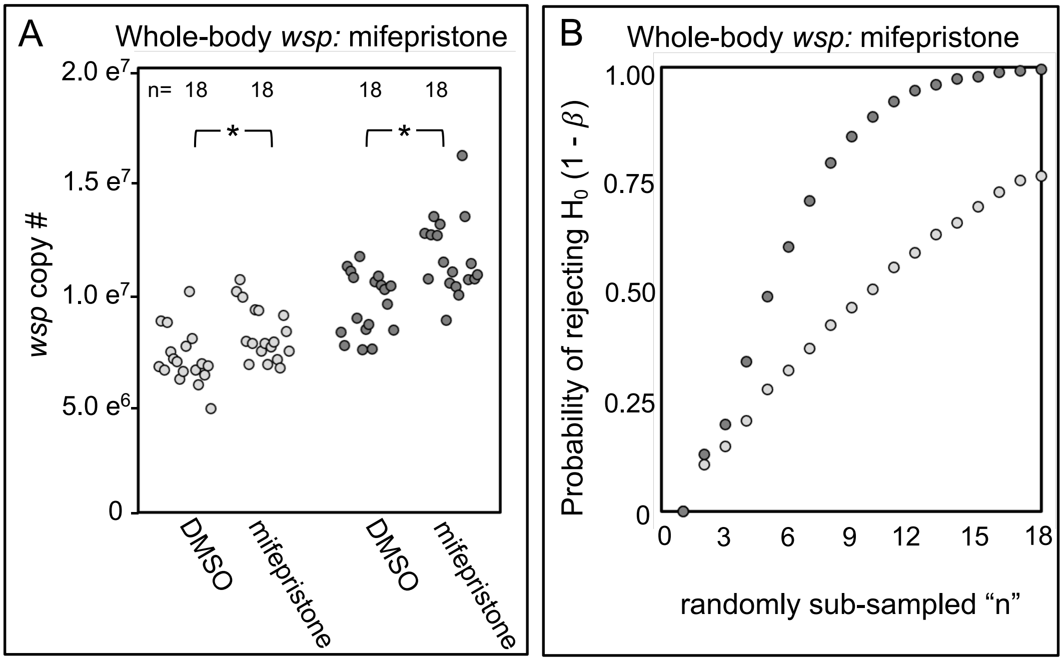


Figure S1. Whole body *wsp* abundance in control vs. *da-GAL4:UAS-RNAi* knockdown flies. Panels show data from 2 independent biological replicates. A) Body-wide *wsp* abundance in flies exposed to carrier DMSO versus Mifepristone solubilized in DMSO. B) Power analysis, testing the likelihood of significance as a function of sample size in Panel A. Significance was set at * $p < 0.05$.

**Table S1. Chemicals used**

| Candidate host processes | Drug name | CAS # | Catalog # | Vendor |
|---|---|---|---|---|
| Cell cycle modulation | Flavopiridol<br>Roscovitine<br>Sodium butyrate | 146426-40-6<br>186692-46-6<br>156-54-7 | S1230<br>A1723<br>B5887 | Selleckchem<br>ApexBio<br>Sigma |
| Cytoskeleton based transport | Colchicine<br>Cytochalasin D<br>Taxol | 64-86-8<br>22144-77-0<br>33069-62-4 | C9754<br>B6645<br>T7402 | Sigma<br>ApexBio<br>Sigma |
| Ubiquitin-proteasome system | Bortezomib<br>Epoxomicin | 179324-69-7<br>134381-21-8 | S1013<br>S7038 | Selleckchem<br>Selleckchem |
| Mitochondrial/ Antioxidant | Butylated hydroxytoluene<br>L-Glutathione<br>MitoBloCK-6<br>Niclosamide<br>Resveratrol | 128-37-0<br>70-18-8<br>303215-67-0<br>50-65-7<br>501-36-0 | AAA1686330<br>G4251<br>505759<br>N3510<br>R5010 | Thermo Fisher<br>Sigma<br>Sigma<br>Sigma<br>Sigma |
| Apoptotic pathway | ABT-199<br>Apoptosis inhibitor<br>Caspase 8 inhibitor | 1257044-40-8<br>54135-60-3<br>210344-98-2 | 6960<br>178488<br>SC-3084 | Tocris<br>Sigma<br>Santa cruz |
| Imd pathway | CAY10512<br>SC76741 | 139141-12-1<br>913822-46-5 | SC-205237<br>SML2382 | Santa cruz<br>Sigma |
| GPCR Signaling | Caffeine<br>SQ22536 | 58-08-2<br>17318-31-9 | C0750<br>S8283 | Sigma<br>Selleckchem |
| Phospholipase-related | M-3M3FBS<br>U73122 | 200933-14-8<br>112648-68-7 | T5699<br>1268 | Sigma<br>Tocris |
| Calcium signaling | Nicardipine HCl<br>Verapamil | 259-198-4<br>152-11-4 | N7510<br>V4629 | Sigma<br>Sigma |
| Ras/mTor pathway | Erlotinib HCl<br>Trametinib | 183319-69-9<br>871700-17-3 | SML2156<br>S2673 | Sigma<br>Selleckchem |
| | Rapamycin<br>Wortmanin | 53123-88-9<br>19545-26-7 | 553210<br>S2758 | Sigma<br>Selleckchem |
| Nitric oxide synthase | L-NAME<br>Methylxanthine<br>Sildenafil citrate | 51298-62-5<br>6136-37-4<br>171599-83-0 | N5751<br>69720<br>3784 | Sigma<br>Sigma<br>Tocris |
| Jak-Stat signaling | Ruxolitinib<br>SH-4-54 | 941678-49-5<br>1456632-40-8 | S1378<br>B4789 | Selleckchem<br>ApexBio |
| Wnt pathway | IWR-1<br>Ly090314 | 1127442-82-3<br>603288-22-8 | B2306<br>SML1438 | ApexBio<br>Sigma |
| Kinase modulator | Pyrvinium pamoate<br>Staurosprine | 3546-41-6<br>62996-74-1 | P0027<br>S1421 | Sigma<br>Selleckchem |
| Antibiotic control | Rifampicin | 13292-46-1 | R3501 | Sigma |
| GS-GAL4 activator | Mifepristone | 84371-65-3 | M8046 | Sigma |
| solvent control | DMSO | 67-68-5 | D8418 | Sigma |

**Table S2. Transgenic UAS-dsRNA lines used in this study**

| Process | Stock # | Gene name | Insertion type | Insertion # | Chromosome |
|---|---|---|---|---|---|
| Ubiq-Prot | 42631 | *Ubc6* | VALIUM20 Trip line | HMS02466 | 2 |
| | 53974 | *Prosalpha6* | VALIUM20 Trip line | HMJ21359 | 2 |
| Imd | 33661 | *Relish* | VALIUM20 Trip line | HMS00070 | 3 |
| | 53377 | *Tak1* | VALIUM20 Trip line | HMJ02053 | 2 |
| Ca channel | 77174 | *Cacophony* | VALIUM20 Trip line | HMC03320 | 2 |
| | 33413 | *Ca-alpha1D* | VALIUM20 Trip line | HMS00294 | 3 |
| Ras/mTOR | 60012 | *Egfr* | VALIUM20 Trip line | HMS05003 | 2 |
| | 58312 | *Tor* | VALIUM20 Trip line | HMJ22419 | 2 |
| Wnt | 35004 | *Armadillo* | VALIUM20 Trip line | HMS01414 | 3 |
| | 38293 | *Sgg* | VALIUM20 Trip line | HMS01751 | 2 |
| Autophagy | 37541 | ATG6 | VALIUM20 Trip line | HMS01483 | 3 |

**Table S3. Whole body *wsp* abundance, in response to bortezomib treatment**

| *wsp* absolute counts, whole-body samples DMSO vs bortezomib treatment | | |
|---|---|---|
| Condition | Plate 1 | Plate 2 |
| DMSO | 2.69E+06 | 2.11E+06 |
| Whole body | 2.79E+06 | 2.16E+06 |
| Well 1 | 2.76E+06 | 2.17E+06 |
| DMSO | 2.45E+06 | 2.63E+06 |
| Whole body | 2.36E+06 | 2.53E+06 |
| Well 2 | 1.97E+06 | 2.10E+06 |
| DMSO | 2.48E+06 | 2.30E+06 |
| Whole body | 2.62E+06 | 2.23E+06 |
| Well 3 | 2.44E+06 | 2.21E+06 |
| DMSO | 2.29E+06 | 2.51E+06 |
| Whole body | 2.40E+06 | 2.53E+06 |
| Well 4 | 2.18E+06 | 2.39E+06 |
| DMSO | 2.53E+06 | 2.22E+06 |
| Whole body | 2.42E+06 | 2.02E+06 |
| Well 5 | 2.12E+06 | 2.08E+06 |
| DMSO | 3.17E+06 | 2.58E+06 |
| Whole body | 3.53E+06 | 2.50E+06 |
| Well 6 | 3.41E+06 | 2.31E+06 |
| Bortezomib | 1.25E+06 | 9.86E+05 |
| Whole body | 1.25E+06 | 1.03E+06 |
| Well 1 | 1.20E+06 | 9.55E+05 |
| Bortezomib | 1.28E+06 | 1.24E+06 |
| Whole body | 1.21E+06 | 1.13E+06 |
| Well 2 | 1.24E+06 | 1.13E+06 |
| Bortezomib | 1.22E+06 | 1.11E+06 |
| Whole body | 1.33E+06 | 1.20E+06 |
| Well 3 | 1.25E+06 | 1.11E+06 |
| Bortezomib | 1.36E+06 | 1.11E+06 |
| Whole body | 1.28E+06 | 1.11E+06 |
| Well 4 | 1.14E+06 | 1.45E+06 |
| Bortezomib | 1.37E+06 | 1.18E+06 |
| Whole body | 1.41E+06 | 9.42E+05 |
| Well 5 | 1.31E+06 | 9.39E+05 |
| Bortezomib | 1.45E+06 | 1.25E+06 |
| Whole body | 1.64E+06 | 1.20E+06 |
| Well 6 | 1.47E+06 | 1.10E+06 |

**Table S4. Statistical analyses of *wsp* abundance from whole body samples, in response to bortezomib treatment**

| Plate replicate # | Shapiro-wilk: p-values, interpretation | | | Levene's test: p-values, interpretation | | Outcome: analysis of different combinations | | | "n" required to satisfy $\beta < 0.05$ |
|---|---|---|---|---|---|---|---|---|---|
| | DMSO | Bortezomib | Normal distribution? | Overall value | Variance homogeneous? | Analysis required | Resulting p-value | Significantly different? | |
| 1 | 0.087 | 0.107 | both normal | 0.014 | no | Welch's T-test | <0.001 | yes | 4 |
| 2 | 0.166 | 0.171 | both normal | 0.024 | no | Welch's T-test | <0.001 | yes | 3 |

* represent whole body comparisons, all technical replicates, n = 18

**Table S5. Absolute quantification of *wsp* copy number from ovarian samples, in response to bortezomib treatment**

| *wsp* absolute counts from ovarian samples DMSO vs bortezomib treatment | | |
|---|---|---|
| Condition | Plate 1 | Plate 2 |
| DMSO Ovary only Well 1 | 1.39E+06 | 1.09E+06 |
| | 1.43E+06 | 1.05E+06 |
| | 1.41E+06 | 9.65E+05 |
| DMSO Ovary only Well 2 | 1.41E+06 | 1.00E+06 |
| | 1.38E+06 | 1.09E+06 |
| | 1.40E+06 | 9.89E+05 |
| DMSO Ovary only Well 3 | 1.26E+06 | 1.10E+06 |
| | 1.17E+06 | 1.04E+06 |
| | 1.07E+06 | 1.01E+06 |
| DMSO Ovary only Well 4 | 9.35E+05 | 7.35E+05 |
| | 9.11E+05 | 8.33E+05 |
| | 9.36E+05 | 8.61E+05 |
| DMSO Ovary only Well 5 | 1.19E+06 | 9.34E+05 |
| | 1.13E+06 | 8.95E+05 |
| | 1.24E+06 | 1.02E+06 |
| DMSO Ovary only Well 6 | 1.80E+06 | 8.87E+05 |
| | 1.28E+06 | 8.59E+05 |
| | 1.81E+06 | 1.24E+06 |
| Bortezomib Ovary only Well 1 | 7.00E+05 | 3.54E+05 |
| | 7.05E+05 | 3.48E+05 |
| | 6.71E+05 | 3.57E+05 |
| Bortezomib Ovary only Well 2 | 5.35E+05 | 3.59E+05 |
| | 5.42E+05 | 3.44E+05 |
| | 4.65E+05 | 3.22E+05 |
| Bortezomib Ovary only Well 3 | 6.81E+05 | 3.75E+05 |
| | 7.03E+05 | 4.48E+05 |
| | 7.15E+05 | 4.60E+05 |
| Bortezomib Ovary only Well 4 | 6.55E+05 | 4.35E+05 |
| | 6.64E+05 | 5.10E+05 |
| | 6.39E+05 | 4.71E+05 |
| Bortezomib Ovary only Well 5 | 6.69E+05 | 4.91E+05 |
| | 6.59E+05 | 4.74E+05 |
| | 5.53E+05 | 4.02E+05 |
| Bortezomib Ovary only Well 6 | 7.40E+05 | 5.17E+05 |
| | 6.92E+05 | 5.10E+05 |
| | 7.81E+05 | 5.44E+05 |

**Table S6. Statistical analyses of *wsp* abundance from ovary tissues, in response to bortezomib treatment**

| Plate replicate # | Shapiro-wilk: p-values, interpretation | | | Levene's test: p-values, interpretation | | Outcome: analysis of different combinations | | | "n" required to satisfy $\beta < 0.05$ |
|---|---|---|---|---|---|---|---|---|---|
| | DMSO | Bortezomib | Normal distribution? | Overall value | Variance homogeneous? | Analysis required | Resulting p-value | Significantly different? | |
| 1 | 0.163 | 0.065 | both normal | 0.003 | no | Welch's T-test | <0.001 | yes | 5 |
| 2 | 0.97 | 0.125 | both normal | 0.124 | yes | T-test | <0.001 | yes | 3 |

* represent ovarian comparisons, all technical replicates, n = 18

**Table S7. Overview of bacterial taxa analyzed, based on the literature**

| Class | Genus | # species researched | # species pursued |
|---|---|---|---|
| *Alphaproteobacteria* | *Brucella* | 5 | 2 |
| *Alphaproteobacteria* | *Rickettsia* | 5 | 5 |
| *Alphaproteobacteria* | *Anaplasma* | 3 | 1 |
| *Alphaproteobacteria* | *Ehrlichia* | 3 | 3 |
| *Alphaproteobacteria* | *Wolbachia* | 1 | 1 |
| *Gammaroteobacteria* | *Legionella* | 8 | 1 |
| *Gammaroteobacteria* | *Francisella* | 3 | 1 |
| *Gammaroteobacteria* | *Salmonella* | 2 | 1 |
| *Gammaroteobacteria* | *Wigglesworthia* | 1 | 1 |
| *Gammaroteobacteria* | *Coxiella* | 1 | 1 |
| *Gammaroteobacteria* | *Pseudomonas* | 3 | 1 |
| *Actinobacteria* | *Mycobacterium* | 2 | 2 |
| *Bacilli* | *Listeria* | 5 | 1 |
| *Chlamydiae* | *Chlamydia* | 3 | 2 |
| *Clostridia* | *Candidatus* | 1 | 1 |
| *Mollicutes* | *Spiroplasma* | 1 | 1 |
| *Spirochaetales* | *Borrelia* | 5 | 1 |

**Table S8. Mechanisms reported to affect intracellular density of commonly studied microbes**

| Candidate host processes | Number of organisms represented | Number of classes represented | Name of the Organisms | References |
|---|---|---|---|---|
| Cell cycle modulation | 4 | 4 | *M. fortuitum*; *Listeria monocytogenes*; *Chlamydia caviae*; *Wolbachia* | Agaisse et al., 2005; Derre et al., 2007; Grobler et al., 2018 |
| Cytoskeleton-based transport | 6 | 4 | *Ehrlichia risticii; Ehrlichia canis*; *Listeria monocytogenes*; *Mycobacterium fortuitum*; *Chlamydia caviae*; *Wolbachia* | Rikihisa et al., 2004; Kuhbacher et al., 2015; Derre et al., 2007; Philips et al., 2005; Ferree et al., 2005, Serbus et al., 2011; Newton et al., 2015; Sheehan et al., 2016 |
| Ubiquitin-proteasome system | 3 | 1 | *Ehrlichia chaffeensis*; *Anaplasma phagocytophilum*; *Wolbachia;* | Zhu et al., 2017; Severo et al., 2013., White et al., 2017 |
| Mitochondrial/ Antioxidant | 4 | 4 | *Chlamydia cavia*; *Wolbachia*; *Francisella tularensis*; *Borrelia burgdorferi* | Derre et al., 2007; White et al., 2017 |
| Apoptotic pathway | 3 | 3 | *Brucella abortus*; *Legionella pneumophila*; *Mycobacteria* | Wei et al., 2015; Gross et al., 2000; Ashida et al., 2011 |
| Imd pathway | 5 | 4 | *Rickettsia typhi*; *Wigglesworthia glossinidia*; *Borrelia burgdorferi*; *Spiroplasma poulsonii*; *Pseudomonas aeruginosa* | Rennoll et al., 2018; Wang et al., 2009; Carroll et al., 2019; Herren & Lemaitre et al., 2011; Limmer et al., 2011 |
| GPCR Signaling | 5 | 2 | *Rickettsia conorii*; *Coxiella burnetii*; *Brucella abortus*; *Legionella pneumophila*; *Wolbachia*; *E. chaffeensis* | Czyz et al., 2014; Kim et al., 2012; White et al., 2017 |
| Phospholipase-related | 3 | 1 | *Ehrlichia chaffeensis*; *Ehrlichia canis*; *Rickettsia rickettsii*; | Lin et al., 2002; Levenhagen et al., 2012; Walker et al., 1983 |
| Calcium signaling | 7 | 2 | *E. risticii; Ehrlichia chaffeensis; Ehrlichia canis*; *Rickettsia conorii*; *Coxiella burnetii*; *Brucella abortus*; *Legionella pneumophila* | Rikihisa & Park., 1995; Lin et al., 2002; Levenhagen et al., 2012; Czyz et al., 2014; Kim et al., 2012; |
| Ras/mTor pathway | 13 | 7 | *Salmonella typhimorium; Chlamydia cavia; Listeria monocytogenes; Wolbachia; Franciella tularensis; Candidatus Arthromitus; Ehrlicia chaffeensis; Rickettsia australis; Mycobacterium tuberculosis*; *Pseudomonas aeruginosa; Anaplasma phagocytophilum; Coxiella burnetii ; Brucella suis* | Xiong et al., 2009; Czyz et al., 2014; Bagues et al., 2005; Luo et al., 2017; Birmingham et al., 2006; Derre et al., 2007; Steele et al., 2013; Yang et al., 2018; Niu et al., 2008; Niu et al., 2012; Lina et al., 2017; Bechelli et al., 2019; Gutierre et al., 2004; Wang et al., 2013; Yuan et al., 2012; Foldenauer et al., 2013 |
| Nitric oxide synthase | 4 | 2 | *Ehrlicia; Rickettsia prowazekii*; *Brucella abortus*; *Chlamydia cavia; Listeria monocytogenes* | Banerjee et al., 2000; Turco et al., 1998; Czyz et al., 2016; Zhang et al., 2012; Cole et al., 2012 |
| Jak-Stat signaling | 4 | 2 | *Anaplasma phagocytophilum*; *Wolbachia*; *Rickettsia conorii*; *Chlamydia trachomatis* | Liu et al., 2012; Grobler et al., 2018; Colonne et al., 2011; Lad et al., 2005; Zhang et al., 2018 |
| Wnt pathway | 4 | 3 | *Ehrlichia chaffeensis*; *Wolbachia*; *Chlamydia trachomatis*; *Mycobacterium tuberculosis* | Luo et al., 2016; Grobler et al., 2018; Kintner et al., 2017; Kessler et al., 2012; Rogan et al., 2019; Pean et al., 2017 |
| Kinase modulator | 7 | 4 | *Ehrlichia risticii & chaffeensis & canis*; *Mycobacterium tuberculosis*; *S. typhimurium*; *Coxiella burnetii*; *Brucella abortus*; *Pseudomonas aeruginosa* | Zhang & Rikihisa., 1997; Lin et al., 2002; Levenhagen et al., 2012; Kuijl et al., 2007; Czyz et al 2014; Czyz et al., 2016; Alhazmi et al., 2018 |

**Table S9. *wsp* absolute counts, following exposure of *D. melanogaster* to candidate drugs (non-hits from replicates 1-2)**

| Host cellular process | Drug name | Well # | *wsp* absolute count Plate replicate 1 | | | *wsp* absolute count Plate replicate 2 | | |
|---|---|---|---|---|---|---|---|---|
| | | | DMSO | Treatment | Rifampicin | DMSO | Treatment | Rifampicin |
| Cell cycle modulation | Sodium butyrate | Well 1 | 1.25E+07 | 9.54E+06 | 4.09E+06 | 9.56E+06 | 8.40E+06 | 3.26E+06 |
| | | | 1.08E+07 | 1.31E+07 | 3.60E+06 | 9.19E+06 | 8.90E+06 | 3.46E+06 |
| | | Well 2 | 1.14E+07 | 1.13E+07 | 3.02E+06 | 9.00E+06 | 9.38E+06 | 2.71E+06 |
| | | | 1.42E+07 | 8.87E+06 | 2.64E+06 | 9.00E+06 | 9.62E+06 | 3.27E+06 |
| | | Well 3 | 1.33E+07 | 1.09E+07 | 3.50E+06 | 8.21E+06 | 8.55E+06 | 3.06E+06 |
| | | | 1.31E+07 | 1.18E+07 | 3.28E+06 | 8.52E+06 | 9.58E+06 | 3.34E+06 |
| Cytoskeleton-based transport | Colchicine | Well 1 | 3.04E+06 | 2.65E+06 | 7.35E+05 | 3.92E+06 | 3.25E+06 | 1.37E+06 |
| | | | 3.47E+06 | 2.65E+06 | 7.27E+05 | 3.03E+06 | 3.68E+06 | 1.04E+06 |
| | | Well 2 | 2.92E+06 | 2.75E+06 | 6.89E+05 | 3.73E+06 | 3.93E+06 | 1.14E+06 |
| | | | 2.94E+06 | 2.36E+06 | 6.24E+05 | 3.54E+06 | 3.74E+06 | 1.86E+06 |
| | | Well 3 | 3.20E+06 | 2.97E+06 | 9.21E+05 | 3.46E+06 | 3.40E+06 | 1.13E+06 |
| | | | 3.49E+06 | 2.48E+06 | 8.64E+05 | 3.90E+06 | 2.75E+06 | 1.13E+06 |
| | Cytochalasin D | Well 1 | 1.19E+07 | 1.20E+07 | 2.49E+06 | 7.64E+06 | 6.08E+06 | 1.73E+06 |
| | | | 1.10E+07 | 8.98E+06 | 2.16E+06 | 6.44E+06 | 5.31E+06 | 1.66E+06 |
| | | Well 2 | 8.73E+06 | 9.07E+06 | 2.59E+06 | 5.98E+06 | 5.55E+06 | 2.10E+06 |
| | | | 8.70E+06 | 9.90E+06 | 2.56E+06 | 6.74E+06 | 5.32E+06 | 1.90E+06 |
| | | Well 3 | 1.05E+07 | 8.89E+06 | 2.43E+06 | 5.15E+06 | 6.79E+06 | 1.58E+06 |
| | | | 1.24E+07 | 8.06E+06 | 3.33E+06 | 6.31E+06 | 6.12E+06 | 1.54E+06 |
| Ubiquitin-proteasome system | Epoxomicin | Well 1 | 8.09E+06 | 7.86E+06 | 2.48E+06 | 7.88E+06 | 7.45E+06 | 2.10E+06 |
| | | | 7.34E+06 | 6.45E+06 | 2.54E+06 | 8.52E+06 | 5.90E+06 | 2.02E+06 |
| | | Well 2 | 7.66E+06 | 7.93E+06 | 2.48E+06 | 7.72E+06 | 5.77E+06 | 2.07E+06 |
| | | | 8.48E+06 | 6.80E+06 | 2.28E+06 | 6.89E+06 | 5.47E+06 | 1.90E+06 |
| | | Well 3 | 7.72E+06 | 6.02E+06 | 2.05E+06 | 8.52E+06 | 6.88E+06 | 2.00E+06 |
| | | | 7.34E+06 | 7.10E+06 | 2.04E+06 | 7.97E+06 | 7.33E+06 | 1.71E+06 |
| Mitochondrial/ Antioxidant | Butylated hydroxytoluene | Well 1 | 1.25E+07 | 7.37E+06 | 4.09E+06 | 9.56E+06 | 9.60E+06 | 3.26E+06 |
| | | | 1.08E+07 | 6.17E+06 | 3.60E+06 | 9.19E+06 | 8.73E+06 | 3.46E+06 |
| | | Well 2 | 1.14E+07 | 6.98E+06 | 3.02E+06 | 9.00E+06 | 8.09E+06 | 2.71E+06 |
| | | | 1.42E+07 | 7.78E+06 | 2.64E+06 | 9.00E+06 | 9.84E+06 | 3.27E+06 |
| | | Well 3 | 1.33E+07 | 7.45E+06 | 3.50E+06 | 8.21E+06 | 8.64E+06 | 3.06E+06 |
| | | | 1.31E+07 | 8.97E+06 | 3.28E+06 | 8.52E+06 | 9.23E+06 | 3.34E+06 |
| | Glutathione | Well 1 | 1.25E+07 | 1.65E+07 | 4.09E+06 | 9.56E+06 | 9.69E+06 | 3.26E+06 |
| | | | 1.08E+07 | 1.55E+07 | 3.60E+06 | 9.19E+06 | 8.95E+06 | 3.46E+06 |
| | | Well 2 | 1.14E+07 | 1.31E+07 | 3.02E+06 | 9.00E+06 | 1.01E+07 | 2.71E+06 |
| | | | 1.42E+07 | 1.45E+07 | 2.64E+06 | 9.00E+06 | 1.11E+07 | 3.27E+06 |
| | | Well 3 | 1.33E+07 | 1.20E+07 | 3.50E+06 | 8.21E+06 | 1.14E+07 | 3.06E+06 |
| | | | 1.31E+07 | 1.21E+07 | 3.28E+06 | 8.52E+06 | 1.07E+07 | 3.34E+06 |
| | Mitoblocker | Well 1 | 7.64E+06 | 8.34E+06 | 1.73E+06 | 6.54E+06 | 6.99E+06 | 2.38E+06 |
| | | | 6.44E+06 | 7.15E+06 | 1.66E+06 | 6.67E+06 | 6.80E+06 | 2.99E+06 |
| | | Well 2 | 5.98E+06 | 7.93E+06 | 2.10E+06 | 6.52E+06 | 6.66E+06 | 2.77E+06 |
| | | | 6.74E+06 | 8.03E+06 | 1.90E+06 | 6.72E+06 | 6.51E+06 | 2.39E+06 |
| | | Well 3 | 5.15E+06 | 7.59E+06 | 1.58E+06 | 6.42E+06 | 6.50E+06 | 2.87E+06 |
| | | | 6.31E+06 | 6.88E+06 | 1.54E+06 | 5.69E+06 | 6.78E+06 | 2.38E+06 |
| | Niclosamide | Well 1 | 5.94E+06 | 6.45E+06 | 1.94E+06 | 4.78E+06 | 5.28E+06 | 1.56E+06 |
| | | | 5.31E+06 | 6.45E+06 | 1.60E+06 | 4.64E+06 | 5.18E+06 | 1.50E+06 |
| | | Well 2 | 5.86E+06 | 6.81E+06 | 2.04E+06 | 5.04E+06 | 6.50E+06 | 1.49E+06 |
| | | | 5.37E+06 | 4.82E+06 | 2.28E+06 | 5.25E+06 | 5.74E+06 | 1.37E+06 |
| | | Well 3 | 5.10E+06 | 8.67E+06 | 1.45E+06 | 4.64E+06 | 5.69E+06 | 1.30E+06 |
| | | | 5.80E+06 | 7.25E+06 | 1.36E+06 | 4.73E+06 | 5.28E+06 | 1.22E+06 |
| | Resveratrol | Well 1 | 4.63E+06 | 5.44E+06 | 1.40E+06 | 5.74E+06 | 7.04E+06 | 2.55E+06 |
| | | | 5.36E+06 | 4.30E+06 | 1.37E+06 | 5.47E+06 | 6.47E+06 | 2.32E+06 |
| | | Well 2 | 4.64E+06 | 4.50E+06 | 1.23E+06 | 6.01E+06 | 1.09E+07 | 1.94E+06 |
| | | | 4.66E+06 | 5.51E+06 | 1.57E+06 | 6.00E+06 | 8.94E+06 | 2.00E+06 |
| | | Well 3 | 4.40E+06 | 5.85E+06 | 1.04E+06 | 6.32E+06 | 8.87E+06 | 2.02E+06 |
| | | | 4.39E+06 | 5.83E+06 | 1.20E+06 | 6.48E+06 | 9.42E+06 | 2.10E+06 |
| Apoptotic pathway | Apoptosis inhibitor | Well 1 | 8.09E+06 | 8.76E+06 | 2.48E+06 | 7.88E+06 | 7.03E+06 | 2.10E+06 |
| | | | 7.34E+06 | 9.26E+06 | 2.54E+06 | 8.52E+06 | 8.01E+06 | 2.02E+06 |
| | | Well 2 | 7.66E+06 | 9.93E+06 | 2.48E+06 | 7.72E+06 | 8.13E+06 | 2.07E+06 |
| | | | 8.48E+06 | 9.34E+06 | 2.28E+06 | 6.89E+06 | 7.13E+06 | 1.90E+06 |
| | | Well 3 | 7.72E+06 | 9.52E+06 | 2.05E+06 | 8.52E+06 | 7.52E+06 | 2.00E+06 |
| | | | 7.34E+06 | 7.47E+06 | 2.04E+06 | 7.97E+06 | 7.44E+06 | 1.71E+06 |
| | Caspase 8 inhibitor II | Well 1 | 5.82E+06 | 7.91E+06 | 1.68E+06 | 6.24E+06 | 8.95E+06 | 2.21E+06 |
| | | | 5.73E+06 | 7.47E+06 | 1.63E+06 | 7.21E+06 | 9.41E+06 | 2.16E+06 |
| | | Well 2 | 5.78E+06 | 6.27E+06 | 1.45E+06 | 7.25E+06 | 9.82E+06 | 2.37E+06 |
| | | | 5.61E+06 | 6.02E+06 | 1.40E+06 | 8.09E+06 | 8.68E+06 | 1.98E+06 |
| | | Well 3 | 5.71E+06 | 4.91E+06 | 1.53E+06 | 5.69E+06 | 6.67E+06 | 2.05E+06 |
| | | | 5.93E+06 | 6.39E+06 | 1.53E+06 | 5.77E+06 | 9.31E+06 | 1.76E+06 |
| | Caffeine | Well 1 | 5.40E+06 | 7.40E+06 | 2.38E+06 | 5.66E+06 | 5.56E+06 | 2.51E+06 |
| | | | 6.56E+06 | 7.01E+06 | 2.18E+06 | 5.11E+06 | 5.63E+06 | 2.35E+06 |
| | | Well 2 | 7.07E+06 | 6.46E+06 | 1.97E+06 | 6.26E+06 | 5.70E+06 | 2.34E+06 |

| | | | | | | | | |
|---|---|---|---|---|---|---|---|---|
| Gpcr Signaling | Caffeine | Well 2 | 6.38E+06 | 6.28E+06 | 2.02E+06 | 5.61E+06 | 5.77E+06 | 2.40E+06 |
| | | Well 3 | 5.86E+06<br>6.15E+06 | 6.70E+06<br>6.11E+06 | 1.93E+06<br>1.78E+06 | 6.52E+06<br>7.27E+06 | 5.84E+06<br>5.50E+06 | 2.45E+06<br>2.50E+06 |
| | SQ22536 | Well 1 | 7.94E+06<br>7.91E+06 | 1.08E+07<br>1.10E+07 | 3.94E+06<br>3.74E+06 | 7.03E+06<br>6.94E+06 | 7.24E+06<br>7.00E+06 | 2.14E+06<br>2.12E+06 |
| | | Well 2 | 8.61E+06<br>8.64E+06 | 1.13E+07<br>1.04E+07 | 3.13E+06<br>2.86E+06 | 6.83E+06<br>6.14E+06 | 7.99E+06<br>6.48E+06 | 2.31E+06<br>2.26E+06 |
| | | Well 3 | 7.72E+06<br>7.91E+06 | 1.02E+07<br>1.26E+07 | 2.94E+06<br>2.93E+06 | 6.12E+06<br>6.06E+06 | 6.40E+06<br>7.89E+06 | 2.43E+06<br>2.27E+06 |
| Calcium signaling | Verapamil | Well 1 | 3.04E+06<br>3.47E+06 | 3.62E+06<br>3.59E+06 | 7.35E+05<br>7.27E+05 | 3.92E+06<br>3.03E+06 | 3.00E+06<br>3.56E+06 | 1.37E+06<br>1.04E+06 |
| | | Well 2 | 2.92E+06<br>2.94E+06 | 3.80E+06<br>3.46E+06 | 6.89E+05<br>6.24E+05 | 3.73E+06<br>3.54E+06 | 3.53E+06<br>3.43E+06 | 1.14E+06<br>1.86E+06 |
| | | Well 3 | 3.20E+06<br>3.49E+06 | 3.80E+06<br>3.30E+06 | 9.21E+05<br>8.64E+05 | 3.46E+06<br>3.90E+06 | 4.05E+06<br>3.20E+06 | 1.13E+06<br>1.13E+06 |
| Ras/mTor pathway | Trametinib | Well 1 | 5.40E+06<br>6.56E+06 | 8.24E+06<br>7.72E+06 | 2.38E+06<br>2.18E+06 | 5.66E+06<br>5.11E+06 | 5.54E+06<br>4.62E+06 | 2.51E+06<br>2.35E+06 |
| | | Well 2 | 7.07E+06<br>6.38E+06 | 5.53E+06<br>5.36E+06 | 1.97E+06<br>2.02E+06 | 6.26E+06<br>5.61E+06 | 5.92E+06<br>6.77E+06 | 2.34E+06<br>2.40E+06 |
| | | Well 3 | 5.86E+06<br>6.15E+06 | 7.14E+06<br>6.62E+06 | 1.93E+06<br>1.78E+06 | 6.52E+06<br>7.27E+06 | 5.72E+06<br>5.72E+06 | 2.45E+06<br>2.50E+06 |
| | Wortmanin | Well 1 | 1.25E+07<br>1.08E+07 | 8.23E+06<br>8.58E+06 | 4.09E+06<br>3.60E+06 | 9.56E+06<br>9.19E+06 | 1.02E+07<br>1.02E+07 | 3.26E+06<br>3.46E+06 |
| | | Well 2 | 1.14E+07<br>1.42E+07 | 1.20E+07<br>8.36E+06 | 3.02E+06<br>2.64E+06 | 9.00E+06<br>9.00E+06 | 1.05E+07<br>1.03E+07 | 2.71E+06<br>3.27E+06 |
| | | Well 3 | 1.33E+07<br>1.31E+07 | 9.47E+06<br>1.21E+07 | 3.50E+06<br>3.28E+06 | 8.21E+06<br>8.52E+06 | 9.97E+06<br>1.05E+07 | 3.06E+06<br>3.34E+06 |
| Nitric oxide synthase | Methylxanthine | Well 1 | 5.94E+06<br>5.31E+06 | 7.84E+06<br>6.26E+06 | 1.94E+06<br>1.60E+06 | 4.78E+06<br>4.64E+06 | 4.59E+06<br>4.39E+06 | 1.56E+06<br>1.50E+06 |
| | | Well 2 | 5.86E+06<br>5.37E+06 | 7.13E+06<br>7.22E+06 | 2.04E+06<br>2.28E+06 | 5.04E+06<br>5.25E+06 | 4.51E+06<br>4.62E+06 | 1.49E+06<br>1.37E+06 |
| | | Well 3 | 5.10E+06<br>5.80E+06 | 5.55E+06<br>5.85E+06 | 1.45E+06<br>1.36E+06 | 4.64E+06<br>4.73E+06 | 4.31E+06<br>3.49E+06 | 1.30E+06<br>1.22E+06 |
| | Sildenafil citrate | Well 1 | 5.94E+06<br>5.31E+06 | 7.59E+06<br>8.20E+06 | 1.94E+06<br>1.60E+06 | 4.78E+06<br>4.64E+06 | 3.39E+06<br>3.91E+06 | 1.56E+06<br>1.50E+06 |
| | | Well 2 | 5.86E+06<br>5.37E+06 | 7.48E+06<br>6.85E+06 | 2.04E+06<br>2.28E+06 | 5.04E+06<br>5.25E+06 | 3.45E+06<br>3.47E+06 | 1.49E+06<br>1.37E+06 |
| | | Well 3 | 5.10E+06<br>5.80E+06 | 7.62E+06<br>6.77E+06 | 1.45E+06<br>1.36E+06 | 4.64E+06<br>4.73E+06 | 3.96E+06<br>3.97E+06 | 1.30E+06<br>1.22E+06 |
| Jak-Stat signaling | Ruxolitinib | Well 1 | 8.09E+06<br>7.34E+06 | 8.16E+06<br>7.95E+06 | 2.48E+06<br>2.54E+06 | 7.88E+06<br>8.52E+06 | 6.73E+06<br>5.88E+06 | 2.10E+06<br>2.02E+06 |
| | | Well 2 | 7.66E+06<br>8.48E+06 | 7.93E+06<br>9.11E+06 | 2.48E+06<br>2.28E+06 | 7.72E+06<br>6.89E+06 | 7.32E+06<br>7.88E+06 | 2.07E+06<br>1.90E+06 |
| | | Well 3 | 7.72E+06<br>7.34E+06 | 8.26E+06<br>9.44E+06 | 2.05E+06<br>2.04E+06 | 8.52E+06<br>7.97E+06 | 6.01E+06<br>6.80E+06 | 2.00E+06<br>1.71E+06 |
| | SH-4-54 | Well 1 | 5.82E+06<br>5.73E+06 | 6.50E+06<br>5.61E+06 | 1.68E+06<br>1.63E+06 | 6.24E+06<br>7.21E+06 | 8.67E+06<br>6.65E+06 | 2.21E+06<br>2.16E+06 |
| | | Well 2 | 5.78E+06<br>5.61E+06 | 6.20E+06<br>6.17E+06 | 1.45E+06<br>1.40E+06 | 7.25E+06<br>8.09E+06 | 5.04E+06<br>5.00E+06 | 2.37E+06<br>1.98E+06 |
| | | Well 3 | 5.71E+06<br>5.93E+06 | 6.67E+06<br>7.58E+06 | 1.53E+06<br>1.53E+06 | 5.69E+06<br>5.77E+06 | 5.34E+06<br>5.39E+06 | 2.05E+06<br>1.76E+06 |
| Wnt pathway | Ly090314 | Well 1 | 7.94E+06<br>7.91E+06 | 1.22E+07<br>1.04E+07 | 3.94E+06<br>3.74E+06 | 7.03E+06<br>6.94E+06 | 6.87E+06<br>6.79E+06 | 2.14E+06<br>2.12E+06 |
| | | Well 2 | 8.61E+06<br>8.64E+06 | 1.04E+07<br>1.13E+07 | 3.13E+06<br>2.86E+06 | 6.83E+06<br>6.14E+06 | 7.28E+06<br>6.84E+06 | 2.31E+06<br>2.26E+06 |
| | | Well 3 | 7.72E+06<br>7.91E+06 | 1.19E+07<br>1.13E+07 | 2.94E+06<br>2.93E+06 | 6.12E+06<br>6.06E+06 | 7.09E+06<br>7.09E+06 | 2.43E+06<br>2.27E+06 |

**Table S10. Statistical analyses of *wsp* absolute counts from non-hit drug treatments in *D. melanogaster* (from replicates 1-2)**

| Host cellular process | Drug name | Plate replicate # | Shapiro-wilk: p-values, interpretation | | | Levene's test: p-values, interpretation | | Outcome: analysis of different combinations | | | |
|---|---|---|---|---|---|---|---|---|---|---|---|
| | | | DMSO | Treatment | Normal distribution? | Overall value | Variance homogeneous? | Analysis required | Resulting p-value | Significantly different? | Abundance relative to control |
| Cell cycle modulation | Sodium butyrate | 1 | 0.848 | 0.926 | both normal | 0.733 | yes | T-test | 0.072 | no | n/a |
| | | 2 | 0.837 | 0.293 | both normal | 0.465 | yes | T-test | 0.6 | no | n/a |
| Cytoskeleton-based transport | Colchicine | 1 | 0.194 | 0.911 | both normal | 0.428 | yes | T-test | 0.003 | yes | lower |
| | | 2 | 0.443 | 0.683 | both normal | 0.587 | yes | T-test | 0.544 | no | n/a |
| | Cytochalasin D | 1 | 0.379 | 0.180 | both normal | 0.512 | yes | T-test | 0.241 | no | n/a |
| | | 2 | 0.956 | 0.345 | both normal | 0.714 | yes | T-test | 0.239 | no | n/a |
| Ubiquitin-proteasome system | Epoxomicin | 1 | 0.427 | 0.619 | both normal | 0.191 | yes | T-test | 0.066 | no | no |
| | | 2 | 0.366 | 0.257 | both normal | 0.115 | yes | T-test | 0.007 | yes | lower |
| Mitochondrial/ Antioxidant | Butylated hydroxytoluene | 1 | 0.848 | 0.847 | both normal | 0.367 | yes | T-test | <0.001 | yes | lower |
| | | 2 | 0.837 | 0.854 | both normal | 0.295 | yes | T-test | 0.751 | no | n/a |
| | Glutathione | 1 | 0.848 | 0.501 | both normal | 0.212 | yes | T-test | 0.157 | no | n/a |
| | | 2 | 0.837 | 0.864 | both normal | 0.100 | yes | T-test | 0.008 | yes | higher |
| | Mitoblocker | 1 | 0.956 | 0.792 | both normal | 0.657 | yes | T-test | 0.01 | yes | higher |
| | | 2 | 0.025 | 0.589 | one is non-normal | 0.636 | yes | Mann-Whitney U test | 0.24 | no | n/a |
| | Niclosamide | 1 | 0.323 | 0.751 | both normal | 0.193 | yes | T-test | 0.051 | no | n/a |
| | | 2 | 0.167 | 0.141 | both normal | 0.247 | yes | T-test | 0.007 | yes | higher |
| | Resveratrol | 1 | 0.033 | 0.122 | one is non-normal | 0.243 | yes | Mann-Whitney U test | 0.24 | no | n/a |
| | | 2 | 0.895 | 0.706 | both normal | 0.102 | yes | T-test | 0.003 | yes | higher |
| Apoptotic pathway | Apoptosis inhibitor | 1 | 0.427 | 0.255 | both normal | 0.477 | yes | T-test | 0.009 | yes | higher |
| | | 2 | 0.366 | 0.486 | both normal | 0.698 | yes | T-test | 0.252 | no | n/a |
| | Caspase 8 inhibitor II | 1 | 0.987 | 0.798 | both normal | 0.062 | yes | T-test | 0.127 | no | n/a |
| | | 2 | 0.438 | 0.087 | both normal | 0.802 | yes | T-test | 0.006 | yes | higher |
| Gpcr Signaling | Caffeine | 1 | 1 | 0.832 | both normal | 0.750 | yes | T-test | 0.198 | no | n/a |
| | | 2 | 0.869 | 0.940 | both normal | 0.027 | no | Welch's T-test | 0.258 | no | n/a |
| | SQ22536 | 1 | 0.048 | 0.344 | one is non-normal | 0.302 | yes | Mann-Whitney U test | 0.002 | yes | higher |
| | | 2 | 0.056 | 0.393 | both normal | 0.403 | yes | T-test | 0.081 | no | n/a |
| Calcium signaling | Verapamil | 1 | 0.194 | 0.556 | both normal | 0.371 | yes | T-test | 0.01 | yes | lower |
| | | 2 | 0.443 | 0.806 | both normal | 0.989 | yes | T-test | 0.515 | no | n/a |
| Ras/mTor pathway | Trametinib | 1 | 1 | 0.622 | both normal | 0.100 | yes | T-test | 0.339 | no | n/a |
| | | 2 | 0.869 | 0.578 | both normal | 0.493 | yes | T-test | 0.419 | no | n/a |
| | Wortmanin | 1 | 0.848 | 0.049 | one is non-normal | 0.503 | yes | Mann-Whitney U test | 0.026 | yes | lower |
| | | 2 | 0.837 | 0.458 | both normal | 0.241 | yes | T-test | <0.001 | yes | higher |
| Nitric oxide synthase | Methylxanthine | 1 | 0.323 | 0.669 | both normal | 0.021 | no | Welch's T-test | 0.03 | yes | higher |
| | | 2 | 0.167 | 0.018 | one is non-normal | 0.639 | yes | Mann-Whitney U test | 0.002 | yes | lower |
| | Sildenafil citrate | 1 | 0.323 | 0.494 | both normal | 0.606 | yes | T-test | <0.001 | yes | higher |
| | | 2 | 0.167 | 0.034 | one is non-normal | 0.346 | yes | Mann-Whitney U test | 0.002 | yes | lower |
| Jak-Stat signaling | Ruxolitinib | 1 | 0.427 | 0.108 | both normal | 0.569 | yes | T-test | 0.052 | no | n/a |
| | | 2 | 0.366 | 0.723 | both normal | 0.568 | yes | T-test | 0.016 | yes | lower |
| | SH-4-54 | 1 | 0.987 | 0.701 | both normal | 0.086 | yes | T-test | 0.03 | yes | higher |
| | | 2 | 0.438 | 0.032 | one is non-normal | 0.887 | yes | Mann-Whitney U test | 0.18 | no | n/a |
| Wnt pathway | Ly090314 | 1 | 0.048 | 0.361 | one is non-normal | 0.255 | yes | Mann-Whitney U test | 0.002 | yes | higher |
| | | 2 | 0.056 | 0.404 | both normal | 0.000 | no | Welch's T-test | 0.054 | no | n/a |

* represent whole body comparisons, all technical replicates, n = 6

**Table S11.** ***wsp*** **absolute counts, following exposure of** ***D. melanogaster*** **flies to candidate drugs in 3 total plate replicates**

| Host cellular process | Drug name | Well # | *wsp* absolute count Plate replicate 1 | | | *wsp* absolute count Plate replicate 2 | | | *wsp* absolute count Plate replicate 3 | | |
|---|---|---|---|---|---|---|---|---|---|---|---|
| | | | DMSO | Treatment | Rifampicin | DMSO | Treatment | Rifampicin | DMSO | Treatment | Rifampicin |
| Cell cycle modulation | Flavopiridol | Well 1 | 7.94E+06 | 1.28E+07 | 3.94E+06 | 7.03E+06 | 6.93E+06 | 2.14E+06 | 2.20E+07 | 1.85E+07 | 3.13E+06 |
| | | | 7.91E+06 | 1.18E+07 | 3.74E+06 | 6.94E+06 | 7.05E+06 | 2.12E+06 | 1.85E+07 | 1.79E+07 | 3.06E+06 |
| | | Well 2 | 8.61E+06 | 1.46E+07 | 3.13E+06 | 6.83E+06 | 9.50E+06 | 2.31E+06 | 1.72E+07 | 1.04E+07 | 2.97E+06 |
| | | | 8.64E+06 | 1.33E+07 | 2.86E+06 | 6.14E+06 | 7.92E+06 | 2.26E+06 | 1.63E+07 | 1.24E+07 | 2.49E+06 |
| | | Well 3 | 7.72E+06 | 1.06E+07 | 2.94E+06 | 6.12E+06 | 7.96E+06 | 2.43E+06 | 2.41E+07 | 1.83E+07 | 2.01E+06 |
| | | | 7.91E+06 | 1.06E+07 | 2.93E+06 | 6.06E+06 | 7.39E+06 | 2.27E+06 | 1.58E+07 | 1.59E+07 | 1.53E+06 |
| | Roscovitine | Well 1 | 5.82E+06 | 7.83E+06 | 1.68E+06 | 6.24E+06 | 8.86E+06 | 2.21E+06 | 5.23E+06 | 6.73E+06 | 1.50E+06 |
| | | | 5.73E+06 | 6.45E+06 | 1.63E+06 | 7.21E+06 | 8.21E+06 | 2.16E+06 | 5.29E+06 | 5.74E+06 | 1.51E+06 |
| | | Well 2 | 5.78E+06 | 6.69E+06 | 1.45E+06 | 7.25E+06 | 7.77E+06 | 2.37E+06 | 5.68E+06 | 5.75E+06 | 1.52E+06 |
| | | | 5.61E+06 | 6.26E+06 | 1.40E+06 | 8.09E+06 | 7.98E+06 | 1.98E+06 | 5.57E+06 | 5.82E+06 | 1.52E+06 |
| | | Well 3 | 5.71E+06 | 7.41E+06 | 1.53E+06 | 5.69E+06 | 8.29E+06 | 2.05E+06 | 5.39E+06 | 6.11E+06 | 1.53E+06 |
| | | | 5.93E+06 | 6.24E+06 | 1.53E+06 | 5.77E+06 | 8.96E+06 | 1.76E+06 | 5.80E+06 | 5.81E+06 | 1.54E+06 |
| Cytoskeleton-based transport | Taxol | Well 1 | 3.04E+06 | 3.83E+06 | 7.35E+05 | 3.92E+06 | 4.05E+06 | 1.37E+06 | 1.19E+07 | 8.91E+06 | 2.49E+06 |
| | | | 3.47E+06 | 3.50E+06 | 7.27E+05 | 3.03E+06 | 4.08E+06 | 1.04E+06 | 1.10E+07 | 8.14E+06 | 2.16E+06 |
| | | Well 2 | 2.92E+06 | 3.45E+06 | 6.89E+05 | 3.73E+06 | 4.33E+06 | 1.14E+06 | 8.73E+06 | 7.61E+06 | 2.59E+06 |
| | | | 2.94E+06 | 3.31E+06 | 6.24E+05 | 3.54E+06 | 3.91E+06 | 1.86E+06 | 8.70E+06 | 7.93E+06 | 2.56E+06 |
| | | Well 3 | 3.20E+06 | 3.76E+06 | 9.21E+05 | 3.46E+06 | 4.29E+06 | 1.13E+06 | 1.05E+07 | 8.92E+06 | 2.43E+06 |
| | | | 3.49E+06 | 3.21E+06 | 8.64E+05 | 3.90E+06 | 4.22E+06 | 1.13E+06 | 1.24E+07 | 9.00E+06 | 3.33E+06 |
| Ubiquitin-proteasome system | Bortezomib | Well 1 | 3.04E+06 | 2.58E+06 | 7.35E+05 | 3.92E+06 | 2.45E+06 | 1.37E+06 | 1.19E+07 | 7.09E+06 | 2.49E+06 |
| | | | 3.47E+06 | 2.22E+06 | 7.27E+05 | 3.03E+06 | 2.18E+06 | 1.04E+06 | 1.10E+07 | 4.83E+06 | 2.16E+06 |
| | | Well 2 | 2.92E+06 | 2.21E+06 | 6.89E+05 | 3.73E+06 | 2.78E+06 | 1.14E+06 | 8.73E+06 | 5.13E+06 | 2.59E+06 |
| | | | 2.94E+06 | 2.34E+06 | 6.24E+05 | 3.54E+06 | 2.27E+06 | 1.86E+06 | 8.70E+06 | 4.62E+06 | 2.56E+06 |
| | | Well 3 | 3.20E+06 | 1.94E+06 | 9.21E+05 | 3.46E+06 | 2.55E+06 | 1.13E+06 | 1.05E+07 | 5.20E+06 | 2.43E+06 |
| | | | 3.49E+06 | 1.67E+06 | 8.64E+05 | 3.90E+06 | 2.43E+06 | 1.13E+06 | 1.24E+07 | 5.88E+06 | 3.33E+06 |
| Apoptotic pathway | ABT-199 | Well 1 | 5.40E+06 | 8.73E+06 | 2.38E+06 | 5.66E+06 | 6.24E+06 | 2.51E+06 | 2.20E+07 | 2.66E+07 | 3.13E+06 |
| | | | 6.56E+06 | 9.20E+06 | 2.18E+06 | 5.11E+06 | 8.82E+06 | 2.35E+06 | 1.85E+07 | 1.75E+07 | 3.06E+06 |
| | | Well 2 | 7.07E+06 | 9.99E+06 | 1.97E+06 | 6.26E+06 | 6.79E+06 | 2.34E+06 | 1.72E+07 | 2.98E+07 | 2.97E+06 |
| | | | 6.38E+06 | 9.56E+06 | 2.02E+06 | 5.61E+06 | 7.84E+06 | 2.40E+06 | 1.63E+07 | 2.43E+07 | 2.49E+06 |
| | | Well 3 | 5.86E+06 | 8.37E+06 | 1.93E+06 | 6.52E+06 | 8.11E+06 | 2.45E+06 | 2.41E+07 | 1.85E+07 | 2.01E+06 |
| | | | 6.15E+06 | 7.13E+06 | 1.78E+06 | 7.27E+06 | 8.39E+06 | 2.50E+06 | 1.58E+07 | 2.26E+07 | 1.53E+06 |
| Imd pathway | CAY10512 | Well 1 | 4.40E+06 | 5.26E+06 | 1.35E+06 | 6.96E+06 | 8.32E+06 | 2.45E+06 | 7.64E+06 | 7.33E+06 | 1.73E+06 |
| | | | 4.69E+06 | 5.06E+06 | 1.36E+06 | 7.76E+06 | 9.91E+06 | 3.04E+06 | 6.44E+06 | 7.48E+06 | 1.66E+06 |
| | | Well 2 | 4.26E+06 | 4.78E+06 | 1.22E+06 | 7.76E+06 | 1.13E+07 | 2.61E+06 | 5.98E+06 | 7.24E+06 | 2.10E+06 |
| | | | 4.67E+06 | 4.92E+06 | 1.22E+06 | 7.67E+06 | 1.32E+07 | 2.02E+06 | 6.74E+06 | 6.74E+06 | 1.90E+06 |
| | | Well 3 | 5.10E+06 | 5.53E+06 | 1.14E+06 | 6.27E+06 | 1.07E+07 | 2.77E+06 | 5.15E+06 | 7.02E+06 | 1.58E+06 |
| | | | 4.88E+06 | 5.55E+06 | 1.13E+06 | 5.76E+06 | 1.07E+07 | 2.29E+06 | 6.31E+06 | 7.77E+06 | 1.54E+06 |
| | SC76741 | Well 1 | 8.09E+06 | 9.54E+06 | 2.48E+06 | 7.88E+06 | 8.70E+06 | 2.10E+06 | 5.23E+06 | 7.47E+06 | 1.50E+06 |
| | | | 7.34E+06 | 7.15E+06 | 2.54E+06 | 8.52E+06 | 8.93E+06 | 2.02E+06 | 5.29E+06 | 7.65E+06 | 1.51E+06 |
| | | Well 2 | 7.66E+06 | 8.92E+06 | 2.48E+06 | 7.72E+06 | 8.52E+06 | 2.07E+06 | 5.68E+06 | 8.66E+06 | 1.52E+06 |
| | | | 8.48E+06 | 9.02E+06 | 2.28E+06 | 6.89E+06 | 8.49E+06 | 1.90E+06 | 5.57E+06 | 6.91E+06 | 1.52E+06 |
| | | Well 3 | 7.72E+06 | 1.10E+07 | 2.05E+06 | 8.52E+06 | 8.94E+06 | 2.00E+06 | 5.39E+06 | 7.50E+06 | 1.53E+06 |
| | | | 7.34E+06 | 1.08E+07 | 2.04E+06 | 7.97E+06 | 8.35E+06 | 1.71E+06 | 5.80E+06 | 7.43E+06 | 1.54E+06 |
| Phospholipase-related | U73122 | Well 1 | 3.04E+06 | 3.91E+06 | 7.35E+05 | 3.92E+06 | 4.02E+06 | 1.37E+06 | 5.34E+06 | 4.11E+06 | 1.16E+06 |
| | | | 3.47E+06 | 3.71E+06 | 7.27E+05 | 3.03E+06 | 4.40E+06 | 1.04E+06 | 5.64E+06 | 3.56E+06 | 1.21E+06 |
| | | Well 2 | 2.92E+06 | 3.46E+06 | 6.89E+05 | 3.73E+06 | 5.32E+06 | 1.14E+06 | 4.75E+06 | 4.54E+06 | 7.77E+05 |
| | | | 2.94E+06 | 3.40E+06 | 6.24E+05 | 3.54E+06 | 5.66E+06 | 1.86E+06 | 4.49E+06 | 4.43E+06 | 6.19E+05 |
| | | Well 3 | 3.20E+06 | 3.52E+06 | 9.21E+05 | 3.46E+06 | 3.83E+06 | 1.13E+06 | 4.49E+06 | 3.88E+06 | 1.40E+06 |
| | | | 3.49E+06 | 4.39E+06 | 8.64E+05 | 3.90E+06 | 4.67E+06 | 1.13E+06 | 5.54E+06 | 3.77E+06 | 1.40E+06 |
| | M-3M3FBS | Well 1 | 4.40E+06 | 6.77E+06 | 1.35E+06 | 6.96E+06 | 1.14E+07 | 2.45E+06 | 7.17E+06 | 1.18E+07 | 2.14E+06 |
| | | | 4.69E+06 | 6.32E+06 | 1.36E+06 | 7.76E+06 | 8.07E+06 | 3.04E+06 | 5.10E+06 | 1.03E+07 | 2.50E+06 |
| | | Well 2 | 4.26E+06 | 6.43E+06 | 1.22E+06 | 7.76E+06 | 1.03E+07 | 2.61E+06 | 7.93E+06 | 1.83E+07 | 1.97E+06 |
| | | | 4.67E+06 | 5.79E+06 | 1.22E+06 | 7.67E+06 | 9.69E+06 | 2.02E+06 | 5.42E+06 | 8.15E+06 | 2.07E+06 |
| | | Well 3 | 5.10E+06 | 4.98E+06 | 1.14E+06 | 6.27E+06 | 8.98E+06 | 2.77E+06 | 7.36E+06 | 8.97E+06 | 1.97E+06 |
| | | | 4.88E+06 | 5.04E+06 | 1.13E+06 | 5.76E+06 | 8.84E+06 | 2.29E+06 | 6.35E+06 | 8.48E+06 | 1.55E+06 |
| Calcium signaling | Nicardipine HCl | Well 1 | 8.09E+06 | 9.80E+06 | 2.48E+06 | 7.88E+06 | 1.10E+07 | 2.10E+06 | 5.23E+06 | 8.10E+06 | 1.50E+06 |
| | | | 7.34E+06 | 8.29E+06 | 2.54E+06 | 8.52E+06 | 9.90E+06 | 2.02E+06 | 5.29E+06 | 7.60E+06 | 1.51E+06 |
| | | Well 2 | 7.66E+06 | 8.79E+06 | 2.48E+06 | 7.72E+06 | 8.67E+06 | 2.07E+06 | 5.68E+06 | 7.10E+06 | 1.52E+06 |
| | | | 8.48E+06 | 8.11E+06 | 2.28E+06 | 6.89E+06 | 8.80E+06 | 1.90E+06 | 5.57E+06 | 6.60E+06 | 1.52E+06 |
| | | Well 3 | 7.72E+06 | 8.18E+06 | 2.05E+06 | 8.52E+06 | 8.68E+06 | 2.00E+06 | 5.39E+06 | 6.79E+06 | 1.53E+06 |
| | | | 7.34E+06 | 9.67E+06 | 2.04E+06 | 7.97E+06 | 8.85E+06 | 1.71E+06 | 5.80E+06 | 6.59E+06 | 1.54E+06 |
| Ras/mTor pathway | Erlotinib HCl | Well 1 | 5.40E+06 | 8.09E+06 | 2.38E+06 | 5.66E+06 | 8.91E+06 | 2.51E+06 | 5.23E+06 | 6.60E+06 | 1.50E+06 |
| | | | 6.56E+06 | 7.20E+06 | 2.18E+06 | 5.11E+06 | 7.33E+06 | 2.35E+06 | 5.29E+06 | 6.94E+06 | 1.51E+06 |
| | | Well 2 | 7.07E+06 | 8.06E+06 | 1.97E+06 | 6.26E+06 | 7.90E+06 | 2.34E+06 | 5.68E+06 | 8.01E+06 | 1.52E+06 |
| | | | 6.38E+06 | 7.89E+06 | 2.02E+06 | 5.61E+06 | 8.07E+06 | 2.40E+06 | 5.57E+06 | 8.20E+06 | 1.52E+06 |
| | | Well 3 | 5.86E+06 | 8.37E+06 | 1.93E+06 | 6.52E+06 | 7.64E+06 | 2.45E+06 | 5.39E+06 | 8.91E+06 | 1.53E+06 |
| | | | 6.15E+06 | 7.87E+06 | 1.78E+06 | 7.27E+06 | 6.84E+06 | 2.50E+06 | 5.80E+06 | 9.60E+06 | 1.54E+06 |
| | Rapamycin | Well 1 | 5.82E+06 | 8.04E+06 | 1.68E+06 | 6.24E+06 | 9.66E+06 | 2.21E+06 | 6.96E+06 | 1.14E+07 | 2.45E+06 |
| | | | 5.73E+06 | 8.18E+06 | 1.63E+06 | 7.21E+06 | 8.81E+06 | 2.16E+06 | 7.76E+06 | 1.33E+07 | 3.04E+06 |
| | | Well 2 | 5.78E+06 | 8.63E+06 | 1.45E+06 | 7.25E+06 | 8.29E+06 | 2.37E+06 | 7.76E+06 | 9.89E+06 | 2.61E+06 |
| | | | 5.61E+06 | 9.07E+06 | 1.40E+06 | 8.09E+06 | 1.28E+07 | 1.98E+06 | 7.67E+06 | 1.02E+07 | 2.02E+06 |
| | | Well 3 | 5.71E+06 | 9.52E+06 | 1.53E+06 | 5.69E+06 | 1.04E+07 | 2.05E+06 | 6.27E+06 | 1.06E+07 | 2.77E+06 |
| | | | 5.93E+06 | 8.39E+06 | 1.53E+06 | 5.77E+06 | 1.08E+07 | 1.76E+06 | 5.76E+06 | 1.09E+07 | 2.29E+06 |
| Nitric oxide synthase | L-NAME | Well 1 | 4.63E+06 | 6.97E+06 | 1.40E+06 | 5.74E+06 | 7.95E+06 | 2.55E+06 | 5.23E+06 | 7.32E+06 | 1.50E+06 |
| | | | 5.36E+06 | 5.61E+06 | 1.37E+06 | 5.47E+06 | 7.05E+06 | 2.32E+06 | 5.29E+06 | 7.60E+06 | 1.51E+06 |
| | | Well 2 | 4.64E+06 | 5.85E+06 | 1.23E+06 | 6.01E+06 | 6.90E+06 | 1.94E+06 | 5.68E+06 | 6.73E+06 | 1.52E+06 |
| | | | 4.66E+06 | 6.03E+06 | 1.57E+06 | 6.00E+06 | 6.88E+06 | 2.00E+06 | 5.57E+06 | 6.37E+06 | 1.52E+06 |
| | | Well 3 | 4.40E+06 | 5.80E+06 | 1.04E+06 | 6.32E+06 | 6.86E+06 | 2.02E+06 | 5.39E+06 | 7.11E+06 | 1.53E+06 |
| | | | 4.39E+06 | 5.46E+06 | 1.20E+06 | 6.48E+06 | 7.83E+06 | 2.10E+06 | 5.80E+06 | 7.56E+06 | 1.54E+06 |
| Wnt pathway | IWR-1 | Well 1 | 5.82E+06 | 6.90E+06 | 1.68E+06 | 6.24E+06 | 8.95E+06 | 2.21E+06 | 6.96E+06 | 8.99E+06 | 2.45E+06 |
| | | | 5.73E+06 | 7.14E+06 | 1.63E+06 | 7.21E+06 | 1.02E+07 | 2.16E+06 | 7.76E+06 | 1.04E+07 | 3.04E+06 |
| | | Well 2 | 5.78E+06 | 8.55E+06 | 1.45E+06 | 7.25E+06 | 1.06E+07 | 2.37E+06 | 7.76E+06 | 9.69E+06 | 2.61E+06 |
| | | | 5.61E+06 | 7.79E+06 | 1.40E+06 | 8.09E+06 | 1.05E+07 | 1.98E+06 | 7.67E+06 | 1.05E+07 | 2.02E+06 |
| | | Well 3 | 5.71E+06 | 7.44E+06 | 1.53E+06 | 5.69E+06 | 1.17E+07 | 2.05E+06 | 6.27E+06 | 9.91E+06 | 2.77E+06 |
| | | | 5.93E+06 | 6.00E+06 | 1.53E+06 | 5.77E+06 | 1.01E+07 | 1.76E+06 | 5.76E+06 | 8.97E+06 | 2.29E+06 |
| Kinase modulator | Pyrvinium pamoate | Well 1 | 4.63E+06 | 9.40E+06 | 1.40E+06 | 5.74E+06 | 7.16E+06 | 2.55E+06 | 1.19E+07 | 7.86E+06 | 2.49E+06 |
| | | | 5.36E+06 | 8.52E+06 | 1.37E+06 | 5.47E+06 | 7.34E+06 | 2.32E+06 | 1.10E+07 | 8.76E+06 | 2.16E+06 |
| | | Well 2 | 4.64E+06 | 7.48E+06 | 1.23E+06 | 6.01E+06 | 6.84E+06 | 1.94E+06 | 8.73E+06 | 1.10E+07 | 2.59E+06 |
| | | | 4.66E+06 | 7.11E+06 | 1.57E+06 | 6.00E+06 | 7.46E+06 | 2.00E+06 | 8.70E+06 | 9.44E+06 | 2.56E+06 |
| | | Well 3 | 4.40E+06 | 7.46E+06 | 1.04E+06 | 6.32E+06 | 7.30E+06 | 2.02E+06 | 1.05E+07 | 1.10E+07 | 2.43E+06 |
| | | | 4.39E+06 | 7.81E+06 | 1.20E+06 | 6.48E+06 | 7.34E+06 | 2.10E+06 | 1.24E+07 | 8.93E+06 | 3.33E+06 |
| | Staurosprine | Well 1 | 7.94E+06 | 1.18E+07 | 3.94E+06 | 7.03E+06 | 7.51E+06 | 2.14E+06 | 6.96E+06 | 9.55E+06 | 2.45E+06 |
| | | | 7.91E+06 | 1.20E+07 | 3.74E+06 | 6.94E+06 | 7.44E+06 | 2.12E+06 | 7.76E+06 | 1.15E+07 | 3.04E+06 |
| | | Well 2 | 8.61E+06 | 1.26E+07 | 3.13E+06 | 6.83E+06 | 7.46E+06 | 2.31E+06 | 7.76E+06 | 1.15E+07 | 2.61E+06 |
| | | | 8.64E+06 | 1.30E+07 | 2.86E+06 | 6.14E+06 | 6.98E+06 | 2.26E+06 | 7.67E+06 | 1.16E+07 | 2.02E+06 |
| | | Well 3 | 7.72E+06 | 1.16E+07 | 2.94E+06 | 6.12E+06 | 8.01E+06 | 2.43E+06 | 6.27E+06 | 1.13E+07 | 2.77E+06 |
| | | | 7.91E+06 | 9.57E+06 | 2.93E+06 | 6.06E+06 | 6.91E+06 | 2.27E+06 | 5.76E+06 | 1.18E+07 | 2.29E+06 |

**Table S12. Statistical analyses of *wsp* absolute counts from candidate drugs tested in 3 total plate replicates in *D. melanogaster***

| Host cellular process | Drug name | Plate replicate # | Shapiro-wilk: p-values, interpretation | | | Levene's test: p-values, interpretation | | Outcome: analysis of different combinations | | | | Interpreted as a hit? |
|---|---|---|---|---|---|---|---|---|---|---|---|---|
| | | | DMSO | Treatment | Normal distribution? | Overall value | Variance homogeneous? | Analysis required | Resulting p-value | Significantly different? | Abundance relative to control | |
| Cell cycle modulation | Flavopiridol | 1 | 0.048 | 0.577 | one is non-normal | 0.018 | no | T-test w/bootstrap | 0.001 | yes | higher | no |
| | | 2 | 0.056 | 0.207 | both normal | 0.354 | yes | T-test | 0.014 | yes | higher | |
| | | 3 | 0.305 | 0.156 | both normal | 0.931 | yes | T-test | 0.111 | no | n/a | |
| | Roscovitine | 1 | 0.987 | 0.188 | both normal | 0.076 | yes | T-test | 0.003 | yes | higher | yes |
| | | 2 | 0.438 | 0.509 | both normal | 0.041 | no | Welch's T-test | 0.004 | yes | higher | |
| | | 3 | 0.696 | 0.014 | one is non-normal | 0.814 | yes | Mann-Whitney U | 0.009 | yes | higher | |
| Cytoskeleton-based transport | Taxol | 1 | 0.194 | 0.647 | both normal | 0.768 | yes | T-test | 0.043 | yes | higher | no |
| | | 2 | 0.443 | 0.707 | both normal | 0.194 | yes | T-test | 0.005 | yes | higher | |
| | | 3 | 0.379 | 0.167 | both normal | 0.092 | yes | T-test | 0.011 | yes | lower | |
| Ubiquitin-proteasome system | Bortezomib | 1 | 0.194 | 0.883 | both normal | 0.924 | yes | T-test | <0.001 | yes | lower | yes |
| | | 2 | 0.443 | 0.883 | both normal | 0.305 | yes | T-test | <0.001 | yes | lower | |
| | | 3 | 0.379 | 0.218 | both normal | 0.186 | yes | T-test | <0.001 | yes | lower | |
| Apoptotic pathway | ABT-199 | 1 | 1.000 | 0.792 | both normal | 0.290 | yes | T-test | <0.001 | yes | higher | no |
| | | 2 | 0.869 | 0.593 | both normal | 0.684 | yes | T-test | 0.010 | yes | higher | |
| | | 3 | 0.305 | 0.816 | both normal | 0.422 | yes | T-test | 0.103 | no | n/a | |
| Imd pathway | CAY10512 | 1 | 0.914 | 0.515 | both normal | 0.681 | yes | T-test | 0.017 | yes | higher | yes |
| | | 2 | 0.159 | 0.857 | both normal | 0.515 | yes | T-test | 0.001 | yes | higher | |
| | | 3 | 0.956 | 0.998 | both normal | 0.254 | yes | T-test | 0.036 | yes | higher | |
| | SC76741 | 1 | 0.427 | 0.594 | both normal | 0.095 | yes | T-test | 0.022 | yes | higher | yes |
| | | 2 | 0.366 | 0.404 | both normal | 0.293 | yes | T-test | 0.020 | yes | higher | |
| | | 3 | 0.696 | 0.168 | both normal | 0.489 | yes | T-test | <0.001 | yes | higher | |
| Phospholipase-related | U73122 | 1 | 0.194 | 0.249 | both normal | 0.615 | yes | T-test | 0.013 | yes | higher | no |
| | | 2 | 0.443 | 0.676 | both normal | 0.118 | yes | T-test | 0.009 | yes | higher | |
| | | 3 | 0.156 | 0.736 | both normal | 0.134 | yes | T-test | 0.004 | yes | lower | |
| | M-3M3FBS | 1 | 0.914 | 0.355 | both normal | 0.050 | yes | T-test | 0.004 | yes | higher | yes |
| | | 2 | 0.159 | 0.909 | both normal | 0.540 | yes | T-test | 0.002 | yes | higher | |
| | | 3 | 0.602 | 0.044 | one is non-normal | 0.281 | yes | Mann-Whitney U | 0.002 | yes | higher | |
| Calcium signaling | Nicardipine HCl | 1 | 0.427 | 0.116 | both normal | 0.224 | yes | T-test | 0.016 | yes | higher | yes |
| | | 2 | 0.366 | 0.025 | both normal | 0.658 | yes | Mann-Whitney U | 0.002 | yes | higher | |
| | | 3 | 0.696 | 0.287 | both normal | 0.135 | yes | T-test | <0.001 | yes | higher | |
| Ras/mTor pathway | Erlotinib HCl | 1 | 1.000 | 0.316 | both normal | 0.350 | yes | T-test | <0.001 | yes | higher | yes |
| | | 2 | 0.869 | 0.965 | both normal | 0.685 | yes | T-test | 0.002 | yes | higher | |
| | | 3 | 0.696 | 0.823 | both normal | 0.052 | yes | T-test | <0.001 | yes | higher | |
| | Rapamycin | 1 | 0.987 | 0.644 | both normal | 0.048 | no | Welch's T-test | <0.001 | yes | higher | yes |
| | | 2 | 0.438 | 0.782 | both normal | 0.364 | yes | T-test | 0.001 | yes | higher | |
| | | 3 | 0.159 | 0.241 | both normal | 0.784 | yes | T-test | <0.001 | yes | higher | |
| Nitric oxide synthase | L-NAME | 1 | 0.033 | 0.114 | one is non-normal | 0.564 | yes | Mann-Whitney U | 0.002 | yes | higher | yes |
| | | 2 | 0.895 | 0.021 | one is non-normal | 0.633 | yes | Mann-Whitney U | 0.002 | yes | higher | |
| | | 3 | 0.696 | 0.519 | both normal | 0.162 | yes | T-test | <0.001 | yes | higher | |
| Wnt pathway | IWR-1 | 1 | 0.987 | 0.995 | both normal | 0.051 | yes | T-test | 0.001 | yes | higher | yes |
| | | 2 | 0.438 | 0.719 | both normal | 0.476 | yes | T-test | <0.001 | yes | higher | |
| | | 3 | 0.159 | 0.316 | both normal | 0.494 | yes | T-test | <0.001 | yes | higher | |
| Kinase modulator | Pyrvinium pamoate | 1 | 0.033 | 0.351 | one is non-normal | 0.190 | yes | Mann-Whitney U | 0.002 | yes | higher | no |
| | | 2 | 0.895 | 0.188 | both normal | 0.307 | yes | T-test | <0.001 | yes | higher | |
| | | 3 | 0.379 | 0.368 | both normal | 0.601 | yes | T-test | 0.234 | no | n/a | |
| | Staurosprine | 1 | 0.048 | 0.269 | one is non-normal | 0.231 | yes | Mann-Whitney U | 0.002 | yes | higher | yes |
| | | 2 | 0.056 | 0.467 | both normal | 0.281 | yes | T-test | 0.006 | yes | higher | |
| | | 3 | 0.159 | 0.004 | one is non-normal | 0.476 | yes | Mann-Whitney U | 0.002 | yes | higher | |

* represent whole body comparisons, all technical replicates, n = 6

**Table S13.** ***wsp*** **absolute counts, following exposure of** ***D. simulans flies*** **to candidate drugs in 3 total plate replicates**

| Host cellular process | Drug name | Well # | *wsp* absolute count Plate replicate 1 | | | *wsp* absolute count Plate replicate 2 | | | *wsp* absolute count Plate replicate 3 | | |
|---|---|---|---|---|---|---|---|---|---|---|---|
| | | | DMSO | Treatment | Rifampicin | DMSO | Treatment | Rifampicin | DMSO | Treatment | Rifampicin |
| Cell cycle modulation | Roscovitine | Well 1 | 1.89E+07 | 1.73E+07 | 8.67E+06 | 2.13E+07 | 1.92E+07 | 9.90E+06 | 1.20E+07 | 1.34E+07 | 7.46E+06 |
| | | | 1.53E+07 | 1.60E+07 | 9.41E+06 | 2.07E+07 | 1.73E+07 | 9.53E+06 | 1.25E+07 | 1.20E+07 | 6.45E+06 |
| | | Well 2 | 1.37E+07 | 1.59E+07 | 9.06E+06 | 1.76E+07 | 8.89E+06 | 9.01E+06 | 1.46E+07 | 9.41E+06 | 5.58E+06 |
| | | | 1.79E+07 | 1.60E+07 | 9.29E+06 | 1.82E+07 | 1.09E+07 | 8.62E+06 | 1.52E+07 | 9.50E+06 | 5.83E+06 |
| | | Well 3 | 1.85E+07 | 1.31E+07 | 8.46E+06 | 1.85E+07 | 1.43E+07 | 9.24E+06 | 1.44E+07 | 1.07E+07 | 6.18E+06 |
| | | | 1.57E+07 | 1.47E+07 | 7.84E+06 | 1.87E+07 | 1.41E+07 | 8.62E+06 | 1.36E+07 | 9.84E+06 | 6.61E+06 |
| Ubiquitin-proteasome system | Bortezomib | Well 1 | 1.89E+07 | 1.03E+07 | 8.67E+06 | 2.13E+07 | 1.44E+07 | 9.90E+06 | 1.20E+07 | 9.53E+06 | 7.46E+06 |
| | | | 1.53E+07 | 1.02E+07 | 9.41E+06 | 2.07E+07 | 1.32E+07 | 9.53E+06 | 1.25E+07 | 9.65E+06 | 6.45E+06 |
| | | Well 2 | 1.37E+07 | 9.90E+06 | 9.06E+06 | 1.76E+07 | 1.12E+07 | 9.01E+06 | 1.46E+07 | 6.87E+06 | 5.58E+06 |
| | | | 1.79E+07 | 8.78E+06 | 9.29E+06 | 1.82E+07 | 8.25E+06 | 8.62E+06 | 1.52E+07 | 9.71E+06 | 5.83E+06 |
| | | Well 3 | 1.85E+07 | 8.17E+06 | 8.46E+06 | 1.85E+07 | 9.65E+06 | 9.24E+06 | 1.44E+07 | 6.91E+06 | 6.18E+06 |
| | | | 1.57E+07 | 8.29E+06 | 7.84E+06 | 1.87E+07 | 9.47E+06 | 8.62E+06 | 1.36E+07 | 1.08E+07 | 6.61E+06 |
| Imd pathway | CAY10512 | Well 1 | 1.89E+07 | 2.16E+07 | 8.67E+06 | 2.13E+07 | 2.33E+07 | 9.90E+06 | 1.20E+07 | 1.61E+07 | 7.46E+06 |
| | | | 1.53E+07 | 1.92E+07 | 9.41E+06 | 2.07E+07 | 2.85E+07 | 9.53E+06 | 1.25E+07 | 1.77E+07 | 6.45E+06 |
| | | Well 2 | 1.37E+07 | 2.24E+07 | 9.06E+06 | 1.76E+07 | 2.64E+07 | 9.01E+06 | 1.46E+07 | 1.94E+07 | 5.58E+06 |
| | | | 1.79E+07 | 2.19E+07 | 9.29E+06 | 1.82E+07 | 2.91E+07 | 8.62E+06 | 1.52E+07 | 2.00E+07 | 5.83E+06 |
| | | Well 3 | 1.85E+07 | 2.11E+07 | 8.46E+06 | 1.85E+07 | 3.09E+07 | 9.24E+06 | 1.44E+07 | 1.94E+07 | 6.18E+06 |
| | | | 1.57E+07 | 2.06E+07 | 7.84E+06 | 1.87E+07 | 2.68E+07 | 8.62E+06 | 1.36E+07 | 2.00E+07 | 6.61E+06 |
| | SC76741 | Well 1 | 1.89E+07 | 1.34E+07 | 8.67E+06 | 2.13E+07 | 2.03E+07 | 9.90E+06 | 1.20E+07 | 1.58E+07 | 7.46E+06 |
| | | | 1.53E+07 | 1.50E+07 | 9.41E+06 | 2.07E+07 | 2.03E+07 | 9.53E+06 | 1.25E+07 | 1.32E+07 | 6.45E+06 |
| | | Well 2 | 1.37E+07 | 1.22E+07 | 9.06E+06 | 1.76E+07 | 1.83E+07 | 9.01E+06 | 1.46E+07 | 9.90E+06 | 5.58E+06 |
| | | | 1.79E+07 | 1.80E+07 | 9.29E+06 | 1.82E+07 | 1.71E+07 | 8.62E+06 | 1.52E+07 | 1.03E+07 | 5.83E+06 |
| | | Well 3 | 1.85E+07 | 1.52E+07 | 8.46E+06 | 1.85E+07 | 1.87E+07 | 9.24E+06 | 1.44E+07 | 1.41E+07 | 6.18E+06 |
| | | | 1.57E+07 | 1.58E+07 | 7.84E+06 | 1.87E+07 | 1.73E+07 | 8.62E+06 | 1.36E+07 | 1.83E+07 | 6.61E+06 |
| Phospholipase-related | M-3M3FBS | Well 1 | 1.89E+07 | 1.41E+07 | 8.67E+06 | 2.13E+07 | 2.09E+07 | 9.90E+06 | 1.20E+07 | 1.81E+07 | 7.46E+06 |
| | | | 1.53E+07 | 1.61E+07 | 9.41E+06 | 2.07E+07 | 2.07E+07 | 9.53E+06 | 1.25E+07 | 1.44E+07 | 6.45E+06 |
| | | Well 2 | 1.37E+07 | 1.40E+07 | 9.06E+06 | 1.76E+07 | 1.61E+07 | 9.01E+06 | 1.46E+07 | 2.12E+07 | 5.58E+06 |
| | | | 1.79E+07 | 1.42E+07 | 9.29E+06 | 1.82E+07 | 1.60E+07 | 8.62E+06 | 1.52E+07 | 1.61E+07 | 5.83E+06 |
| | | Well 3 | 1.85E+07 | 1.41E+07 | 8.46E+06 | 1.85E+07 | 2.00E+07 | 9.24E+06 | 1.44E+07 | 1.59E+07 | 6.18E+06 |
| | | | 1.57E+07 | 1.39E+07 | 7.84E+06 | 1.87E+07 | 2.29E+07 | 8.62E+06 | 1.36E+07 | 2.12E+07 | 6.61E+06 |
| Calcium signaling | Nicardipine HCL | Well 1 | 1.36E+07 | 1.46E+07 | 7.86E+06 | 1.86E+07 | 1.99E+07 | 1.39E+07 | 1.79E+07 | 2.45E+07 | 8.51E+06 |
| | | | 1.26E+07 | 1.52E+07 | 9.27E+06 | 1.65E+07 | 2.04E+07 | 1.37E+07 | 1.88E+07 | 2.66E+07 | 6.70E+06 |
| | | Well 2 | 1.45E+07 | 1.91E+07 | 8.15E+06 | 1.81E+07 | 2.33E+07 | 1.36E+07 | 1.87E+07 | 2.96E+07 | 6.70E+06 |
| | | | 1.32E+07 | 1.52E+07 | 8.64E+06 | 1.77E+07 | 3.55E+07 | 1.26E+07 | 1.88E+07 | 2.48E+07 | 7.46E+06 |
| | | Well 3 | 1.23E+07 | 2.00E+07 | 9.11E+06 | 1.73E+07 | 2.24E+07 | 1.09E+07 | 1.86E+07 | 2.51E+07 | 5.65E+06 |
| | | | 1.27E+07 | 1.57E+07 | 9.50E+06 | 1.59E+07 | 2.20E+07 | 1.03E+07 | 1.82E+07 | 2.32E+07 | 5.62E+06 |
| Ras/mTor pathway | Erlotinib HCl | Well 1 | 1.36E+07 | 1.47E+07 | 7.86E+06 | 1.86E+07 | 1.95E+07 | 1.39E+07 | 1.79E+07 | 2.22E+07 | 8.51E+06 |
| | | | 1.26E+07 | 1.43E+07 | 9.27E+06 | 1.65E+07 | 2.66E+07 | 1.37E+07 | 1.88E+07 | 2.29E+07 | 6.70E+06 |
| | | Well 2 | 1.45E+07 | 2.05E+07 | 8.15E+06 | 1.81E+07 | 2.64E+07 | 1.36E+07 | 1.87E+07 | 2.26E+07 | 6.70E+06 |
| | | | 1.32E+07 | 1.39E+07 | 8.64E+06 | 1.77E+07 | 2.22E+07 | 1.26E+07 | 1.88E+07 | 2.35E+07 | 7.46E+06 |
| | | Well 3 | 1.23E+07 | 1.88E+07 | 9.11E+06 | 1.73E+07 | 1.91E+07 | 1.09E+07 | 1.86E+07 | 2.55E+07 | 5.65E+06 |
| | | | 1.27E+07 | 1.52E+07 | 9.50E+06 | 1.59E+07 | 2.02E+07 | 1.03E+07 | 1.82E+07 | 2.39E+07 | 5.62E+06 |
| | Rapamycin | Well 1 | 1.31E+07 | 2.15E+07 | 6.19E+06 | 1.35E+07 | 2.47E+07 | 6.97E+06 | 1.41E+07 | 1.61E+07 | 5.83E+06 |
| | | | 1.31E+07 | 2.01E+07 | 6.07E+06 | 1.17E+07 | 2.72E+07 | 6.70E+06 | 1.17E+07 | 1.64E+07 | 5.21E+06 |
| | | Well 2 | 1.16E+07 | 1.75E+07 | 5.61E+06 | 1.31E+07 | 2.87E+07 | 7.59E+06 | 1.10E+07 | 1.75E+07 | 5.18E+06 |
| | | | 1.26E+07 | 1.97E+07 | 6.11E+06 | 1.73E+07 | 2.47E+07 | 7.15E+06 | 1.15E+07 | 1.77E+07 | 5.08E+06 |
| | | Well 3 | 1.20E+07 | 1.69E+07 | 6.97E+06 | 9.68E+06 | 2.15E+07 | 5.79E+06 | 1.08E+07 | 1.69E+07 | 3.98E+06 |
| | | | 9.68E+06 | 1.67E+07 | 7.11E+06 | 1.03E+07 | 2.29E+07 | 5.50E+06 | 1.02E+07 | 1.73E+07 | 3.98E+06 |
| Nitric oxide synthase | L-NAME | Well 1 | 1.36E+07 | 1.48E+07 | 7.86E+06 | 1.86E+07 | 2.18E+07 | 1.39E+07 | 1.79E+07 | 2.50E+07 | 8.51E+06 |
| | | | 1.26E+07 | 1.23E+07 | 9.27E+06 | 1.65E+07 | 1.94E+07 | 1.37E+07 | 1.88E+07 | 2.59E+07 | 6.70E+06 |
| | | Well 2 | 1.45E+07 | 1.44E+07 | 8.15E+06 | 1.81E+07 | 1.93E+07 | 1.36E+07 | 1.87E+07 | 2.35E+07 | 6.70E+06 |
| | | | 1.32E+07 | 1.42E+07 | 8.64E+06 | 1.77E+07 | 2.04E+07 | 1.26E+07 | 1.88E+07 | 3.19E+07 | 7.46E+06 |
| | | Well 3 | 1.23E+07 | 1.49E+07 | 9.11E+06 | 1.73E+07 | 2.29E+07 | 1.09E+07 | 1.86E+07 | 2.06E+07 | 5.65E+06 |
| | | | 1.27E+07 | 1.58E+07 | 9.50E+06 | 1.59E+07 | 2.17E+07 | 1.03E+07 | 1.82E+07 | 2.11E+07 | 5.62E+06 |
| Wnt pathway | IWR-1 | Well 1 | 1.36E+07 | 1.41E+07 | 7.86E+06 | 1.86E+07 | 2.03E+07 | 1.39E+07 | 1.79E+07 | 2.38E+07 | 8.51E+06 |
| | | | 1.26E+07 | 1.61E+07 | 9.27E+06 | 1.65E+07 | 2.36E+07 | 1.37E+07 | 1.88E+07 | 2.73E+07 | 6.70E+06 |
| | | Well 2 | 1.45E+07 | 1.40E+07 | 8.15E+06 | 1.81E+07 | 2.03E+07 | 1.36E+07 | 1.87E+07 | 2.68E+07 | 6.70E+06 |
| | | | 1.32E+07 | 1.42E+07 | 8.64E+06 | 1.77E+07 | 1.99E+07 | 1.26E+07 | 1.88E+07 | 2.35E+07 | 7.46E+06 |
| | | Well 3 | 1.23E+07 | 1.41E+07 | 9.11E+06 | 1.73E+07 | 2.08E+07 | 1.09E+07 | 1.86E+07 | 2.55E+07 | 5.65E+06 |
| | | | 1.27E+07 | 1.39E+07 | 9.50E+06 | 1.59E+07 | 2.09E+07 | 1.03E+07 | 1.82E+07 | 2.36E+07 | 5.62E+06 |
| Kinase modulator | Staurosprine | Well 1 | 1.31E+07 | 1.46E+07 | 6.19E+06 | 1.35E+07 | 1.80E+07 | 6.97E+06 | 1.41E+07 | 1.42E+07 | 5.83E+06 |
| | | | 1.31E+07 | 1.31E+07 | 6.07E+06 | 1.17E+07 | 1.92E+07 | 6.70E+06 | 1.17E+07 | 1.46E+07 | 5.21E+06 |
| | | Well 2 | 1.16E+07 | 1.17E+07 | 5.61E+06 | 1.31E+07 | 1.71E+07 | 7.59E+06 | 1.10E+07 | 1.65E+07 | 5.18E+06 |
| | | | 1.26E+07 | 1.34E+07 | 6.11E+06 | 1.73E+07 | 1.61E+07 | 7.15E+06 | 1.15E+07 | 1.17E+07 | 5.08E+06 |
| | | Well 3 | 1.20E+07 | 1.42E+07 | 6.97E+06 | 9.68E+06 | 1.64E+07 | 5.79E+06 | 1.08E+07 | 1.20E+07 | 3.98E+06 |
| | | | 9.68E+06 | 1.42E+07 | 7.11E+06 | 1.03E+07 | 1.69E+07 | 5.50E+06 | 1.02E+07 | 1.23E+07 | 3.98E+06 |

**Table S14. Statistical analyses of wsp absolute counts from candidate drugs tested in 3 total plate replicates in D. simulans**

| Host cellular process | Drug name | Plate replicate # | Shapiro-wilk: p-values, interpretation | | | Levene's test: p-values, interpretation | | Outcome: analysis of different combinations | | | | Interpreted as a hit? |
|---|---|---|---|---|---|---|---|---|---|---|---|---|
| | | | DMSO | Treatment | Normal distribution? | Overall value | Variance homogeneous? | Analysis required | Resulting p-value | Significantly different? | Abundance relative to control | |
| Cell cycle modulation | Roscovitine | 1 | 0.479 | 0.488 | both normal | 0.166 | yes | T-test | 0.283 | no | n/a | no |
| | | 2 | 0.253 | 0.876 | both normal | 0.133 | yes | T-test | 0.013 | yes | lower | |
| | | 3 | 0.629 | 0.253 | both normal | 0.690 | yes | T-test | 0.006 | yes | lower | |
| Ubiquitin-proteasome system | Bortezomib | 1 | 0.479 | 0.154 | both normal | 0.031 | no | Welch's T-test | <0.001 | yes | lower | yes |
| | | 2 | 0.253 | 0.639 | both normal | 0.258 | yes | T-test | <0.001 | yes | lower | |
| | | 3 | 0.629 | 0.108 | both normal | 0.838 | yes | T-test | <0.001 | yes | lower | |
| Imd pathway | CAY10512 | 1 | 0.479 | 0.705 | both normal | 0.053 | yes | T-test | 0.001 | yes | higher | yes |
| | | 2 | 0.253 | 0.920 | both normal | 0.241 | yes | T-test | <0.001 | yes | higher | |
| | | 3 | 0.629 | 0.092 | both normal | 0.978 | yes | T-test | <0.001 | yes | higher | |
| | SC76741 | 1 | 0.479 | 0.927 | both normal | 0.567 | yes | T-test | 0.171 | no | n/a | no |
| | | 2 | 0.253 | 0.267 | both normal | 0.952 | yes | T-test | 0.561 | no | n/a | |
| | | 3 | 0.629 | 0.743 | both normal | 0.104 | yes | T-test | 0.936 | no | n/a | |
| Phospholipase-related | M-3M3FBS | 1 | 0.479 | 0.001 | one is non-normal | 0.012 | no | T-test w/bootstrap | 0.044 | yes | higher | no |
| | | 2 | 0.253 | 0.253 | both normal | 0.300 | yes | T-test | 0.840 | no | n/a | |
| | | 3 | 0.629 | 0.283 | both normal | 0.087 | yes | T-test | 0.009 | yes | higher | |
| Calcium signaling | Nicardipine HCl | 1 | 0.576 | 0.060 | both normal | 0.264 | yes | T-test | 0.006 | yes | higher | yes |
| | | 2 | 0.902 | 0.006 | one is non-normal | 0.304 | yes | Mann-Whitney U | 0.002 | yes | higher | |
| | | 3 | 0.131 | 0.377 | both normal | 0.149 | yes | T-test | <0.001 | yes | higher | |
| Ras/mTor pathway | Erlotinib HCl | 1 | 0.576 | 0.096 | both normal | 0.212 | yes | T-test | 0.024 | yes | higher | yes |
| | | 2 | 0.902 | 0.113 | both normal | 0.066 | yes | T-test | 0.006 | yes | higher | |
| | | 3 | 0.131 | 0.522 | both normal | 0.113 | yes | T-test | <0.001 | yes | higher | |
| | Rapamycin | 1 | 0.184 | 0.355 | both normal | 0.113 | yes | T-test | <0.001 | yes | higher | yes |
| | | 2 | 0.566 | 0.846 | both normal | 0.902 | yes | T-test | <0.001 | yes | higher | |
| | | 3 | 0.168 | 0.647 | both normal | 0.427 | yes | T-test | <0.001 | yes | higher | |
| Nitric oxide synthase | L-NAME | 1 | 0.576 | 0.395 | both normal | 0.716 | yes | T-test | 0.056 | no | n/a | no |
| | | 2 | 0.902 | 0.480 | both normal | 0.218 | yes | T-test | 0.001 | yes | higher | |
| | | 3 | 0.131 | 0.400 | both normal | 0.054 | yes | T-test | 0.004 | yes | higher | |
| Wnt pathway | IWR-1 | 1 | 0.576 | 0.001 | one is non-normal | 0.583 | yes | Mann-Whitney U | 0.041 | yes | higher | yes |
| | | 2 | 0.902 | 0.019 | one is non-normal | 0.975 | yes | Mann-Whitney U | 0.002 | yes | higher | |
| | | 3 | 0.131 | 0.150 | both normal | 0.011 | no | Welch's T-test | <0.001 | yes | higher | |
| Kinase modulator | Staurosprine | 1 | 0.184 | 0.376 | both normal | 0.793 | yes | T-test | 0.050 | yes | higher | no |
| | | 2 | 0.566 | 0.536 | both normal | 0.139 | yes | T-test | 0.003 | yes | higher | |
| | | 3 | 0.168 | 0.380 | both normal | 0.247 | yes | T-test | 0.061 | no | n/a | |

* represent whole body comparisons, all technical replicates, n = 6

**Table S15.Genetic tools used for disrupting candidate host processes**

| Host cellular process | Gene name | Symbol | BDSC stock # | Insertion # | Chromosome |
|---|---|---|---|---|---|
| Ubiquitin-proteasome | *Proteasome alpha6 subunit* | *Pros alpha6* | 53974 | P{TRiP.HMJ21359}attP40 | 2 |
| | *Ubiquitin conjugating enzyme 6* | *Ubc6* | 42631 | P{TRiP.HMS02466}attP40 | 2 |
| IMD | *NF kappa B/ Relish* | *Rel* | 33661 | P{TRiP.HMS00070}attP2 | 3 |
| | *TGF-β activated kinase 1* | *Tak1* | 53377 | P{TRiP.HMJ02053}attP40 | 2 |
| Ca channel | *Calcium channel protein alpha1 subunit D* | *Ca-alpha1D* | 33413 | P{TRiP.HMS00294}attP2 | 3 |
| | *Calcium channel alpha1 subunit, type A/ Cacophany* | *Cac* | 77174 | P{TRiP.HMC03320}attP40 | 2 |
| Wnt | *Beta-catenin/Armadillo* | *Arm* | 35004 | P{TRiP.HMS01414}attP2 | 3 |
| | *Glycogen Synthase Kinase 3/ Shaggy* | *Sgg* | 38293 | P{TRiP.HMS01751}attP40 | 2 |
| Ras/mTOR | *Epidermal growth factor receptor* | *EGFR* | 60012 | P{TRiP.HMS05003}attP40 | 2 |
| | *Target of rapamycin* | *Tor* | 58312 | P{TRiP.HMJ22419}attP40 | 2 |

**Table S16. Reported expression levels for endogenous A*ctin5C* and *daughterless* genes**

| Developmental stages | *Actin5C* | *daughterless* |
|---|---|---|
| Embryo 0-2hr | 1417 | 103 |
| Embryo12-14hr | 1285 | 48 |
| Embryo 22-24hr | 1825 | 20 |
| Larva L1 | 1325 | 22 |
| Larva L2 | 1744 | 13 |
| Larva L3 12hr | 1063 | 12 |
| Pupa 1d | 1978 | 41 |
| Pupa 3d | 382 | 27 |
| Pupa 4d | 1458 | 24 |
| Adult female 1d | 1133 | 43 |
| Adult female 5d | 1126 | 54 |

* The data was obtained from the modENCODE project (Graveley et al., 2011), as communicated via the community resource website flybase.net.

**Table S17. Body-wide *wsp* abundance, following the crosses of RNAi lines with daughterless GAL4 driver**

| Host cellular process | RNAi target | Tube # | *wsp* absolute count, experimental replicate 1 | | | *wsp* absolute count, experimental replicate 2 | | |
|---|---|---|---|---|---|---|---|---|
| | | | OreR outcrossed control | Non-expressing | RNAi Expressing | OreR outcrossed control | Non-expressing | RNAi Expressing |
| Ubiquitin-proteasome | *Pros alpha6* | Tube 1 | 5.57E+06 | 4.48E+06 | 5.64E+06 | 6.66E+06 | 6.67E+06 | 7.34E+06 |
| | | | 5.31E+06 | 4.69E+06 | 5.31E+06 | 6.68E+06 | 7.40E+06 | 7.47E+06 |
| | | Tube 2 | 5.21E+06 | 4.87E+06 | 5.98E+06 | 6.46E+06 | 6.62E+06 | 7.34E+06 |
| | | | 5.26E+06 | 4.57E+06 | 5.41E+06 | 6.32E+06 | 6.23E+06 | 8.00E+06 |
| | | Tube 3 | 4.53E+06 | 3.76E+06 | 5.64E+06 | 7.23E+06 | 7.01E+06 | 9.90E+06 |
| | | | 4.79E+06 | 5.17E+06 | 4.93E+06 | 7.29E+06 | 6.85E+06 | 6.64E+06 |
| | *Ubc6* | Tube 1 | 1.11E+07 | 1.03E+07 | 1.20E+07 | 5.41E+06 | 6.46E+06 | 7.04E+06 |
| | | | 1.10E+07 | 1.36E+07 | 1.31E+07 | 5.28E+06 | 6.79E+06 | 7.57E+06 |
| | | Tube 2 | 1.28E+07 | 1.36E+07 | 1.26E+07 | 5.67E+06 | 6.53E+06 | 7.97E+06 |
| | | | 1.30E+07 | 1.11E+07 | 1.25E+07 | 5.13E+06 | 6.12E+06 | 8.28E+06 |
| | | Tube 3 | 1.22E+07 | 1.10E+07 | 1.29E+07 | 5.62E+06 | 7.16E+06 | 6.85E+06 |
| | | | 1.17E+07 | 1.03E+07 | 1.35E+07 | 5.72E+06 | 7.80E+06 | 7.12E+06 |
| IMD | *Relish* | Tube 1 | 4.91E+06 | n/a | 5.79E+06 | 4.63E+06 | n/a | 5.15E+06 |
| | | | 4.76E+06 | | 4.85E+06 | 5.64E+06 | | 4.45E+06 |
| | | Tube 2 | 4.60E+06 | n/a | 4.48E+06 | 6.74E+06 | n/a | 5.68E+06 |
| | | | 5.05E+06 | | 4.14E+06 | 6.27E+06 | | 5.05E+06 |
| | | Tube 3 | 4.34E+06 | n/a | 3.85E+06 | 5.57E+06 | n/a | 5.05E+06 |
| | | | 4.17E+06 | | 3.78E+06 | 6.92E+06 | | 4.88E+06 |
| | *Tak1* | Tube 1 | 5.08E+06 | n/a | 5.72E+06 | 4.91E+06 | n/a | 6.23E+06 |
| | | | 4.54E+06 | | 5.46E+06 | 6.79E+06 | | 6.65E+06 |
| | | Tube 2 | 5.15E+06 | n/a | 4.85E+06 | 4.36E+06 | n/a | 4.85E+06 |
| | | | 5.53E+06 | | 4.31E+06 | 3.98E+06 | | 4.34E+06 |
| | | Tube 3 | 5.68E+06 | n/a | 4.22E+06 | 5.18E+06 | n/a | 3.98E+06 |
| | | | 5.91E+06 | | 4.03E+06 | 4.14E+06 | | 5.57E+06 |
| Ca channel | *Ca1alpha D* | Tube 1 | 3.01E+06 | n/a | 4.70E+06 | 8.43E+06 | n/a | 8.54E+06 |
| | | | 3.15E+06 | | 3.80E+06 | 8.00E+06 | | 9.49E+06 |
| | | Tube 2 | 4.24E+06 | n/a | 4.33E+06 | 1.49E+07 | n/a | 9.01E+06 |
| | | | 3.54E+06 | | 3.85E+06 | 8.77E+06 | | 7.84E+06 |
| | | Tube 3 | 4.00E+06 | n/a | 3.34E+06 | 7.54E+06 | n/a | 8.71E+06 |
| | | | 3.82E+06 | | 3.69E+06 | 7.59E+06 | | 8.60E+06 |
| | *Cac* | Tube 1 | 3.51E+06 | n/a | 4.84E+06 | 8.54E+06 | n/a | 8.16E+06 |
| | | | 3.57E+06 | | 4.30E+06 | 8.66E+06 | | 8.05E+06 |
| | | Tube 2 | 4.70E+06 | n/a | 4.14E+06 | 9.31E+06 | n/a | 9.75E+06 |
| | | | 4.50E+06 | | 3.80E+06 | 9.37E+06 | | 9.07E+06 |
| | | Tube 3 | 4.45E+06 | n/a | 3.59E+06 | 8.16E+06 | n/a | 1.23E+07 |
| | | | 4.56E+06 | | 3.41E+06 | 7.59E+06 | | 8.60E+06 |
| Wnt | *Arm* | Tube 1 | 7.54E+06 | n/a | 7.79E+06 | 2.48E+06 | n/a | 3.08E+06 |
| | | | 6.61E+06 | | 7.54E+06 | 2.92E+06 | | 3.25E+06 |
| | | Tube 2 | 6.83E+06 | n/a | 7.11E+06 | 2.27E+06 | n/a | 3.08E+06 |
| | | | 6.57E+06 | | 7.39E+06 | 2.59E+06 | | 3.40E+06 |
| | | Tube 3 | 6.97E+06 | n/a | 6.88E+06 | 2.58E+06 | n/a | 3.49E+06 |
| | | | 6.92E+06 | | 7.74E+06 | 2.51E+06 | | 3.22E+06 |
| | *Sgg* | Tube 1 | 5.57E+06 | 5.28E+06 | 5.05E+06 | 4.98E+06 | 3.42E+06 | 4.05E+06 |
| | | | 4.67E+06 | 5.11E+06 | 5.53E+06 | 4.40E+06 | 3.72E+06 | 4.01E+06 |
| | | Tube 2 | 4.50E+06 | 4.38E+06 | 5.57E+06 | 3.37E+06 | 3.42E+06 | 4.46E+06 |
| | | | 3.47E+06 | 6.02E+06 | 5.22E+06 | 3.36E+06 | 3.54E+06 | 4.47E+06 |
| | | Tube 3 | 4.33E+06 | 5.90E+06 | 5.54E+06 | 3.91E+06 | 3.86E+06 | 3.65E+06 |
| | | | 4.02E+06 | 6.66E+06 | 5.69E+06 | 3.99E+06 | 3.55E+06 | 4.04E+06 |
| Ras/mTor | *EGFR* | Tube 1 | 8.33E+06 | 6.55E+06 | 9.68E+06 | 5.23E+06 | 4.15E+06 | 6.14E+06 |
| | | | 7.88E+06 | 7.41E+06 | 1.01E+07 | 5.40E+06 | 3.90E+06 | 6.07E+06 |
| | | Tube 2 | 8.22E+06 | 6.36E+06 | 1.07E+07 | 6.45E+06 | 4.78E+06 | 4.54E+06 |
| | | | 7.87E+06 | 6.72E+06 | 9.40E+06 | 5.94E+06 | 4.09E+06 | 5.15E+06 |
| | | Tube 3 | 8.17E+06 | 6.85E+06 | 8.80E+06 | 6.54E+06 | 4.43E+06 | 5.27E+06 |
| | | | 8.35E+06 | 6.25E+06 | 9.12E+06 | 6.83E+06 | 4.23E+06 | 6.38E+06 |
| | *Tor* | Tube 1 | 3.49E+06 | 3.61E+06 | 4.48E+06 | 3.63E+06 | 4.29E+06 | 4.82E+06 |
| | | | 3.38E+06 | 3.61E+06 | 5.04E+06 | 3.83E+06 | 4.64E+06 | 4.55E+06 |
| | | Tube 2 | 3.88E+06 | 3.56E+06 | 4.34E+06 | 3.01E+06 | 3.74E+06 | 4.17E+06 |
| | | | 3.49E+06 | 3.75E+06 | 4.60E+06 | 2.97E+06 | 3.83E+06 | 4.92E+06 |
| | | Tube 3 | 3.95E+06 | 3.61E+06 | 4.09E+06 | 3.62E+06 | 4.43E+06 | 4.54E+06 |
| | | | 3.90E+06 | 3.31E+06 | 4.63E+06 | 3.54E+06 | 4.13E+06 | 5.18E+06 |

**Table S18. Statistical analyses of *wsp* absolute counts from RNAi crosses with daughterless GAL4 driver**

| Host cellular process | RNAi target | Replicate # | Control used for comparison | Shapiro-wilk: p-values, interpretation | | | Levene's test: p-values, interpretation | | Outcome: analysis of different combinations | | | | Interpreted as a hit? |
|---|---|---|---|---|---|---|---|---|---|---|---|---|---|
| | | | | Control | RNAi expressing | Normal distribution? | Overall value | Variance homogeneous? | Analysis required | Resulting p-value | Significantly different? | Abundance relative to control | |
| Ubiquitin-proteasome system | *Pros alpha6* | 1 | OreR outcross | 0.565 | 0.921 | both normal | 1.000 | yes | T-test | 0.11 | no | n/a | no |
| | | 2 | OreR outcross | 0.291 | 0.110 | both normal | 0.378 | yes | T-test | 0.065 | no | n/a | |
| | | 1 | Non-expressed | 0.626 | 0.921 | both normal | 0.750 | yes | T-test | 0.004 | yes | higher | no |
| | | 2 | Non-expressed | 0.979 | 0.110 | both normal | 0.372 | yes | T-test | 0.071 | no | n/a | |
| | *Ubc6* | 1 | OreR outcross | 0.464 | 0.996 | both normal | 0.138 | yes | T-test | 0.077 | no | n/a | no |
| | | 2 | OreR outcross | 0.494 | 0.550 | both normal | 0.065 | yes | T-test | <0.001 | yes | higher | |
| | | 1 | Non-expressed | 0.039 | 0.996 | one is non-normal | 0.193 | yes | Mann-Whitney U | 0.394 | no | n/a | no |
| | | 2 | Non-expressed | 0.731 | 0.550 | both normal | 0.888 | yes | T-test | 0.077 | no | n/a | |
| Imd pathway | *Relish* | 1 | OreR outcross | 0.851 | 0.374 | both normal | 0.212 | yes | T-test | 0.653 | no | n/a | no |
| | | 2 | OreR outcross | 0.694 | 0.693 | both normal | 0.067 | yes | T-test | 0.038 | yes | lower | |
| | | 1 | Non-expressed | n/a | n/a | n/a | n/a | n/a | n/a | n/a | n/a | n/a | n/a |
| | | 2 | Non-expressed | n/a | n/a | n/a | n/a | n/a | n/a | n/a | n/a | n/a | |
| | *Tak1* | 1 | OreR outcross | 0.849 | 0.363 | both normal | 0.321 | yes | T-test | 0.147 | no | n/a | no |
| | | 2 | OreR outcross | 0.187 | 0.740 | both normal | 0.680 | yes | T-test | 0.548 | no | n/a | |
| | | 1 | Non-expressed | n/a | n/a | n/a | n/a | n/a | n/a | n/a | n/a | n/a | n/a |
| | | 2 | Non-expressed | n/a | n/a | n/a | n/a | n/a | n/a | n/a | n/a | n/a | |
| Calcium signaling | *Ca1alpha* | 1 | OreR outcross | 0.752 | 0.742 | both normal | 0.764 | yes | T-test | 0.272 | no | n/a | no |
| | | 2 | OreR outcross | 0.002 | 0.872 | one is non-normal | 0.333 | yes | Mann-Whitney U | 0.394 | no | n/a | |
| | | 1 | Non-expressed | n/a | n/a | n/a | n/a | n/a | n/a | n/a | n/a | n/a | n/a |
| | | 2 | Non-expressed | n/a | n/a | n/a | n/a | n/a | n/a | n/a | n/a | n/a | |
| | *Cac* | 1 | OreR outcross | 0.04 | 0.843 | one is non-normal | 0.848 | yes | Mann-Whitney U | 0.589 | no | n/a | no |
| | | 2 | OreR outcross | 0.678 | 0.093 | both normal | 0.337 | yes | T-test | 0.334 | no | n/a | |
| | | 1 | Non-expressed | n/a | n/a | n/a | n/a | n/a | n/a | n/a | n/a | n/a | n/a |
| | | 2 | Non-expressed | n/a | n/a | n/a | n/a | n/a | n/a | n/a | n/a | n/a | |
| Wnt | *Armadillo* | 1 | OreR | 0.253 | 0.633 | both normal | 0.726 | yes | T-test | 0.034 | yes | higher | yes |
| | | 2 | OreR | 0.539 | 0.463 | both normal | 0.875 | yes | T-test | <0.001 | yes | higher | |
| | | 1 | Non-expressed | n/a | n/a | n/a | n/a | n/a | n/a | n/a | n/a | n/a | n/a |
| | | 2 | Non-expressed | n/a | n/a | n/a | n/a | n/a | n/a | n/a | n/a | n/a | |
| | *Sgg* | 1 | OreR | 0.912 | 0.250 | both normal | 0.160 | yes | T-test | 0.008 | yes | higher | no |
| | | 2 | OreR | 0.541 | 0.310 | both normal | 0.211 | yes | T-test | 0.702 | no | no | |
| | | 1 | Non-expressed | 0.964 | 0.250 | both normal | 0.035 | no | Welch's T-test | 0.727 | no | no | no |
| | | 2 | Non-expressed | 0.338 | 0.310 | both normal | 0.407 | yes | T-test | 0.005 | yes | higher | |
| Ras/mTOR | *EGFR* | 1 | OreR | 0.149 | 0.934 | both normal | 0.073 | yes | T-test | <0.001 | yes | higher | no |
| | | 2 | OreR | 0.497 | 0.490 | both normal | 0.715 | yes | T-test | 0.258 | no | no | |
| | | 1 | Non-expressed | 0.587 | 0.934 | both normal | 0.273 | yes | T-test | <0.001 | yes | higher | yes |
| | | 2 | Non-expressed | 0.731 | 0.490 | both normal | 0.024 | no | Welch's T-test | 0.002 | yes | higher | |
| | *Tor* | 1 | OreR | 0.094 | 0.917 | both normal | 0.985 | yes | T-test | <0.001 | yes | higher | yes |
| | | 2 | OreR | 0.149 | 0.939 | both normal | 0.902 | yes | T-test | <0.001 | yes | higher | |
| | | 1 | Non-expressed | 0.135 | 0.917 | both normal | 0.165 | yes | T-test | <0.001 | yes | higher | yes |
| | | 2 | Non-expressed | 0.806 | 0.939 | both normal | 1.000 | yes | T-test | 0.028 | yes | higher | |

* represent whole body comparisons, all technical replicates, n = 6

**Table S19. Body-wide *wsp* abundance, following the crosses of *tor* and *armadillo* RNAi lines with daughterless GAL4 driver.**

| Bodywide *wsp* absolute counts from RNAi expressing vs. control conditions, all technical replicates | | | | | | |
|---|---|---|---|---|---|---|
| Host cellular Process | RNAi target | Tube # | Experimental replicate 1 | | Experimental replicate 2 | |
| | | | OreR outcrossed control | RNAi Expressing | OreR outcrossed control | RNAi Expressing |
| Wnt | *Armadillo* | Tube 1 | 4.01E+06 | 5.72E+06 | 4.72E+06 | 5.20E+06 |
| | | | 4.09E+06 | 5.76E+06 | 5.20E+06 | 5.32E+06 |
| | | | 3.16E+06 | 6.39E+06 | 4.90E+06 | 4.90E+06 |
| | | Tube 2 | 4.01E+06 | 5.99E+06 | 5.04E+06 | 5.33E+06 |
| | | | 3.68E+06 | 5.79E+06 | 4.53E+06 | 5.85E+06 |
| | | | 3.88E+06 | 6.19E+06 | 4.54E+06 | 6.14E+06 |
| | | Tube 3 | 3.61E+06 | 7.06E+06 | 4.13E+06 | 5.44E+06 |
| | | | 3.42E+06 | 6.35E+06 | 5.15E+06 | 5.56E+06 |
| | | | 3.75E+06 | 5.08E+06 | 5.13E+06 | 4.98E+06 |
| | | Tube 4 | 3.80E+06 | 5.99E+06 | 4.23E+06 | 4.47E+06 |
| | | | 3.73E+06 | 4.63E+06 | 3.92E+06 | 5.60E+06 |
| | | | 3.80E+06 | 5.99E+06 | 4.21E+06 | 6.05E+06 |
| | | Tube 5 | 4.51E+06 | 4.48E+06 | 4.95E+06 | 5.43E+06 |
| | | | 3.98E+06 | 5.64E+06 | 4.66E+06 | 5.85E+06 |
| | | | 4.95E+06 | 6.35E+06 | 4.42E+06 | 7.24E+06 |
| | | Tube 6 | 4.11E+06 | 5.68E+06 | 5.05E+06 | 5.16E+06 |
| | | | 4.82E+06 | 5.79E+06 | 4.69E+06 | 5.33E+06 |
| | | | 4.28E+06 | 6.27E+06 | 4.06E+06 | 6.48E+06 |
| Egfr/mTor | *Tor* | Tube 1 | 3.90E+06 | 5.55E+06 | 4.95E+06 | 5.51E+06 |
| | | | 3.71E+06 | 5.05E+06 | 4.66E+06 | 5.14E+06 |
| | | | 3.86E+06 | 5.08E+06 | 4.42E+06 | 5.94E+06 |
| | | Tube 2 | 3.80E+06 | 5.21E+06 | 4.45E+06 | 7.54E+06 |
| | | | 3.56E+06 | 5.11E+06 | 4.26E+06 | 6.47E+06 |
| | | | 3.48E+06 | 5.27E+06 | 4.36E+06 | 6.72E+06 |
| | | Tube 3 | 3.94E+06 | 5.54E+06 | 3.86E+06 | 5.93E+06 |
| | | | 3.78E+06 | 4.62E+06 | 3.94E+06 | 5.74E+06 |
| | | | 3.70E+06 | 4.12E+06 | 3.71E+06 | 5.44E+06 |
| | | Tube 4 | 3.75E+06 | 5.05E+06 | 3.81E+06 | 6.69E+06 |
| | | | 3.61E+06 | 4.90E+06 | 3.69E+06 | 5.62E+06 |
| | | | 3.33E+06 | 5.06E+06 | 3.66E+06 | 5.67E+06 |
| | | Tube 5 | 3.87E+06 | 5.16E+06 | 4.10E+06 | 6.01E+06 |
| | | | 4.03E+06 | 5.22E+06 | 4.33E+06 | 5.85E+06 |
| | | | 3.88E+06 | 4.97E+06 | 4.06E+06 | 5.81E+06 |
| | | Tube 6 | 3.58E+06 | 5.43E+06 | 4.59E+06 | 5.75E+06 |
| | | | 3.43E+06 | 5.22E+06 | 4.53E+06 | 6.06E+06 |
| | | | 3.01E+06 | 5.29E+06 | 4.14E+06 | 5.02E+06 |

**Table S20. Statistical analyses of *wsp* absolute counts following the crosses of *tor* and *armadillo* RNAi lines with daughterless GAL4 driver.**

| Host cellular process | RNAi target | Replicate # | Control used for comparison | Shapiro-wilk: p-values, interpretation | | | Levene's test: p-values, interpretation | | Outcome: analysis of different combinations | | | | "n" required to satisfy $\beta < 0.05$ (out of 18) | Consistent high power? |
|---|---|---|---|---|---|---|---|---|---|---|---|---|---|---|
| | | | | Control | RNAi expressing | Normal distribution? | Overall value | Variance homogeneous? | Analysis required | Resulting p-value | Significantly different? | Abundance relative to control | | |
| Wnt | *Armadillo* | 1 | Ore-R outcross | 0.447 | 0.158 | both normal | 0.358 | yes | T-test | < 0.001 | yes | higher | 6 | yes |
| | | 2 | Ore-R outcross | 0.291 | 0.344 | both normal | 0.370 | yes | T-test | < 0.001 | yes | higher | 12 | |
| Egfr/mTor | *Tor* | 1 | Ore-R outcross | 0.174 | 0.017 | one is non-normal | 0.679 | yes | Mann-Whitney U | < 0.001 | yes | higher | 4 | yes |
| | | 2 | Ore-R outcross | 0.683 | 0.136 | both normal | 0.354 | yes | T-test | < 0.001 | yes | higher | 4 | |

* represent whole body comparisons, all technical replicates, n = 18

**Table S21. Whole body *wsp* abundance, in response to mifepristone treatment**

| *wsp* absolute counts, whole-body samples DMSO vs mifepristone treatment | | |
|---|---|---|
| Condition | Plate 1 | Plate 2 |
| DMSO | 6.55E+06 | 8.09E+06 |
| Whole body | 8.62E+06 | 7.50E+06 |
| Well 1 | 6.38E+06 | 1.11E+07 |
| DMSO | 8.53E+06 | 1.08E+07 |
| Whole body | 7.20E+06 | 1.06E+07 |
| Well 2 | 6.91E+06 | 8.73E+06 |
| DMSO | 6.78E+06 | 1.15E+07 |
| Whole body | 5.97E+06 | 7.30E+06 |
| Well 3 | 6.32E+06 | 8.22E+06 |
| DMSO | 7.46E+06 | 8.44E+06 |
| Whole body | 9.93E+06 | 7.33E+06 |
| Well 4 | 7.83E+06 | 1.04E+07 |
| DMSO | 6.41E+06 | 1.06E+07 |
| Whole body | 5.74E+06 | 1.02E+07 |
| Well 5 | 6.68E+06 | 1.00E+07 |
| DMSO | 6.17E+06 | 9.38E+06 |
| Whole body | 6.59E+06 | 1.02E+07 |
| Well 6 | 4.66E+06 | 8.18E+06 |
| Mifepristone | 9.94E+06 | 1.25E+07 |
| Whole body | 1.05E+07 | 1.05E+07 |
| Well 1 | 9.66E+06 | 1.25E+07 |
| Mifepristone | 7.67E+06 | 1.33E+07 |
| Whole body | 6.65E+06 | 1.25E+07 |
| Well 2 | 7.60E+06 | 1.30E+07 |
| Mifepristone | 9.10E+06 | 1.12E+07 |
| Whole body | 9.09E+06 | 8.64E+06 |
| Well 3 | 7.24E+06 | 1.03E+07 |
| Mifepristone | 7.58E+06 | 1.08E+07 |
| Whole body | 6.64E+06 | 1.01E+07 |
| Well 4 | 7.45E+06 | 9.78E+06 |
| Mifepristone | 7.65E+06 | 1.61E+07 |
| Whole body | 6.88E+06 | 1.33E+07 |
| Well 5 | 6.48E+06 | 1.05E+07 |
| Mifepristone | 8.87E+06 | 1.12E+07 |
| Whole body | 8.13E+06 | 1.05E+07 |
| Well 6 | 7.26E+06 | 1.07E+07 |

**Table S22. Statistical analyses of *wsp* abundance from whole body samples, in response to mifepristone treatment**

| Plate replicate # | Shapiro-wilk: p-values, interpretation | | | Levene's test: p-values, interpretation | | Outcome: analysis of different combinations | | | "n" required to satisfy $\beta < 0.05$ |
|---|---|---|---|---|---|---|---|---|---|
| | DMSO | Mifepristone | Normal distribution? | Overall value | Variance homogeneous? | Analysis required | Resulting p-value | Significantly different? | |
| 1 | 0.261 | 0.115 | yes | 0.751 | yes | T-test | 0.011 | yes | > 18 |
| 2 | 0.117 | 0.145 | yes | 0.794 | yes | T-test | <0.001 | yes | > 18 |

* represent whole body comparisons, all technical replicates, n = 18

**Table S23. Body-wide *wsp* abundance, following the crosses of RNAi lines with Gene-Switch GAL4 driver**

| Host cellular process | RNAi target | Tube # | *wsp* absolute count, Experimental replicate 1 | | | | *wsp* absolute count, Experimental replicate 2 | | | |
|---|---|---|---|---|---|---|---|---|---|---|
| | | | DMSO control | | DMSO+Mifepristone | | DMSO control | | DMSO+Mifepristone | |
| | | | Non-expressing | RNAi Expressing | Non-expressing | RNAi Expressing | Non-expressing | RNAi Expressing | Non-expressing | RNAi Expressing |
| Wnt | *Arm* | Tube 1 | 2.72E+06 | 2.34E+06 | 2.18E+06 | 1.16E+06 | 1.86E+06 | 1.91E+06 | 2.16E+06 | 1.16E+06 |
| | | | 2.04E+06 | 2.06E+06 | 2.51E+06 | 1.08E+06 | 2.96E+06 | 2.16E+06 | 2.08E+06 | 1.03E+06 |
| | | | 1.91E+06 | 2.18E+07 | 3.58E+06 | 1.15E+06 | 2.52E+06 | 2.27E+06 | 3.58E+06 | 1.06E+06 |
| | | Tube 2 | 1.90E+06 | 1.91E+06 | 2.85E+06 | 1.56E+06 | 1.91E+06 | 1.83E+06 | 2.86E+06 | 1.74E+06 |
| | | | 2.66E+06 | 1.36E+06 | 2.85E+06 | 1.41E+06 | 2.38E+06 | 2.40E+06 | 2.52E+06 | 1.38E+06 |
| | | | 1.89E+06 | 2.76E+06 | 2.61E+06 | 1.52E+06 | 1.91E+06 | 2.14E+06 | 2.18E+06 | 1.53E+06 |
| | | Tube 3 | 2.06E+06 | 2.04E+06 | 2.54E+06 | 1.04E+06 | 2.76E+06 | 2.51E+06 | 2.16E+06 | 1.76E+06 |
| | | | 1.91E+06 | 2.51E+06 | 2.45E+06 | 1.09E+06 | 2.46E+06 | 2.06E+06 | 2.16E+06 | 1.83E+06 |
| | | | 2.45E+06 | 2.06E+06 | 2.35E+06 | 1.11E+06 | 2.04E+06 | 1.96E+06 | 2.21E+06 | 1.64E+06 |
| Ras/mTOR | *Tor* | Tube 1 | 1.91E+06 | 1.95E+06 | 1.04E+06 | 3.60E+06 | 2.01E+06 | 1.90E+06 | 2.73E+06 | 4.20E+06 |
| | | | 1.79E+06 | 1.60E+06 | 1.10E+06 | 3.87E+06 | 1.88E+06 | 1.60E+06 | 1.96E+06 | 2.56E+06 |
| | | | 1.67E+06 | 1.85E+06 | 2.22E+06 | 4.78E+06 | 1.57E+06 | 2.09E+06 | 3.01E+06 | 3.08E+06 |
| | | Tube 2 | 1.56E+06 | 1.98E+06 | 1.33E+06 | 4.41E+06 | 1.99E+06 | 1.79E+06 | 1.72E+06 | 5.54E+06 |
| | | | 2.07E+06 | 2.10E+06 | 1.25E+06 | 3.85E+06 | 1.93E+06 | 2.46E+06 | 1.61E+06 | 2.74E+06 |
| | | | 1.57E+06 | 1.52E+06 | 1.16E+06 | 3.82E+06 | 2.08E+06 | 1.46E+06 | 1.43E+06 | 2.67E+06 |
| | | Tube 3 | 1.45E+06 | 2.16E+06 | 1.89E+06 | 2.95E+06 | 2.55E+06 | 1.99E+06 | 1.95E+06 | 3.65E+06 |
| | | | 1.50E+06 | 2.12E+06 | 2.17E+06 | 3.67E+06 | 1.79E+06 | 2.19E+06 | 2.33E+06 | 4.65E+06 |
| | | | 1.20E+06 | 1.63E+06 | 1.90E+06 | 4.01E+06 | 2.10E+06 | 2.41E+06 | 2.50E+06 | 3.61E+06 |
| Autophagy | *ATG6* | Tube 1 | 2.12E+06 | 2.65E+06 | 2.75E+06 | 6.75E+06 | 3.01E+06 | 2.63E+06 | 2.10E+06 | 5.28E+06 |
| | | | 2.76E+06 | 2.47E+06 | 2.65E+06 | 7.39E+06 | 3.12E+06 | 2.28E+06 | 2.89E+06 | 4.67E+06 |
| | | | 2.89E+06 | 2.69E+06 | 2.75E+06 | 8.55E+06 | 2.97E+06 | 2.52E+06 | 2.37E+05 | 5.22E+06 |
| | | Tube 2 | 2.95E+06 | 2.25E+06 | 3.55E+06 | 7.50E+06 | 3.05E+06 | 3.25E+06 | 4.75E+06 | 8.44E+06 |
| | | | 2.37E+06 | 2.67E+06 | 2.93E+06 | 6.77E+06 | 2.01E+07 | 2.55E+06 | 2.23E+06 | 7.77E+06 |
| | | | 2.81E+06 | 2.05E+06 | 2.97E+06 | 4.69E+06 | 2.61E+06 | 2.75E+06 | 3.47E+06 | 5.09E+06 |
| | | Tube 3 | 2.69E+06 | 2.23E+06 | 2.45E+06 | 5.42E+06 | 2.71E+06 | 3.03E+06 | 2.85E+06 | 3.92E+06 |
| | | | 2.25E+06 | 2.47E+06 | 2.51E+06 | 4.07E+06 | 3.25E+06 | 2.28E+07 | 2.74E+06 | 4.87E+06 |
| | | | 2.64E+07 | 2.05E+06 | 2.75E+06 | 5.06E+06 | 2.54E+07 | 3.05E+06 | 3.72E+06 | 3.96E+06 |

**Table S24. Statistical analyses of *wsp* absolute counts from RNAi crosses with Gene-Switch GAL4 driver**

| Host cellular process | RNAi target | Comparison performed | Replicate # | Outcome: analysis of different combinations Resulting p-value | Significantly different? | Abundance relative to control | Interpreted as a hit? |
|---|---|---|---|---|---|---|---|
| Wnt | *Armadillo* | Expr+DMSO versus Expr+Mif | 1<br>2 | 0.008<br>0.017 | yes<br>yes | lower<br>lower | yes |
| | | Non-expr+Mif versus Expr+Mif | 1<br>2 | < 0.001<br>< 0.001 | yes<br>yes | lower<br>lower | yes |
| | | Non-expr+DMSO versus Expr+Mif | 1<br>2 | 0.033<br>0.002 | yes<br>yes | lower<br>lower | yes |
| Ras/mTOR | *Tor* | Expr+DMSO versus Expr+Mif | 1<br>2 | 0.031<br>0.002 | yes<br>yes | higher<br>higher | yes |
| | | Non-expr+Mif versus Expr+Mif | 1<br>2 | <0.001<br>0.008 | yes<br>yes | higher<br>higher | yes |
| | | Non-expr+DMSO versus Expr+Mif | 1<br>2 | < 0.001<br>0.002 | yes<br>yes | higher<br>higher | yes |
| Autophagy | *ATG6* | Expr+DMSO versus Expr+Mif | 1<br>2 | < 0.001<br>0.015 | yes<br>yes | higher<br>higher | yes |
| | | Non-expr+Mif versus Expr+Mif | 1<br>2 | 0.045<br>0.007 | yes<br>yes | higher<br>higher | yes |
| | | Non-expr+DMSO versus Expr+Mif | 1<br>2 | 0.031<br>0.326 | yes<br>no | higher<br>n/a | no |

* represent whole body comparisons, all technical replicates, n = 9

**Table S25. Involvement of autophagy pathway in the density regulation of various microbes.**

| Species | Disruption tool/method | Effect on density | Reference |
|---|---|---|---|
| *A. phagocytophilum* | Rapamycin | Increase | Niu et al., 2008 |
| *Candidatus Arthromitus* | *atg5* null mutant | Increase | Yang et al., 2018 |
| *Chlamydia cavia* | *tor* RNAi | Decrease | Derre et al., 2007 |
| *Ehrlicia chaffeensis* | *rheb* RNAi | Decrease | Lina et al., 2017 |
| *Listeria monocytogenes* | *tor* RNAi | Decrease | Derre et al., 2007 |
| *Mycobacterium tuberculosis* | Rapamycin | Decrease | Gutierrez et al., 2004 |
| *Pseudomonas aeruginosa* | Rapamycin | Decrease | Yuan et al., 2012 |
| *Rickettsia australis* | Rapamycin | Increase | Bechelli et al., 2019 |
| *Salmonella typhimurium* | *atg5* null mutant | Increase | Birmingham et al., 2006 |
| *Wolbachia spp.* | Rapamycin (*B. malayi*) | Decrease | Voronin et al., 2012 |
| | Rapamycin (*D. mel*) | Decrease | Voronin et al., 2012 |
| | *atg1* RNAi (*D. mel* germline) | Decrease | Deehan et al., 2021 |
| | *atg8* RNAi (*D. mel* germline) | Decrease | Deehan et al., 2021 |
| | Rapamycin (*D. mel* germline) | Increase | Serbus et al., 2015 |
| | *atg6* RNAi (*D. mel* cell culture) | Increase | Grobler et al., 2018 |
| | *atg18b* RNAi (*D. mel* cell culture) | Increase | Grobler et al., 2018 |
| | Rapamycin (*D. mel*) | Increase | this study |
| | *tor* RNAi (*D. mel*) | Increase | this study |
| | *atg6* RNAi (*D. mel*) | Increase | this study |

**Additional File 1. References cited in Table S8**


Agaisse, H., Burrack, L. S., Philips, J. A., Rubin, E. J., Perrimon, N., & Higgins, D. E. (2005). Genome-wide RNAi screen for host factors required for intracellular bacterial infection. *Science (New York, N.Y.)*, *309*(5738), 1248–1251. https://doi.org/10.1126/science.1116008

Alhazmi, A. (2018). Spleen Tyrosine Kinase as a Target Therapy for Pseudomonas aeruginosa Infection. *Journal of Innate Immunity*, *10*(4), 255–263. https://doi.org/10.1159/000489863

Ashida, H., Mimuro, H., Ogawa, M., Kobayashi, T., Sanada, T., Kim, M., & Sasakawa, C. (2011). Cell death and infection: A double-edged sword for host and pathogen survival. *The Journal of Cell Biology*, *195*(6), 931–942. https://doi.org/10.1083/jcb.201108081

Banerjee, R., Anguita, J., & Fikrig, E. (2000). Granulocytic Ehrlichiosis in Mice Deficient in Phagocyte Oxidase or Inducible Nitric Oxide Synthase. *Infection and Immunity*, *68*(7), 4361–4362.

Bechelli, J., Vergara, L., Smalley, C., Buzhdygan, T. P., Bender, S., Zhang, W., Liu, Y., Popov, V. L., Wang, J., Garg, N., Hwang, S., Walker, D. H., & Fang, R. (2019). Atg5 Supports Rickettsia australis Infection in Macrophages In Vitro and In Vivo. *Infection and Immunity*, *87*(1). https://doi.org/10.1128/IAI.00651-18

Birmingham, C. L., Smith, A. C., Bakowski, M. A., Yoshimori, T., & Brumell, J. H. (2006). Autophagy controls Salmonella infection in response to damage to the Salmonella-containing vacuole. *The Journal of Biological Chemistry*, *281*(16), 11374–11383. https://doi.org/10.1074/jbc.M509157200

Carroll, E. E. M., Wang, X., Shaw, D. K., O'Neal, A. J., Chávez, A. S. O., Brown, L. J., Boradia, V. M., Hammond, H. L., & Pedra, J. H. F. (2019). P47 licenses activation of the immune deficiency pathway in the tick Ixodes scapularis. *Proceedings of the National Academy of Sciences*, *116*(1), 205–210. https://doi.org/10.1073/pnas.1808905116

Cole, C., Thomas, S., Filak, H., Henson, P. M., & Lenz, L. L. (2012). Nitric Oxide Increases Susceptibility of Toll-like Receptor-Activated Macrophages to Spreading Listeria monocytogenes. *Immunity*, *36*(5), 807–820. https://doi.org/10.1016/j.immuni.2012.03.011

Colonne, P. M., Eremeeva, M. E., & Sahni, S. K. (2011). Beta Interferon-Mediated Activation of Signal Transducer and Activator of Transcription Protein 1 Interferes with Rickettsia conorii Replication in Human Endothelial Cells. *Infection and Immunity*, *79*(9), 3733–3743. https://doi.org/10.1128/IAI.05008-11

Czyż, D. M., Jain-Gupta, N., Shuman, H. A., & Crosson, S. (2016). A dual-targeting approach to inhibit *Brucella abortus* replication in human cells. *Scientific Reports*, *6*, 35835. https://doi.org/10.1038/srep35835

Czyż, D. M., Potluri, L.-P., Jain-Gupta, N., Riley, S. P., Martinez, J. J., Steck, T. L., Crosson, S., Shuman, H. A., & Gabay, J. E. (2014). Host-Directed Antimicrobial Drugs with Broad-Spectrum Efficacy against Intracellular Bacterial Pathogens. *MBio*, *5*(4). https://doi.org/10.1128/mBio.01534-14

Derré, I., Pypaert, M., Dautry-Varsat, A., & Agaisse, H. (2007). RNAi Screen in Drosophila Cells Reveals the Involvement of the Tom Complex in Chlamydia Infection. *PLoS Pathogens*, *3*(10). https://doi.org/10.1371/journal.ppat.0030155

Ferree, P. M., Frydman, H. M., Li, J. M., Cao, J., Wieschaus, E., & Sullivan, W. (2005). Wolbachia Utilizes Host Microtubules and Dynein for Anterior Localization in the

Drosophila Oocyte. *PLoS Pathogens*, *1*(2). https://doi.org/10.1371/journal.ppat.0010014

Foldenauer, M. E. B., McClellan, S. A., Berger, E. A., & Hazlett, L. D. (2013). Mammalian Target of Rapamycin Regulates IL-10 and Resistance to Pseudomonas aeruginosa Corneal Infection. *The Journal of Immunology*, *190*(11), 5649–5658. https://doi.org/10.4049/jimmunol.1203094

Grobler, Y., Yun, C. Y., Kahler, D. J., Bergman, C. M., Lee, H., Oliver, B., & Lehmann, R. (2018). Whole genome screen reveals a novel relationship between Wolbachia levels and Drosophila host translation. *PLoS Pathogens*, *14*(11), e1007445. https://doi.org/10.1371/journal.ppat.1007445

Gross, A., Terraza, A., Ouahrani-Bettache, S., Liautard, J.-P., & Dornand, J. (2000). In Vitro Brucella suis Infection Prevents the Programmed Cell Death of Human Monocytic Cells. *Infection and Immunity*, *68*(1), 342–351.

Gutierrez, M. G., Master, S. S., Singh, S. B., Taylor, G. A., Colombo, M. I., & Deretic, V. (2004). Autophagy is a defense mechanism inhibiting BCG and Mycobacterium tuberculosis survival in infected macrophages. *Cell*, *119*(6), 753–766. https://doi.org/10.1016/j.cell.2004.11.038

Herren, J. K., & Lemaitre, B. (2011). Spiroplasma and host immunity: Activation of humoral immune responses increases endosymbiont load and susceptibility to certain Gram-negative bacterial pathogens in Drosophila melanogaster. *Cellular Microbiology*, *13*(9), 1385–1396. https://doi.org/10.1111/j.1462-5822.2011.01627.x

Jiménez de Bagüés, M. P., Gross, A., Terraza, A., & Dornand, J. (2005). Regulation of the Mitogen-Activated Protein Kinases by Brucella spp. Expressing a Smooth and Rough Phenotype: Relationship to Pathogen Invasiveness. *Infection and Immunity*, *73*(5), 3178–3183. https://doi.org/10.1128/IAI.73.5.3178-3183.2005

Kessler, M., Zielecki, J., Thieck, O., Mollenkopf, H.-J., Fotopoulou, C., & Meyer, T. F. (2012). Chlamydia trachomatis disturbs epithelial tissue homeostasis in fallopian tubes via paracrine Wnt signaling. *The American Journal of Pathology*, *180*(1), 186–198. https://doi.org/10.1016/j.ajpath.2011.09.015

Kim, D. H., Lim, J. J., Lee, J. J., Kim, D. G., Lee, H. J., Min, W., Kim, K. D., Chang, H. H., Rhee, M. H., Watarai, M., & Kim, S. (2012). Identification of genes contributing to the intracellular replication of Brucella abortus within HeLa and RAW 264.7 cells. *Veterinary Microbiology*, *158*(3–4), 322–328. https://doi.org/10.1016/j.vetmic.2012.02.019

Kintner, J., Moore, C. G., Whittimore, J. D., Butler, M., & Hall, J. V. (2017). Inhibition of Wnt Signaling Pathways Impairs Chlamydia trachomatis Infection in Endometrial Epithelial Cells. *Frontiers in Cellular and Infection Microbiology*, *7*, 501. https://doi.org/10.3389/fcimb.2017.00501

Kühbacher, A., Emmenlauer, M., Rämo, P., Kafai, N., Dehio, C., Cossart, P., & Pizarro-Cerdá, J. (2015). Genome-Wide siRNA Screen Identifies Complementary Signaling Pathways Involved in Listeria Infection and Reveals Different Actin Nucleation Mechanisms during Listeria Cell Invasion and Actin Comet Tail Formation. *MBio*, *6*(3), e00598-00515. https://doi.org/10.1128/mBio.00598-15

Kuijl, C., Savage, N. D. L., Marsman, M., Tuin, A. W., Janssen, L., Egan, D. A., Ketema, M., van den Nieuwendijk, R., van den Eeden, S. J. F., Geluk, A., Poot, A., van der Marel, G., Beijersbergen, R. L., Overkleeft, H., Ottenhoff, T. H. M., & Neefjes, J. (2007). Intracellular bacterial growth is controlled by a kinase network around PKB/AKT1. *Nature*, *450*(7170), 725–730. https://doi.org/10.1038/nature06345

Lad, S. P., Fukuda, E. Y., Li, J., Maza, L. M. de la, & Li, E. (2005). Up-Regulation of the JAK/STAT1 Signal Pathway during Chlamydia trachomatis Infection. *The Journal of Immunology*, *174*(11), 7186–7193. https://doi.org/10.4049/jimmunol.174.11.7186

Levenhagen, M. A., Alves, R. N., Rieck, S. E., Labruna, M. B., & Beletti, M. E. (2012). The role of cytoskeleton, components of inositol phospholipid signaling pathway and iron in <Emphasis Type="Italic">Ehrlichia canis</Emphasis> in vitro proliferation. *Veterinary Research Communications*, *36*(3), 195–199. https://doi.org/10.1007/s11259-012-9525-y

Limmer, S., Haller, S., Drenkard, E., Lee, J., Yu, S., Kocks, C., Ausubel, F. M., & Ferrandon, D. (2011). Pseudomonas aeruginosa RhlR is required to neutralize the cellular immune response in a Drosophila melanogaster oral infection model. *Proceedings of the National Academy of Sciences*, *108*(42), 17378–17383. https://doi.org/10.1073/pnas.1114907108

Lin, M., Zhu, M. X., & Rikihisa, Y. (2002). Rapid Activation of Protein Tyrosine Kinase and Phospholipase C-γ2 and Increase in Cytosolic Free Calcium Are Required by Ehrlichia chaffeensis for Internalization and Growth in THP-1 Cells. *Infection and Immunity*, *70*(2), 889–898. https://doi.org/10.1128/IAI.70.2.889-898.2002

Lina, T. T., Luo, T., Velayutham, T.-S., Das, S., & McBride, J. W. (2017). Ehrlichia Activation of Wnt-PI3K-mTOR Signaling Inhibits Autolysosome Generation and Autophagic Destruction by the Mononuclear Phagocyte. *Infection and Immunity*, *85*(12). https://doi.org/10.1128/IAI.00690-17

Liu, L., Dai, J., Zhao, Y. O., Narasimhan, S., Yang, Y., Zhang, L., & Fikrig, E. (2012). Ixodes scapularis JAK-STAT pathway regulates tick antimicrobial peptides, thereby controlling the agent of human granulocytic anaplasmosis. *The Journal of Infectious Diseases*, *206*(8), 1233–1241. https://doi.org/10.1093/infdis/jis484

Luo, T., Dunphy, P. S., Lina, T. T., & McBride, J. W. (2016). Ehrlichia chaffeensis Exploits Canonical and Noncanonical Host Wnt Signaling Pathways To Stimulate Phagocytosis and Promote Intracellular Survival. *Infection and Immunity*, *84*(3), 686–700. https://doi.org/10.1128/IAI.01289-15

Luo, T., Dunphy, P. S., & McBride, J. W. (2017). Ehrlichia chaffeensis Tandem Repeat Effector Targets Differentially Influence Infection. *Frontiers in Cellular and Infection Microbiology*, *7*. https://doi.org/10.3389/fcimb.2017.00178

Newton, I. L. G., & Sheehan, K. B. (2015). Passage of Wolbachia pipientis through Mutant Drosophila melanogaster Induces Phenotypic and Genomic Changes. *Applied and Environmental Microbiology*, *81*(3), 1032–1037. https://doi.org/10.1128/AEM.02987-14

Niu, H., Xiong, Q., Yamamoto, A., Hayashi-Nishino, M., & Rikihisa, Y. (2012). Autophagosomes induced by a bacterial Beclin 1 binding protein facilitate obligatory intracellular infection. *Proceedings of the National Academy of Sciences*, *109*(51), 20800–20807. https://doi.org/10.1073/pnas.1218674109

Niu, H., Yamaguchi, M., & Rikihisa, Y. (2008). Subversion of cellular autophagy by Anaplasma phagocytophilum. *Cellular Microbiology*, *10*(3), 593–605. https://doi.org/10.1111/j.1462-5822.2007.01068.x

Philips, J. A., Rubin, E. J., & Perrimon, N. (2005). Drosophila RNAi screen reveals CD36 family member required for mycobacterial infection. *Science (New York, N.Y.)*, *309*(5738), 1251–1253. https://doi.org/10.1126/science.1116006

Rennoll, S. A., Rennoll-Bankert, K. E., Guillotte, M. L., Lehman, S. S., Driscoll, T. P., Beier-Sexton, M., Rahman, M. S., Gillespie, J. J., & Azad, A. F. (2017). The Cat

Flea (Ctenocephalides felis) Immune Deficiency Signaling Pathway Regulates Rickettsia typhi Infection. *Infection and Immunity*, *86*(1), e00562-17. https://doi.org/10.1128/IAI.00562-17

Rikihisa, Y., Zhang, Y., & Park, J. (1994). Inhibition of infection of macrophages with Ehrlichia risticii by cytochalasins, monodansylcadaverine, and taxol. *Infection and Immunity*, *62*(11), 5126–5132.

Rikihisa, Y., Zhang, Y., & Park, J. (1995). Role of Ca2+ and calmodulin in ehrlichial infection in macrophages. *Infection and Immunity*, *63*(6), 2310–2316.

Rogan, M. R., Patterson, L. L., Wang, J. Y., & McBride, J. W. (2019). Bacterial Manipulation of Wnt Signaling: A Host-Pathogen Tug-of-Wnt. *Frontiers in Immunology*, *10*. https://doi.org/10.3389/fimmu.2019.02390

Serbus, L. R., Ferreccio, A., Zhukova, M., McMorris, C. L., Kiseleva, E., & Sullivan, W. (2011). A feedback loop between Wolbachia and the Drosophila gurken mRNP complex influences Wolbachia titer. *Journal of Cell Science*, *124*(Pt 24), 4299–4308. https://doi.org/10.1242/jcs.092510

Severo, M. S., Choy, A., Stephens, K. D., Sakhon, O. S., Chen, G., Chung, D.-W. D., Le Roch, K. G., Blaha, G., & Pedra, J. H. F. (2013). The E3 ubiquitin ligase XIAP restricts Anaplasma phagocytophilum colonization of Ixodes scapularis ticks. *The Journal of Infectious Diseases*, *208*(11), 1830–1840. https://doi.org/10.1093/infdis/jit380

Steele, S., Brunton, J., Ziehr, B., Taft-Benz, S., Moorman, N., & Kawula, T. (2013). Francisella tularensis Harvests Nutrients Derived via ATG5-Independent Autophagy to Support Intracellular Growth. *PLOS Pathogens*, *9*(8), e1003562. https://doi.org/10.1371/journal.ppat.1003562

Turco, J., Liu, H., Gottlieb, S. F., & Winkler, H. H. (1998). Nitric oxide-mediated inhibition of the ability of Rickettsia prowazekii to infect mouse fibroblasts and mouse macrophagelike cells. *Infection and Immunity*, *66*(2), 558–566.

Walker, D. H., Firth, W. T., Ballard, J. G., & Hegarty, B. C. (1983). Role of Phospholipase-Associated Penetration Mechanism in Cell Injury by Rickettsia rickettsii. *Infection and Immunity*, *40*(2), 840–842.

Wang, J., Wu, Y., Yang, G., & Aksoy, S. (2009). Interactions between mutualist Wigglesworthia and tsetse peptidoglycan recognition protein (PGRP-LB) influence trypanosome transmission. *Proceedings of the National Academy of Sciences*, *106*(29), 12133–12138. https://doi.org/10.1073/pnas.0901226106

Wang, J., Yang, K., Zhou, L., MinhaoWu, Wu, Y., Zhu, M., Lai, X., Chen, T., Feng, L., Li, M., Huang, C., Zhong, Q., & Huang, X. (2013). MicroRNA-155 Promotes Autophagy to Eliminate Intracellular Mycobacteria by Targeting Rheb. *PLOS Pathogens*, *9*(10), e1003697. https://doi.org/10.1371/journal.ppat.1003697

Wei, P., Cui, G., Lu, Q., Yang, L., Guan, Z., Sun, W., Zhao, Y., Wang, S., & Peng, Q. (2015). A20 promotes Brucella intracellular growth via inhibition of macrophage cell death and activation. *Veterinary Microbiology*, *175*(1), 50–57. https://doi.org/10.1016/j.vetmic.2014.11.006

White, P. M., Serbus, L. R., Debec, A., Codina, A., Bray, W., Guichet, A., Lokey, R. S., & Sullivan, W. (2017). Reliance of Wolbachia on High Rates of Host Proteolysis Revealed by a Genome-Wide RNAi Screen of Drosophila Cells. *Genetics*, *205*(4), 1473–1488. https://doi.org/10.1534/genetics.116.198903

Xiong, Q., Lin, M., & Rikihisa, Y. (2009). Cholesterol-Dependent Anaplasma phagocytophilum Exploits the Low-Density Lipoprotein Uptake Pathway. *PLoS Pathogens*, *5*(3). https://doi.org/10.1371/journal.ppat.1000329

Yang, L., Liu, C., Zhao, W., He, C., Ding, J., Dai, R., Xu, K., Xiao, L., Luo, L., Liu, S., Li, W., & Meng, H. (2018). Impaired Autophagy in Intestinal Epithelial Cells Alters Gut Microbiota and Host Immune Responses. *Applied and Environmental Microbiology*, *84*(18). https://doi.org/10.1128/AEM.00880-18

Yuan, K., Huang, C., Fox, J., Laturnus, D., Carlson, E., Zhang, B., Yin, Q., Gao, H., & Wu, M. (2012). Autophagy plays an essential role in the clearance of Pseudomonas aeruginosa by alveolar macrophages. *Journal of Cell Science*, *125*(Pt 2), 507–515. https://doi.org/10.1242/jcs.094573

Zhang, Q., Wang, C., Liu, Z., Liu, X., Han, C., Cao, X., & Li, N. (2012). Notch signal suppresses Toll-like receptor-triggered inflammatory responses in macrophages by inhibiting extracellular signal-regulated kinase 1/2-mediated nuclear factor κB activation. *The Journal of Biological Chemistry*, *287*(9), 6208–6217. https://doi.org/10.1074/jbc.M111.310375

Zhang, Y., & Rikihisa, Y. (1997). Tyrosine phosphorylation is required for ehrlichial internalization and replication in P388D1 cells. *Infection and Immunity*, *65*(7), 2959–2964.

Zhu, B., Das, S., Mitra, S., Farris, T. R., & McBride, J. W. (2017). Ehrlichia chaffeensis TRP120 Moonlights as a HECT E3 Ligase Involved in Self- and Host Ubiquitination To Influence Protein Interactions and Stability for Intracellular Survival. *Infection and Immunity*, *85*(9). https://doi.org/10.1128/IAI.00290-17
